# Accelerated diversification correlated with functional traits shapes extant diversity of the early divergent angiosperm family Annonaceae

**DOI:** 10.1101/652065

**Authors:** B. Xue, X. Guo, J.B. Landis, M. Sun, C.C. Tang, P.S. Soltis, D.E. Soltis, R.M.K. Saunders

## Abstract

**Background:** A major goal of phylogenetic systematics is to understand both the patterns of diversification and the processes by which these patterns are formed. Few studies have focused on the ancient, species-rich Magnoliales clade and its diversification pattern. Within Magnoliales, the pantropically distributed Annonaceae are by far the most genus-rich and species-rich family-level clade, with *c.* 110 genera and *c*. 2,400 species. We investigated the diversification patterns across Annonaceae and identified traits that show varied associations with diversification rates using a time-calibrated phylogeny of 835 species (34.6% sampling) and 11,211 aligned bases from eight regions of the plastid genome (*rbcL*, *matK*, *ndhF*, *psbA-trnH*, *trnL-F*, *atpB-rbcL*, *trnS-G*, and *ycf1*). Two hypotheses that might explain patterns of diversification—the ‘museum model’ and heterogeneous diversification rates—are also evaluated.

**Results:** Twelve rate shifts were identified using BAMM: in *Annona*, *Artabotrys*, *Asimina*, *Drepananthus*, *Duguetia*, *Goniothalamus*, *Guatteria*, *Uvaria*, *Xylopia*, the tribes Miliuseae and Malmeeae, and the *Desmos*-*Dasymaschalon*-*Friesodielsia*-*Monanthotaxis* clade (which collectively account for over 80% of the total species richness in the family). TurboMEDUSA and method-of-moments estimator analyses showed largely congruent results. A positive relationship between species richness and diversification rate is revealed using PGLS. We further explore the possible role of selected traits (habit, pollinator trapping, floral sex expression, pollen dispersal unit, anther septation, and seed dispersal unit) in shaping diversification patterns, based on inferences of BiSSE, MuSSE, HiSSE, and FiSSE analyses. Our results suggest that the liana habit, the presence of circadian pollinator trapping, androdioecy, and the dispersal of seeds as single-seeded monocarp fragments are closely correlated with higher diversification rates; pollen aggregation and anther septation, in contrast, are associated with lower diversification rates.

**Conclusion:** Our results show that the high species richness in Annonaceae is likely the result of recent increased diversification rather than the steady accumulation of species via the ‘museum model’. BAMM, turboMEDUSA, and the method-of-moments estimator all indicate heterogeneity in diversification rates across the phylogeny, with different traits associated with shifts in diversification rates in different Annonaceae clades.

## Background

The asymmetry of species richness is a conspicuous ecological and evolutionary phenomenon in the Tree of Life. Differences in species richness among clades have been alternatively explained by the ‘museum model’ and diversification rate hypotheses [1,2]: according to the former hypothesis, species diversity is associated with clade age, with older clades having a longer time for richness to accumulate, whereas the diversification rate hypothesis suggests that clades can undergo speciation at different rates, with high net diversification rates generating greater species diversity irrespective of clade age. Accelerated diversification might be linked to changes in a diversity of traits, such as genomic features, ecology, and/or morphology ([2], and references therein).

Although the likely drivers of rapid species radiation have been widely investigated in lineages of eudicots (*e.g.*, Ericaceae: [3]; Campanulaceae: [4]; Proteaceae: [5]; Plantaginaceae: [6]) and monocots (*e.g.*, Bromeliaceae: [7]; Arecaceae: [8]; Orchidaceae: [9]), diversification pattern in more ancient angiosperm clades are poorly understood. Within the magnoliid order Magnoliales, for example, the species-rich family Annonaceae [10] provides a stark contrast with its sister family, Eupomatiaceae, which comprises only three species [11]. Annonaceae are by far the most genus-rich and species-rich family-level clade in Magnoliales, with *c.* 110 genera and *c*. 2,400 species [10,12,13]. Four subfamilies are recognized in Annonaceae: Anaxagoreoideae (30 spp.) and Ambavioideae (56 spp.) are the subsequent sisters to the rest of the family, while most species are placed in two larger subfamilies, Annonoideae (1,515 spp.) and Malmeoideae (783 spp.) [10,12,13]. The family is further subdivided into 15 tribes [10,13]. Annonaceae have a pantropical distribution and are an important ecological component of many lowland tropical forest ecosystems [14,15]. The family is also well-studied taxonomically and phylogenetically. Hence, Annonaceae is an ideal model system for studying the causes of asymmetry in species richness, providing an additional case outside monocots and eudicots.

Two previous studies focused on diversification patterns within Annonaceae. Couvreur *et al.* [15] investigated diversification rates in the family by applying lineage-through-time plots and a maximum likelihood approach based on a 100-taxon phylogeny (83% generic sampling). They demonstrated that the family as a whole has undergone relatively constant, slow diversification with low extinction rates, although this may be an artefact of taxon sampling limitations [16]. Erkens *et al.* [17] subsequently used topological and temporal methods of diversification analysis to identify clades that were likely to have undergone radiation, based on one of the 200-taxon trees (83% generic sampling) generated by Chatrou *et al.* [12]. The topological methods identified three shifts to increased rates: in the Annoneae-Monodoreae-Uvarieae and Miliuseae-Monocarpieae-Malmeeae clades and the tribe Miliuseae. The temporal methods revealed significant rate increases in *Goniothalamus*, *Isolona*, *Monodora*, and *Stenanona*. These studies provide an initial understanding of diversification patterns within the family; nevertheless, taxon sampling was limited, with less than 10% of recognized species sampled, and did not attempt to identify traits that are associated with these rates shifts (and therefore potentially causative), although Erkens *et al.* [17] provided a narrative assessment of possible relationships between character evolution and shifts in diversification rate.

The past decade has seen significant advances in our knowledge of Annonaceae phylogeny (*e.g.*, [10,12,18,19]). DNA sequences of nearly 1,000 species have been generated, enabling a good understanding of phylogenetic relationships and generic delimitation in the family. Considerable progress has also been made in understanding the distribution of morphological characters across the family and their possible roles in diversification, especially floral and fruit traits of ecological importance such as floral chambers [20], the role of stigmatic exudate as an extragynoecial compitum [21], staminate/pistillate-phase floral synchrony [22], the circadian trapping of pollinators [23,24], and seed dispersal units (discussed below; [25]). These advances in both molecular and morphological data accumulation and new analytical methods provide excellent opportunities to use recent comprehensive phylogenies to date divergence times in the family and to study diversification patterns in relation to key functional traits.

We constructed a phylogeny of 835 Annonaceae species, representing the most complete species- and genus-level coverage to date (34.6% of species, 98% of genera) and performed lineage diversification analyses based on this phylogeny using multiple methods, including Bayesian Analysis of Macroevolutionary Mixtures (BAMM), Modeling Evolutionary Diversification Using Stepwise Akaike Information Criterion (turboMEDUSA), and method-of-moments estimator, to assess patterns of rate variation in Annonaceae. We also performed phylogeny-based trait evolution and trait-dependent diversification analyses to test the influence of morphology on species diversification patterns across Annonaceae. We specifically address the following questions: (1) Is species richness in Annonaceae better explained by the clade age or diversification rate hypothesis? (2) Are significant diversification rate shifts evident in Annonaceae? (3) Are some traits strongly correlated with shifts in diversification rates, and what are the possible explanations?

## Results

### Phylogenetic analyses, divergence time estimation, and ancestral character reconstruction

The resultant phylogeny for 923 taxa (Supplementary Fig. S1) is congruent with previous family-level studies [10,12], although there is some incongruence in the position of some genera between plastid-based studies and nuclear gene-based phylogenies [26]. This is the largest phylogeny of Annonaceae to date, and the subfamilies and tribes are all well supported, although the backbone of tribe Miliuseae is not resolved as similarly reported in previous studies [10,12,26]. The dating results (Supplementary Fig. S2) are similar to previous studies: the ages of major tribes, for example, are similar to those estimated by Thomas *et al.* [19] using BEAST based on the same fossil calibrations. The pruned phylogeny (excluding the outgroup, duplicate taxa, subspecies, undescribed or unidentified species) for our diversification analysis consisted of 835 species (Supplementary Fig. S3). The most likely ancestral character states using stochastic character mapping SIMMAP for the ancestral node of Annonaceae were reconstructed as: self-supporting trees/shrubs, lacking a pollination trap, aseptate anthers, and pollen dispersed as monads. Ancestral states for population-level floral sex expression were ambiguous (Supplementary Figure S4).

### Trait-independent diversification

We applied a range of methods to analyze diversification patterns in Annonaceae. Selected methods were of two types: trait-independent and trait-dependent. Three trait-independent diversification methods were employed to detect diversification rate shifts: modelling evolutionary diversification using stepwise AIC (turboMEDUSA; [27]), Bayesian analysis of macroevolutionary mixtures (BAMM; [28]), method-of-moments estimator [29]. All three methods indicated heterogeneity in diversification rates across the phylogeny.

After discarding 10% burn-in, we confirmed convergence of the MCMC chains in the BAMM analyses; ESS values were 1093.803 and 1166.003 for the log likelihoods and number of rate shifts, respectively. BAMM identified 12 clades with increased diversification rates: *Annona*, *Artabotrys*, *Asimina*, *Drepananthus*, *Duguetia*, *Goniothalamus*, *Guatteria*, *Uvaria*, and *Xylopia*; the tribes Miliuseae and Malmeeae; and the *Desmos*-*Dasymaschalon*-*Friesodielsia*-*Monanthotaxis* clade (Fig. 1). The Bayes factors are all greater than 13.

**Fig. 1.**
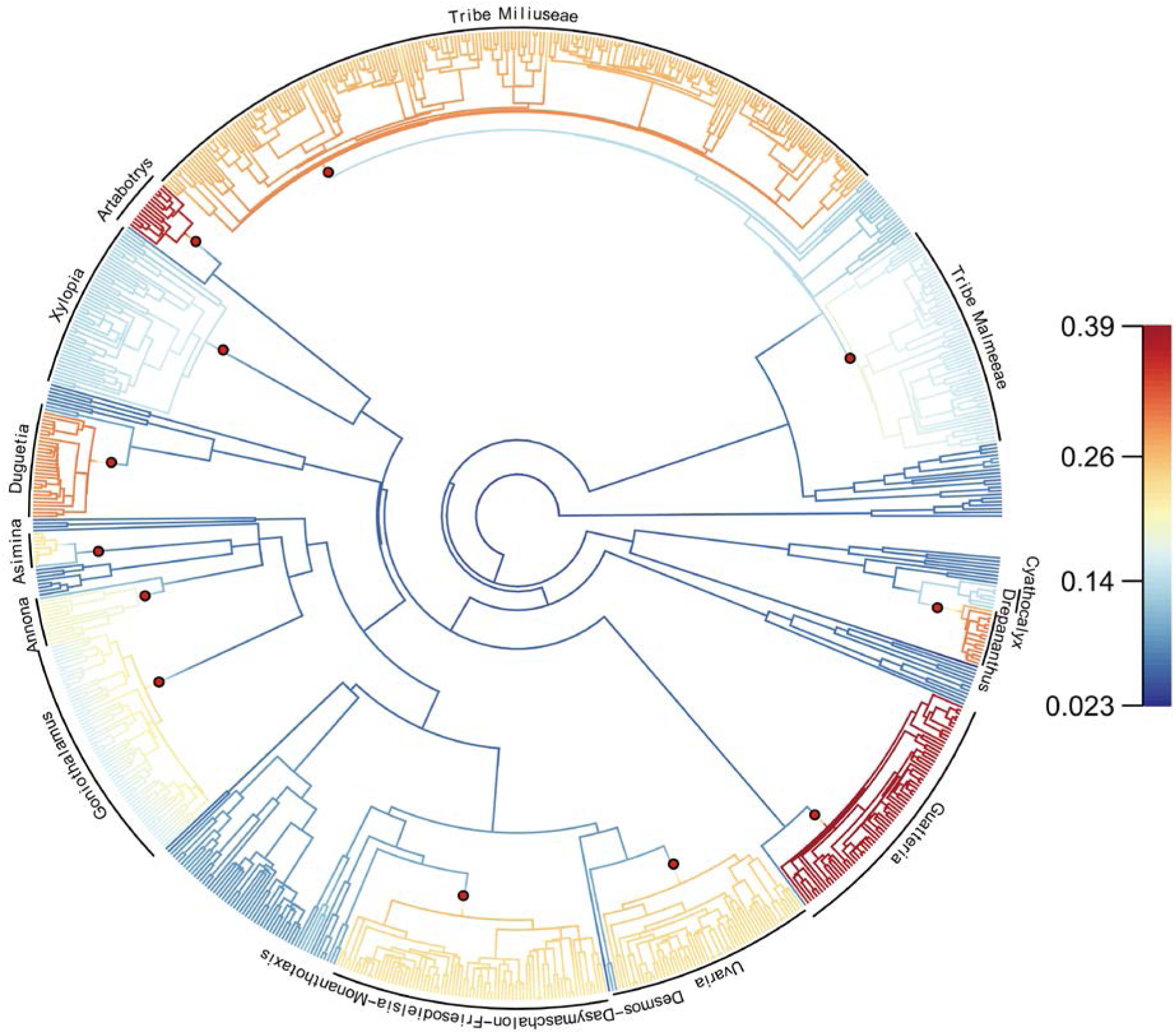
Phylorate plot of Annonaceae with branches colored according to net diversification rate (Myr^−1^), resulting from BAMM analysis. Red dots indicate diversification rate shifts.

In the turboMEDUSA analysis, seven diversification rate shifts were identified across Annonaceae (Fig. 2). The background net diversification rate (r) for the family was estimated as 0.0594 Myr^−1^. A rate increase occurred in the clade comprising the tribes Malmeeae, Miliuseae and five small tribes (Maasieae, Fenerivieae, Phoenicantheae, Dendrokingstonieae, and Monocarpieae) (Clade 1, Fig. 2), with a subsequent decrease in the five small tribes (Clade 6, Fig. 2), and further significant increase in tribe Miliuseae (Clade 5, Fig. 2). A rate increase was also found in tribe Uvarieae (Clade 2, Fig. 2), with a subsequent decrease in *Dielsiothamnus* (Clade 4, Fig. 2), and a further significant increase in the *Desmos*-*Dasymaschalon*-*Friesodielsia*-*Monanthotaxis* clade (Clade 7, Fig. 2). A diversification rate decrease was detected in the monotypic genus *Meiocarpidium* (Clade 3, Fig. 2). Rate shifts in Clades 1, 2, 5, and 7 were also revealed in the BAMM analysis.

**Fig. 2.**
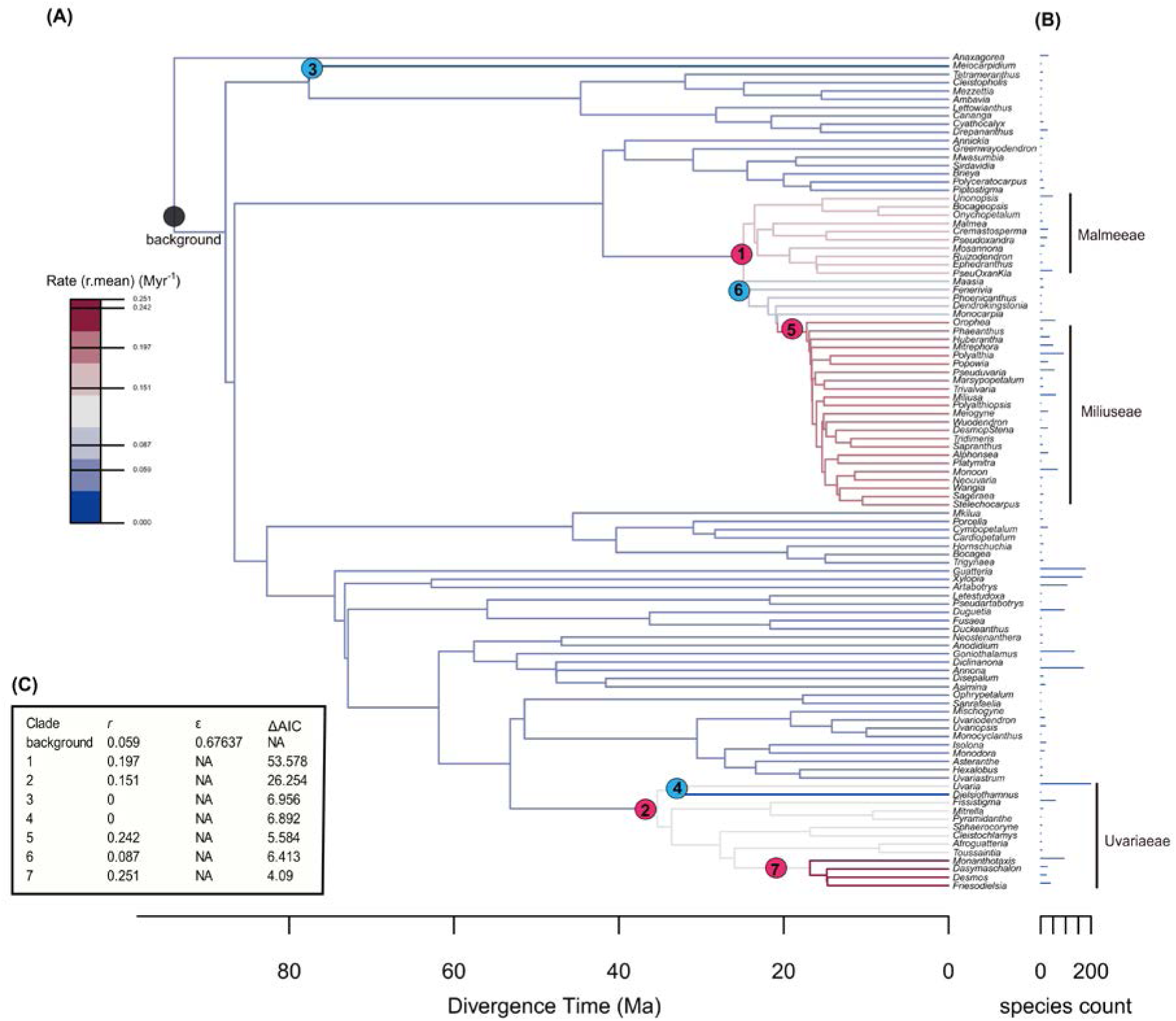
Diversity tree for turboMEDUSA analyses of lineage diversification in Annonaceae. (**A**) The time tree was collapsed to 101 representative stem lineages and colored according to estimated diversification rates. Clades with unusual diversification rates are denoted with numbers that indicate the order in which rate shifts were added by the stepwise Akaike information criterion (AIC) procedure. (**B**) Estimated extant species numbers. (**C**) Estimated net diversification rate (*r* = λ−μ) and relative extinction rate (ε = μ/λ). Red and blue circles represent clades with rate increases and decreases, respectively, compared with the background rate shown as the gray circle. NA, Not applicable, when a yule model was fit to the data, the extinction rate does not exist, so the relative extinction rate is not estimated; Ma, million years ago.

Estimated rates from the method-of-moments estimator are provided in Supplementary Table S1, along with species richness and ages of clades. We found a largely similar pattern of diversification rates as calculated by BAMM. Among the 12 clades with rate shifts inferred from BAMM, four were also identified by turboMEDUSA, eight were supported by the stem-group estimator, and all were supported by the crown-group estimator (Supplementary Table S1). The BAMM analyses were therefore generally concordant with the turboMEDUSA and method-of-moments estimator results.

### Rate-through-time plots

Net diversification rates for the family, as reconstructed by BAMM, indicate a slow increase until around 25 Ma, after which a significant increase is evident until the present (Fig. 3A). Plotting rate through time for the four subfamilies suggested different diversification patterns for each (Fig. 3B). The small subfamily Anaxagoreoideae had a low and constant diversification rate. Subfamily Ambavioideae experienced a more recent rate acceleration during the last 10 Ma. Both subfamilies Annonoideae and Malmeoideae began to diversify rapidly around 25 Ma (Fig. 3B).

**Fig. 3.**
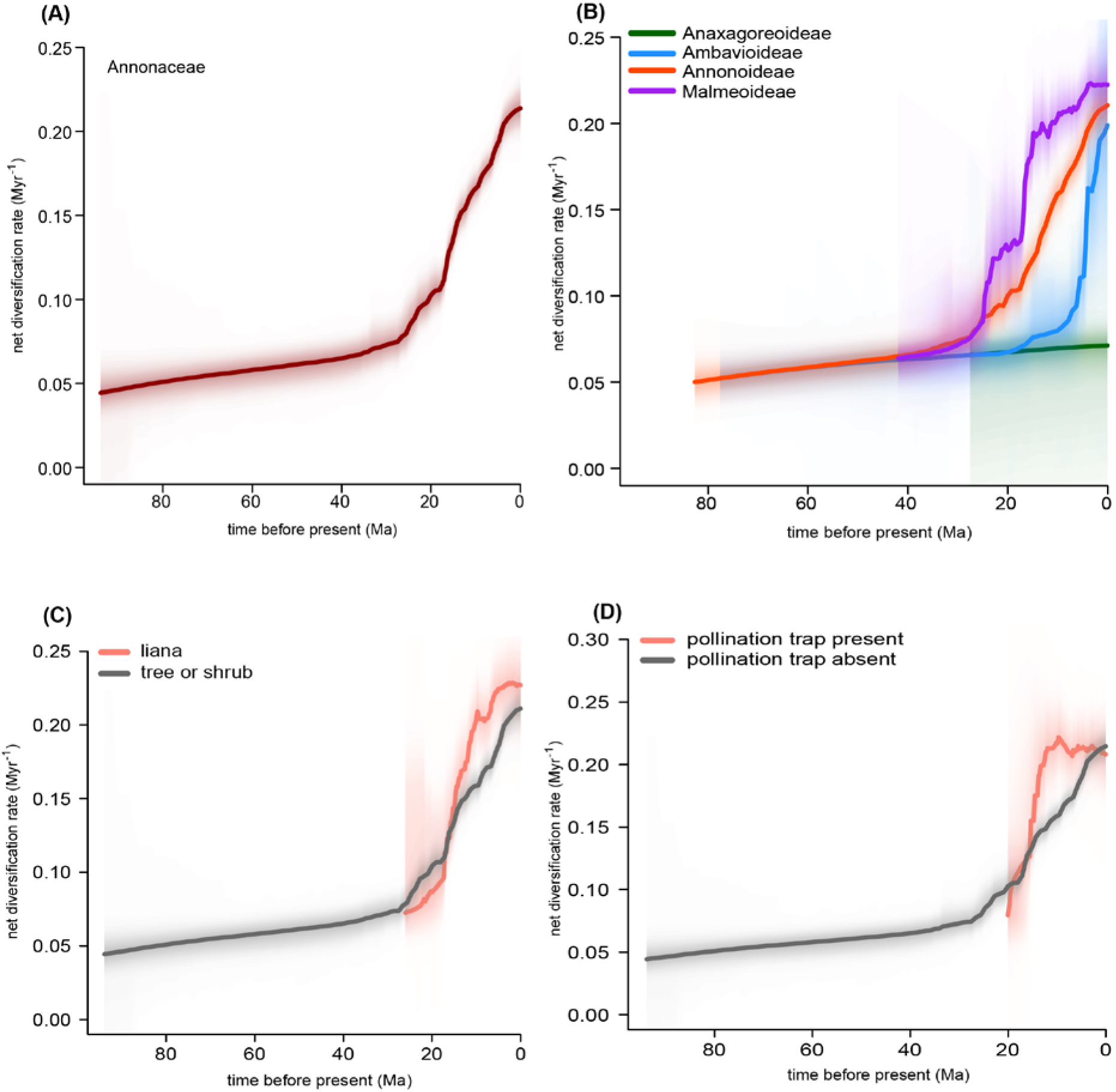
Net diversification rate through time plots from BAMM analysis for Annonaceae (**A**), the four subfamilies (**B**), the liana and tree/shrub lineages (**C**), and lineages with and without a pollination trap (**D**). The shaded areas indicate the 95% confidence interval.

Plotting rate through time for lineages associated with different traits indicated that liana-dominated lineages generally have higher net diversification rates than lineages of trees or shrubs (Fig. 3C). Similarly, lineages with pollinator trapping generally exhibit a higher net diversification rate than those without a trap mechanism (Fig. 3D). A consistent pattern of rate acceleration around 25–20 Ma is observed from all of the rate-through-time plots, except in subfamilies Anaxagoreoideae and Ambavioideae (Fig. 3).

### Museum model *vs* diversification rate hypotheses

There was a weak correlation between species richness and stem age (*r^2^* = 0.058, *p* = 0.013, Fig. 4A). In contrast, species richness is strongly positively correlated with diversification rate under an intermediate ε of 0.45 (*r^2^* = 0.825, *p* < 0.001, Fig. 4B). The patterns hold when testing for diversification rates under low and high values of ε (ε = 0 and 0.9) and are similar (*r^2^* = 0.804, 0.827, respectively, *p* < 0.001). We therefore only present results from the intermediate value (ε = 0.45, Fig. 4B).

**Fig. 4.**
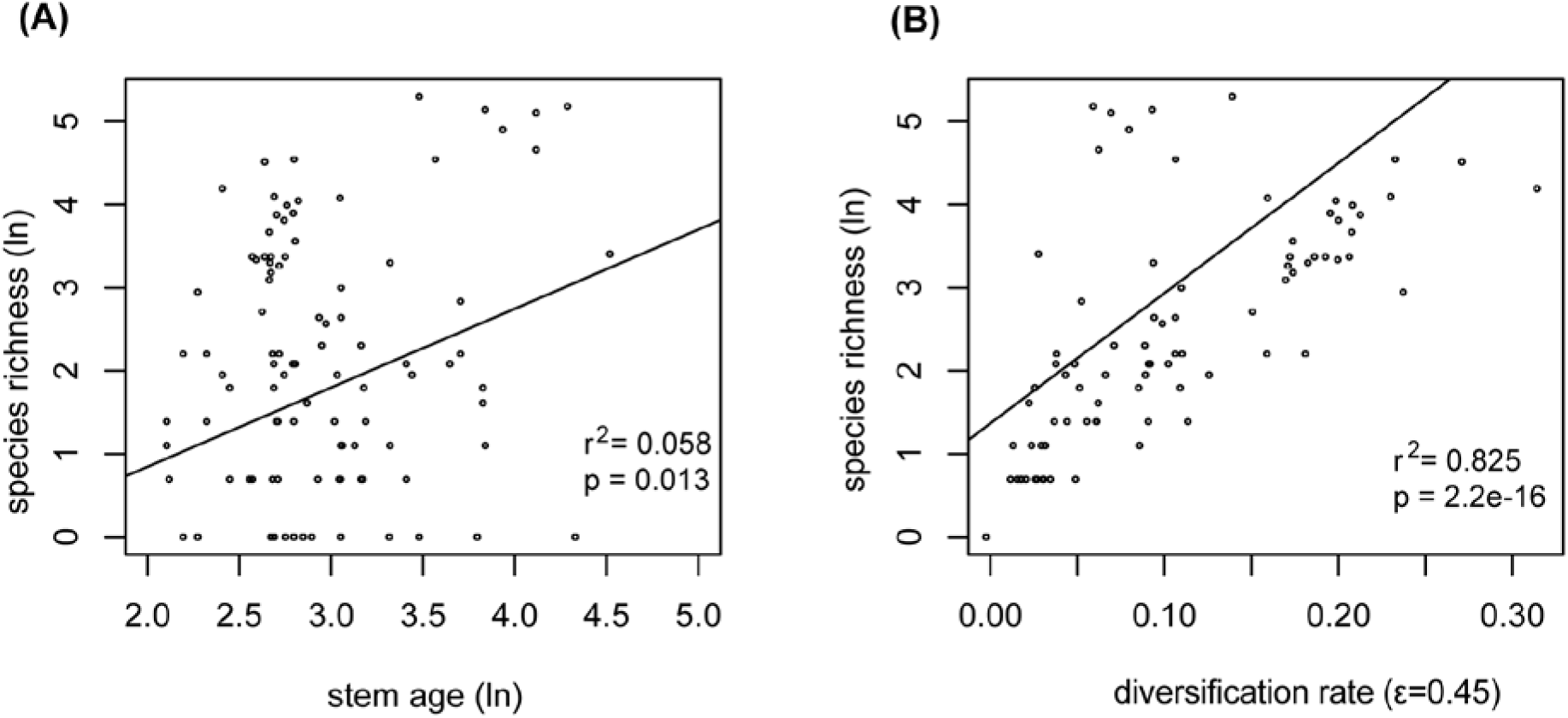
PGLS analyses of the relationship between clade species richness and stem age/diversification rate. (**A**) Clade species richness (ln-transformed) versus stem age (ln-transformed); (**B**) clade species richness (ln-transformed) versus diversification rate (ε=0.45).

### Trait-dependent diversification

Four methods to detect trait-dependent diversification rates were applied: binary-state speciation and extinction (BiSSE; [30]); the multiple state speciation and extinction (MuSSE; [31]); fast, intuitive state-dependent speciation and extinction (FiSSE; [32]); and hidden state speciation and extinction (HiSSE; [33]). The best-fit BiSSE, HiSSE and MuSSE models for different traits are summarized in Supplementary Tables S2–S4, with the rate comparison results summarized in Table 1 and Fig. 5.

**Fig. 5.**
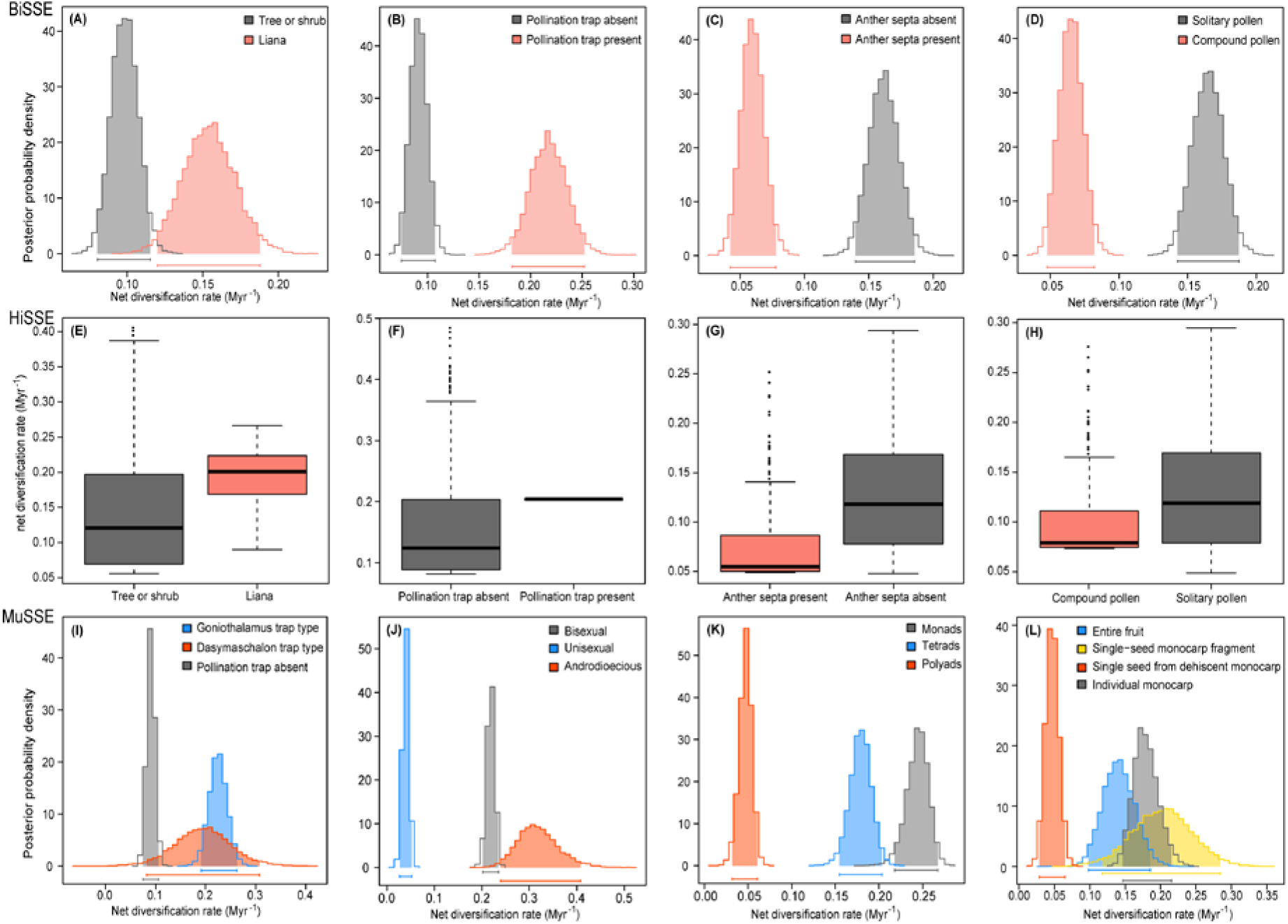
Net diversification rate comparison for traits. (**A–D**) Trait-dependent posterior distributions of net diversification rates from BiSSE analyses with 10,000 generations for four binary traits. (**E–H**) Boxplots of net diversification rates from the best HiSSE models for four binary traits. (**I–L**) Trait-dependent posterior distribution of net diversification rates from MuSSE analyses with 10,000 generations for four multi-state traits.

**Table 1.**
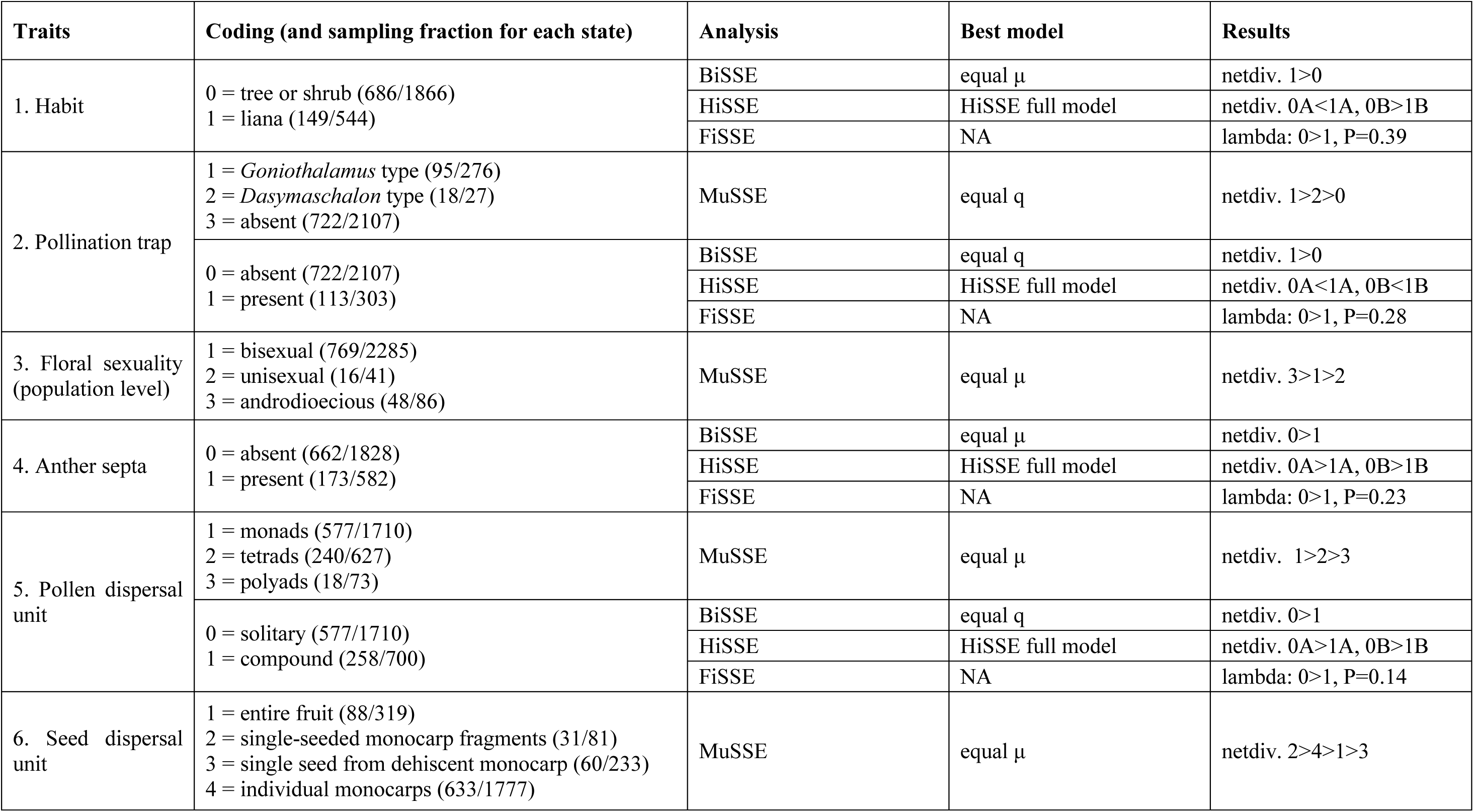
Summary of traits studied, with coding scheme, sampling fraction (*i.e.*, proportion of Annonaceae species with each character state), analyses, best model, and rate comparison results. NA, not applicable.

The Bayesian analysis under the best model of the BiSSE analysis indicated that the lianescent lineages showed higher estimates of net diversification rate than self-supporting tree or shrub lineages, with no overlap of 95% HPD intervals (Fig. 5A). Likewise, lineages with pollinator trapping (irrespective of trap type) showed higher estimates of net diversification rate than lineages without pollinator trapping (Fig. 5B). Lineages with anther septation and compound pollen dispersal units showed lower estimates of net diversification rate than those lacking anther septation and compound pollen dispersal units (Fig. 5C, D).

We compared the fit of 25 models of diversification in the HiSSE framework to our trait data coded as binary (habit, pollinator trapping, pollen dispersal unit, and anther septation). The best-supported model for all of those traits were the full HiSSE model with the lowest AIC value (*i.e.*, HiSSE models performed better than the BiSSE or the CID models; Supplementary Table S3). The HiSSE results therefore suggest that the focal traits explain some, but not all, of the diversification heterogeneity. The boxplots showed a higher mean diversification rate in lianescent lineages than in tree or shrub lineages (Fig. 5E) and in lineages with pollinator trapping than in lineages without pollination trapping (Fig. 5F), whereas a lower mean diversification rate was found in lineages with anther septation and compound pollen dispersal units (Fig. 5G, H). The FiSSE model shows differences (although not statistically significant) in speciation rates between the two categories in all binary traits (Table 1).

The MuSSE results indicate that pollinator trapping, androdioecy, and seeds that are dispersed by single-seeded monocarp fragments are positively correlated with diversification rate (Fig. 5I, J, L). In contrast, unisexual flowers, pollen dispersal as tetrads or larger polyads, and seeds that are dispersed in an entire fruit or from dehiscent monocarps negatively impact diversification rate (Fig. 5J–L).

## Discussion

Our results, based on the most comprehensive phylogenetic tree of Annonaceae to date and the use of various diversification methods, clarify the pattern of evolutionary diversification in the family. We acknowledge that some diversification methods employed are the subject of heated controversy (see Supplementary text “Potential sources of errors”). There is nevertheless clear evidence for diversification rate heterogeneity, and the observed rate shifts are associated with key traits that may provide possible explanations for the shifts in diversification rate.

### Rapid radiation in Annonaceae

Diversification analyses revealed 12 large radiations, which collectively account for over 80% of the total species richness of Annonaceae. Net diversification rates across the family slowly increased until the Miocene (*c.* 25 Ma), with subsequent higher rate increases since then (Fig. 3A). Species richness in Annonaceae is most likely the result of increased diversification (due to increased speciation: Fig. 3A) rather than gradual accumulation of species, thus favoring the ‘diversification rate’ hypothesis over the ‘museum model’ hypothesis.

Although Couvreur *et al.* [15] also found that the overall diversification rate of the family and major lineages was constant up to at least 25 Ma, incomplete species-level sampling in their study (with only 4.8% of species sampled) limited interpretation of diversification rates after this threshold. They therefore adopted an arbitrary threshold that included 85% of all generic stem nodes, and they considered a lineages-through-time (LTT) plot at the deeper nodes of the phylogeny to be accurate. The authors consequently concluded that their results for the family were consistent with the constant birth-death model, with no major diversification rate shift observed in the LTT plot, supporting the ‘museum model’ hypothesis.

In our study, the two major subfamilies, Annonoideae (with 63.5% of species diversity: [10]) and Malmeoideae (32.8% of species diversity) had the highest diversification rates, whereas the small Anaxagoreoideae (1.3% of species diversity) and Ambavioideae (2.3% of species diversity) had the lowest diversification rates (Fig. 3B), consistent with the pattern identified by Couvreur *et al.* [15]. Plotting rates-through-time for the four subfamilies indicates that the sister subfamilies Annonoideae and Malmeoideae began to diversify rapidly around 25 Ma (Fig. 3B). Changes in the net diversification rate across the phylogeny could therefore be explained largely by the dynamics of subfamilies Annonoideae and Malmeoideae. Subfamily Ambavioideae has an accelerated rate within the last 10 Ma, possibly explained by the rapid radiation of the *Cyathocalyx*-*Drepananthus* clade in South East Asia, for which taxon sampling was significantly increased in this study (to 23 spp.).

At the generic level, the species-rich genera *Annona* (170 spp.), *Artabotrys* (105 spp.), *Duguetia* (94 spp.), *Goniothalamus* (134 spp.), *Guatteria* (177 spp.), *Uvaria* (199 spp.), and *Xylopia* (164 spp.) are all associated with high diversification rates (Fig. 1). This contradicts previous studies [15,17] in which no evidence was found to support rapid diversification in large genera (except for *Goniothalamus*, in which significant increases in diversification were identified by Erkens *et al*. [17]), although previous authors may have underestimated diversification rates because their calculations were based on stem ages. Our dense species-level sampling allows us to use crown ages that are estimated to be younger than previous studies. *Goniothalamus*, for example, is regarded as a recent, rapid radiation with a crown age of *c.* 15.47 Ma in this study, much younger than the stem age estimate (*c.* 35 Ma) used by Couvreur *et al.* [15].

Our results show that net diversification rates for Annonaceae as a whole steadily increased until the Miocene (25 Mya), after which lineages generally exhibited significantly increased rates (Fig. 3A). A mixed model of steady accumulation followed by recent rapid diversification therefore seems a plausible explanation for diversification in Annonaceae. Although caution is needed when interpreting relationships between diversification rates and richness, a positive relationship between the two is not inevitable: especially when rates are faster in younger clades, the relationships may not hold [34,35]. In clades within Annonaceae, species richness is strongly correlated with diversification rate (Fig. 4B). The variation in diversification rates also seems to explain most variation in richness among clades in Annonaceae (Fig. 4B), and therefore species-rich clades may be better explained by high diversification rates than by the ‘museum model’ hypothesis. Extrinsic abiotic factors (such as biogeography, climate, Asian monsoon, sea level change, habitat, and range size) may have contributed to increased diversification rates within Annonaceae. Such abiotic factors are beyond the scope of our study, however; here we explore the possible association of intrinsic factors and diversification rate shifts. The relationship between rates and diverse abiotic factors is worthy of exploration in future studies.

### Mechanisms linking traits to diversification

Although it is impossible to unequivocally identify selected traits as the cause of diversification rate shifts, our results provide evidence of strong correlations between some traits and rates. Our trait-dependent analyses show that the presence of circadian pollinator trapping, androdioecy, and the dispersal of seeds as single-seeded monocarp fragments are all correlated with higher diversification rates. Some observed shifts in diversification rate identified by BAMM and turboMEDUSA also match with some observed transitions of character states. Pollen aggregation and anther septation, in contrast, are linked with lower diversification rates.

#### Habit

The self-supporting habit (as trees and/or shrubs) was reconstructed as the most likely ancestral state for Annonaceae, with three independent origins of the lianescent habit, in *Artabotrys*, the *Pseudartabotrys*-*Letestudoxa* clade (tribe Duguetieae), and tribe Uvarieae (consistent with previous results: [36]). Reversals to shrubs occurred three times: once each in *Cleistochlamys*, *Dielsiothamnus*, and *Dasymaschalon*, all in tribe Uvarieae (Supplementary Fig. S4A). The distinction between lianas and self-supporting trees and shrubs is weak in Annonaceae [37], with most ‘climbers’ adopting a rather scandent habit. *Artabotrys* is unusual in possessing highly specialized climbing hooks (shown to be evolutionarily derived from inflorescence stalks: [38]). Although Annonaceae have a pantropical distribution, the lianescent species (all in subfamily Annonoideae) are restricted to the Paleotropics.

The lianescent habit is clearly associated with three clades with accelerated diversification rates based on the BAMM analysis: *Artabotrys*, *Uvaria*, and the *Desmos*-*Dasymaschalon*-*Friesodielsia*-*Monanthotaxis* clade. Trait-dependent diversification analyses also indicate that the liana habit is associated with higher diversification rates (Fig. 5A, E). Unlike trees or shrubs, lianas have a reduced requirement for structural support (xylem biomass) and can therefore allocate a greater proportion of their resources to reproduction, canopy development, and stem and root elongation [39,40]. It has been suggested that the enhanced vascular efficiency of lianas enables them to better cope with water stress conditions [41]; in this context, it is noteworthy that *Artabotrys* and *Uvaria* appear to demonstrate a broader habitat tolerance with regard to precipitation seasonality [42]. In contrast to these lianas with high diversification rates, the *Pseudartabotrys*-*Letestudoxa* clade comprises lianas but has a low diversification rate. This result is consistent, however, with the view that the origin of a trait may not, in itself, be sufficient to increase diversification rate, but instead requires an appropriate combination of traits [43].

#### Pollinator trapping

Most species of Annonaceae possess partially enclosed pollination chambers with apertures that remain open throughout anthesis, enabling pollinators to enter or leave the flowers freely during any phenological stage [20]. Tightly enclosed floral chambers that trap pollinators are rare in the family, but are reported in *Dasymaschalon*, *Friesodielsia*, *Goniothalamus*, and possibly also *Artabotrys* [23,24,44] (Supplementary Fig. S4B). The opening and closing of the floral chambers are precisely aligned with the endogenous periodicities of the pollinating beetles: opening of the flowers coincides with an activity peak, but the floral chamber closes before the beetles enter their next circadian peak [24] (Fig. 6A, B). This timing broadens the range of potential pollinators to include those with circadian rhythms that are both unimodal (with a single daily activity peak) and bimodal (with twice-daily peaks) [24]. Circadian trapping also endows other selective advantages, including the possibility of an extended staminate phase to promote pollen deposition and enhanced inter-floral movement of pollinators. This trapping mechanism is likely to be synapomorphic for each clade and hence may represent a key evolutionary innovation: we demonstrate that *Artabotrys*, *Dasymaschalon*, *Friesodielsia*, and *Goniothalamus* all show accelerated diversification (Fig. 1); lineages with pollinator trapping also show higher rates in trait-dependent analyses (Fig. 5B, F, I).

**Fig. 6.**
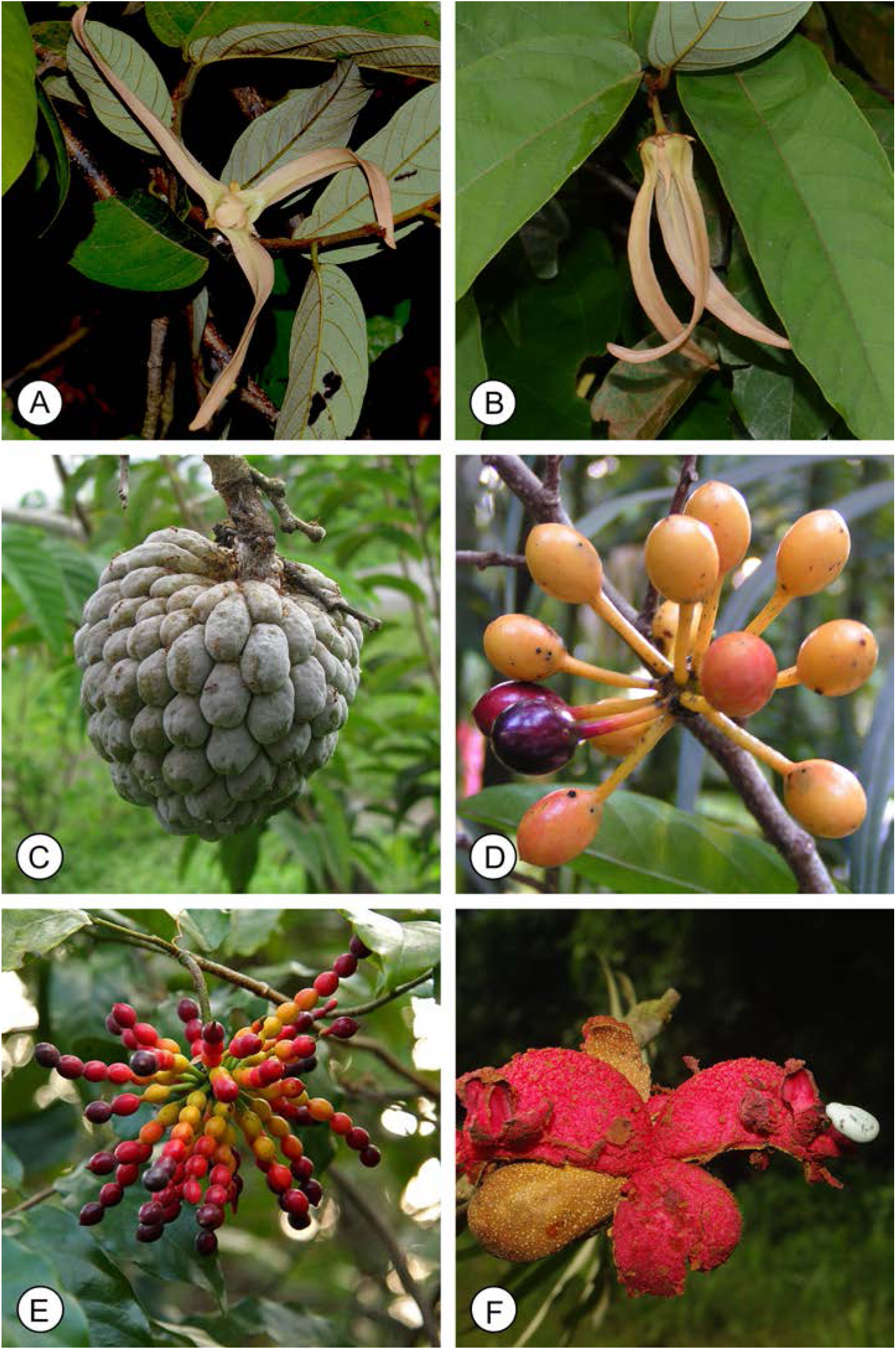
Pollinator trapping, and selected Annonaceae fruits showing different dispersal units. (**A**) *Friesodielsia borneensis*: floral chamber open. (**B**) *Friesodielsia borneensis*: staminate phase, floral chamber closed. (**C**) *Annona cherimola*: entire fruit acts as a single dispersal unit. (**D**) *Phaeanthus ebracteosus*: individual monocarps are the dispersal units. (**E**) *Desmos chinensis*: single-seeded monocarp segments are the dispersal units. (**F**) *Xylopia hypolampra*: monocarps dehisce, exposing seeds that are the dispersal units. (Photos: **A, B**, X. Guo; **C**, X. Cornejo, http://annonaceae.myspecies.info/; **D**, R.M.K. Saunders; **E**, C.C. Pang; **F**, T.L.P. Couvreur, http://annonaceae.myspecies.info/).

Although anthesis usually extends over 48 h in species of Annonaceae with hermaphroditic flowers [23,24], *Artabotrys*, *Dasymaschalon*, *Friesodielsia*, and *Goniothalamus* species, which all exhibit circadian trapping, also have an abbreviated anthesis of only 23–27 h [23,24,44]. A similarly brief anthesis (*c.* 27 h) is reported for *Desmos chinensis* [22], although this species does not trap pollinators. *Desmos* is sister to the *Dasymaschalon-Friesodielsia* clade [18], and hence abbreviated anthesis is likely to be synapomorphic for the entire clade and can be inferred to have evolved prior to pollinator trapping [24].

Annonaceae species with abbreviated anthesis often display pistillate/staminate-phase floral synchrony, in which pistillate-phase and staminate-phase flowers are not borne concurrently on an individual, thereby minimizing opportunities for geitonogamy [22,23,24,44]. Lau *et al.* [23] hypothesized that shortening of anthesis in species with this type of floral synchrony reduces the proportion of non-flowering days and hence increases seedset. Abbreviated anthesis and pistillate/staminate-phase floral synchrony might therefore be prerequisites for circadian pollinator trapping.

Circadian trapping is likely to have evolved independently in the above-mentioned lineages, reflecting the considerable selective advantages of the mechanism [45]. In addition to the four genera listed above, in which pollinator trapping is supported by empirical field studies, several other genera are likely to exhibit circadian trapping based on assessments of their floral morphology, *viz*. *Cyathocalyx*, *Drepananthus*, *Mitrella*, *Neostenanthera*, *Pseudartabotrys*, and *Trivalvaria macrophylla* (although no other species of *Trivalvaria*). Little information on the pollination ecology of these taxa is available, however. Although field observations on floral phenology and petal movement, as well as pollinator activities, are crucial for unequivocal evaluations of whether a pollinator trapping mechanism exists, trait-dependent diversification analyses based on a data set in which these taxa are coded as having pollination traps show similar results.

#### Population-level floral sex expression

Although floral hermaphroditism is widespread in Annonaceae, androdioecy occurs in several lineages and some populations comprise exclusively unisexual flowers (Supplementary Fig. S4C). Androdioecy is rare in angiosperms as a whole, but appears to be relatively common within Annonaceae, especially in tribes Miliuseae (*c.* 6%) and Malmeeae (*c.* 16.7%) [44].

Our MuSSE results suggest that androdioecy is associated with the highest diversification rate, whereas lineages in which plant populations comprise exclusively bisexual flowers are associated with the lowest diversification rate (Fig. 5J). The diversity of floral sex expression at the population level in tribes Miliuseae and Malmeeae—with very frequent occurrences of androdioecy and exclusively unisexual flowers—may provide an explanation for the high diversification rates in these tribes. Androdioecy has been identified as potentially advantageous in obligately outcrossing species in which gene flow among individuals is limited because the reduced level of pistillate function in the population may be counter-balanced by increased seedset in bisexual flowers due to the greater availability of pollen [46]. Although Annonaceae species are self-compatible [44], they have evolved various mechanisms that promote xenogamy, including protogyny (which precludes autogamy) and pistillate/staminate-phase floral synchrony (which minimizes opportunities for geitonogamy). The beetle pollinators of most species of Annonaceae are comparatively sedentary, rarely initiating flight to move between flowers: inter-floral pollen transfer might therefore be promoted by the increased availability of pollen in androdioecious populations.

#### Anther septation

Most Annonaceae species have aseptate anthers, lacking any internal division within the thecae during pollen development, and this has been inferred as plesiomorphic for the family (Supplementary Fig. S4D). Other species have thecae that are at least temporarily compartmentalized into internal chambers with a single compound pollen unit in each chamber [47]. The occurrence of compound pollen and the formation of septate anthers are strongly correlated in Annonaceae [47,48]: this combination may have evolved to enhance contact between the tapetum and the developing pollen, meeting the greater nutritive demands of large pollen aggregates [47]. As with pollen aggregation (discussed below), the occurrence of aseptate anthers is also associated with low diversification rates (Fig. 5C, G), although this may reflect trait correlation.

#### Pollen dispersal unit

The pollen of most Annonaceae species is dispersed as solitary grains (monads) at maturity. This state is inferred as plesiomorphic for the family, with compound pollen (tetrads and larger polyads) derived in several phylogenetically disparate lineages (Supplementary Fig. S4E). These results are consistent with previous studies on pollen evolution in the family [49,50].

Pollen aggregation has been regarded as a mechanism that enhances the efficiency of pollination by enabling the fertilization of multiple ovules following a single pollinator visit [51,52]. The process of pollen aggregation is likely to be particularly important in species that are pollinated by rare or inconstant pollinators, such as flies (*Asimina*: [53]; *Disepalum*: [54]; *Monodora*: [55]; *Pseuduvaria*: [56,57]; *Uvariopsis*: [55]). In apocarpous flowers (which constitute *c.* 98% of Annonaceae species), however, the transfer of pollen aggregates would only be beneficial in cases where there are either multiple ovules per carpel or an effective extragynoecial compitum enabling intercarpellary growth of pollen tubes. Most genera that possess pollen aggregates have multiple ovules per carpel, *viz.*: *Cymbopetalum* (up to 25 ovules), *Goniothalamus* (up to 10), *Meiocarpidium* (up to 20), *Monodora* (up to 70), *Porcelia* (up to 15), *Pseuduvaria* (up to 18), *Porcelia* (up to 15), *Uvariastrum* (up to 25), and *Uvariopsis* (up to 15). Some genera of Annonaceae that possess pollen aggregates have few ovules per carpel, including *Annona* (1 ovule), *Anonidium* (1), *Disepalum* (1–3), *Duckeanthus* (1) and *Fusaea* (1). In these cases, there are numerous carpels in each flower, and the copious stigmatic exudate that forms during the pistillate phase acts as an extragynoecial compitum enabling the intercarpellary growth of pollen tubes [21]. Despite the apparent advantages of pollen aggregation, our trait-dependent diversification analyses indicate that this trait is associated with low diversification rates (Fig. 5D, H, K). Pollen aggregation is likely to result in limited paternal diversity of seeds within a fruit. One hypothesis is that this trait might be correlated with a very effective seed dispersal mechanism, ensuring that genetically closely related seeds are not dispersed together but are spatially scattered (discussed below).

#### Seed dispersal unit

There are relatively few reliable field observations of seed dispersal in Annonaceae. We therefore opted to supplement these sporadic reports by coding this trait based on interpretations of fruit and seed morphology. Most species of Annonaceae have fruits in which the base of each monocarp (the fruit unit derived from a single carpel in apocarpous flowers) is extended to form a stipe that ensures separation of monocarps at maturity (Fig. 6D); in these cases, individual monocarps often mature at different rates and are dispersed separately, either by birds that swallow the monocarps whole and defecate the seed intact, or by primates that regurgitate the seeds. The dispersal of monocarps separately is reconstructed as the plesiomorphic trait for the family (Supplementary Fig. S4F).

Other species of Annonaceae develop relatively large fruits in which the anatomically separate monocarps become closely appressed (*i.e.*, functionally syncarpous) due to the limited development of monocarp stipes; in these cases, the dispersal unit is the entire fruit, with dispersal generally effected by relatively large frugivores such as primates (*e.g.*, *Duguetia*, *p.p.*: [58]; and *Goniothalamus*, *p.p.*: [59]). Some genera, including *Annona* and *Duguetia*, *p.p.*, undergo differing degrees of post-fertilization fusion of carpels (Fig. 6C; [60]), resulting in ‘pseudosyncarpy’, in which the entire fruit is dispersed as a single unit. Entire fruits also function as the dispersal unit in *Isolona* and *Monodora* (and possibly also *Cyathocalyx*), in which ‘true’ syncarpy has been demonstrated [36,61,62] (Supplementary Fig. S4F). Monocarps of many species in the *Monanthotaxis-Dasymaschalon-Desmos* clade are elongated and moniliform, with constrictions between seeds (Fig. 6E; [18]); each monocarp ripens progressively from apex to base, with single-seeded segments removed sequentially as they ripen by avian frugivores. Within Annonaceae, this functional dispersal unit is unique to this clade (Supplementary Fig. S4F), although it parallels the ‘lomentum’ of some legumes [63]. Several genera have shifted the dispersal unit from fruits to individual seeds (Supplementary Fig. S4F): *Cardiopetalum*, *Cymbopetalum*, *Trigynaea* [64], and *Xylopia* [65], for example, possess dehiscent monocarps that split along a dorsal suture to expose bird-dispersed seeds with a brightly colored aril or sarcotesta (Fig. 6F). Some *Anaxagorea* species have dehiscent monocarps with ballistic dispersal of non-arillate seeds over distances up to 5 m [66].

Our MuSSE results show that the evolution of single-seeded monocarp fragments as the dispersal unit is associated with the highest diversification rate, with individual monocarp dispersal associated with the second-highest diversification rate, and entire fruit dispersal the third-highest. Direct seed dispersal from dehiscent monocarps is associated with the lowest diversification rate (Fig. 5L).

The evolution of syncarpy has often been considered a key evolutionary innovation in angiosperms [67]. As well as promoting pollination efficiency by increasing seedset and minimizing ovule wastage due to enhanced ovule access by pollen tubes, syncarpy is also likely to prove selectively advantageous after fruit development, enhancing the protection of ovules and developing seeds and possibly also contributing to seed dispersal [68,69]. Annonaceae seeds within a single fruit are likely to exhibit only limited paternal diversity due to the comparatively sedentary nature of the beetle pollinators and the dispersal of aggregated pollen in many species (see above); we therefore hypothesize that apocarpy might have been retained in Annonaceae flowers as it would potentially enhance the spatial dispersal of seeds by promoting the separate dispersal of fruit monocarps. Speciation might therefore have been facilitated as a consequence of geographic isolation of these propagules and the increased opportunities for establishing isolated populations.

The spatial separation of paternal seed types might be further enhanced by the dispersal of single-seeded monocarp segments and the evolution of dehiscent monocarps in which seeds are dispersed separately. Our results show that dispersal of single-seeded monocarp fragments is associated with the highest diversification rate; this is furthermore consistent with results from the trait-independent analyses that show that the *Monanthotaxis-Dasymaschalon-Desmos* clade has diversified rapidly.

#### Traits not assessed here

Other traits that may have potentially impacted diversification rates in Annonaceae were not included in our analyses due to the scarcity of data or because of the limited statistical power associated with small sample sizes, *viz*. polyploidy, primary pollinator guilds, pistillate/staminate-phase floral synchrony, and anthesis duration.

Polyploidy is undoubtedly a major force in angiosperm evolution [70]. It has been reported in several genera of Annonaceae, including: *Cyathocalyx*, with 2*x* and 8*x* taxa recorded [71]; *Duguetia*, with 2*x*, 3*x*, 4*x*, and 6*x* taxa [58,72,73]; *Annona*, with 2*x*, 4*x*, 6*x*, and 8*x* taxa [71,72,73,74]; and *Guatteria*, which are uniformly 4*x* [71,75,76]. In particular, these four genera all exhibit rapid diversification rates (Fig. 1; Supplementary Table S1): genome duplication via allopolyploidy provides a potential explanation for this pattern (*e.g.*, [77]).

There is a possible correlation in Annonaceae between primary pollinator guild and anthesis duration (fly-pollinated species, for example, typically exhibit extended anthesis: [78]), as well as between abbreviated anthesis duration, pistillate/staminate-phase floral synchrony, and the circadian trapping of pollinators (as discussed earlier in this paper, abbreviated anthesis is likely to increase seedset by minimizing the number of non-flowering days when associated with floral synchrony and pollinator trapping). The limited availability of data precluded further assessment, however.

## Conclusions

Our results show that variation in diversification rates explain most variation in richness among clades in Annonaceae. Species richness in Annonaceae is therefore better explained by the diversification rate hypothesis. Net diversification rates for the family steadily increased until the Miocene (ca. 25 Ma), after which the lineages generally exhibited increased rates of diversification (Fig. 3A). A mixed model of steady accumulation followed by recent rapid diversification therefore seems a plausible explanation for diversification in Annonaceae. Increased rates of diversification either during or after the Miocene have also been reported for other groups, including Saxifragales [79], as well as clades of Ericales and rosids [80], among others.

BAMM, turboMEDUSA, and the method-of-moments estimator all indicate heterogeneity in diversification rates across the phylogeny; for example, 12 shifts were identified in the BAMM analyses, and results from the other methods are generally consistent with these patterns. Trait-dependent diversification analyses show that the liana habit, the presence of circadian pollinator trapping, androdioecy, and the dispersal of seeds as single-seeded monocarp fragments are all correlated with higher diversification rates; pollen aggregation and anther septation, in contrast, are linked with lower diversification rates. Whether any of these features has actually directly served as a driver or barrier to diversification is unknown and requires further investigation.

## Materials and Methods

### Taxon and DNA region sampling

We adopted a super-matrix approach, using the recently published phylogeny by Guo *et al.* [10], supplemented with additional sequences from other sources [18,42,81,82,83,84,85,86,87,88]. Eight regions of the plastid genome that are commonly used in Annonaceae phylogenetics (*rbcL*, *matK*, *ndhF*, *psbA-trnH*, *trnL-F*, *atpB-rbcL*, *trnS-G* and *ycf1*) were downloaded from the nucleotide database of the National Center for Biotechnology Information (http://www.ncbi.nlm.nih.gov). Species with fewer than three of the above-mentioned DNA regions were not included, following Guo *et al.* [10]. Seven species belonging to five other families in Magnoliales were used as outgroups: *Myristica fragrans* and *Coelocaryon preussii* (Myristicaceae), *Galbulimima belgraveana* (Himantandraceae), *Degeneria roseiflora* (Degeneriaceae), *Magnolia kobus* and *Liriodendron chinense* (Magnoliaceae), and *Eupomatia bennettii* (Eupomatiaceae). The final matrix therefore comprised 923 accessions (including 916 ingroup and seven outgroup taxa, representing *c.* 98% of generic diversity and *c.* 35% of species diversity in Annonaceae). GenBank accession numbers are provided in Supplementary Appendix 1.

### Alignment and phylogenetic analyses

Sequences of individual regions were aligned automatically using the MAFFT plugin in Geneious ver. 11 [89] with default settings and then manually edited and optimized. A total of 449 ambiguously aligned positions (including inversions and short repetitive sequences) in *trnL-F* and *psbA-trnH* were excluded from the analyses. The final concatenated alignment for the data set with 923 terminals consisted of 11,211 positions.

Maximum likelihood (ML) analyses were performed using RAxML ver. 8.2.10 [90] provided by the CIPRES Science Gateway [91]. The supermatrix was partitioned into eight partitions corresponding to gene or spacer region. Fifty inferences were run under the GTR+Γ model with 1,000 non-parametric bootstrap replicates.

### Divergence time estimation

Due to the data size and convergence issues in BEAST [92], we dated the tree using the penalized likelihood approach [93] as implemented in the program TreePL [94], which can handle large-scale phylogenies. This program allows better optimization with large trees by combining stochastic optimization with hill-climbing gradient-based methods. The algorithm estimates evolutionary rates and divergence dates on a tree given a set of fossil constraints and a smoothing factor determining the amount of among-branch rate heterogeneity. This strategy incorporates a strong constraint on the root [93] and some other nodes, while estimating the remainder. The root was fixed at 137 Ma according to Foster *et al.* [95], as well as the occurrence of unequivocal angiosperm crown group pollen grain fossils from the Hauterivian (Early Cretaceous, 136.4–130 Ma: [96,97]). The Annonaceae crown node was constrained between 112.6–89 Ma, and the Magnoliineae crown node between 125–112.6 Ma, following calibration scheme CS3 (*sensu* [19,42]). We used the best ML tree from RAxML analyses as input. A priming analysis was conducted to determine the best optimization parameter values, and the optimal smoothing parameter value was then determined as 1,000 which is corresponding to the highest Chisq value of 69648.4 using a cross-validation analysis.

### Diversification rate analysis

To avoid putative inflation of evolutionary rates due to the inclusion of multiple samples per species for the diversification rate analysis [98], duplicate taxa and taxa considered as subspecies of species already included in the tree were pruned from the rate-smoothed 923-tip RAxML tree. Meanwhile, undescribed or unidentified species as well as the outgroup were also excluded (although in this case, re-running the analysis with such tips did not have a qualitative effect on the results; the same rate shifts were identified as in the pruned phylogeny). In total, 88 tips were pruned from the rate-smoothed RAxML tree, resulting in an 835-taxon dated tree. All the following diversification analysis were based on this 835-taxon dated tree.

We applied a range of methods to evaluate the diversification pattern of Annonaceae. Selected methods were of two types: trait independent and trait dependent. Three trait-independent diversification methods to detect diversification rate shifts were employed: modelling evolutionary diversification using stepwise AIC (turboMEDUSA; [27]), Bayesian analysis of macroevolutionary mixtures (BAMM; [28]), and method-of-moments estimator [29]. Four methods to detect trait-dependent diversification rates were applied: binary-state speciation and extinction (BiSSE; [30]); multiple state speciation extinction (MuSSE; [31]); fast, intuitive state-dependent speciation and extinction (FiSSE; [32]); and hidden state speciation and extinction (HiSSE; [33]).

### Species richness data

Species richness data for each genus were obtained from Guo *et al.* [10], supplemented by data from recent taxonomic publications [85,88,99,100,101,102]. Several recognized genera are not monophyletic, and hence a total of 101 ‘genus-level clades’ were adopted. Details of the taxa used are as follows: (1) the monospecific genus *Schefferomitra* is nested within *Friesodielsia* and should therefore be congeneric with it [18,103]; *Friesodielsia*-*Schefferomitra* is therefore treated as a single genus-level clade with 39 species. (2) *Stenanona* is nested within *Desmopsis* [86]: *Stenanona* and *Desmopsis* are therefore combined, leading to the recognition of a clade with 28 species. (3) *Klarobelia*-*Pseudephedranthus*-*Pseudomalmea* are nested within *Oxandra*, a relationship that was also revealed in a recent phylogenomic study [104]; the four genera were collectively recognized as a single genus-level clade with 45 species. (4) *Dasymaschalon* is polyphyletic, with three species (*D. filipes*, *D. longiflorum*, and *D. tibetense*) of intergeneric hybrid origin [45,105], which were therefore excluded from our study. (5) *Winitia* was included in *Stelechocarpus*, following recommendations by Turner [106] and Chatrou *et al.* [13]. The richness data are provided in Supplementary Table S5.

### BAMM

We evaluated diversification in Annonaceae using Bayesian Analysis of Macroevolutionary Mixtures software (BAMM) ver. 2.5.0 [28] on the high-performance computing cluster (HiPerGator) at the University of Florida (Gainesville, Florida, USA) and subsequently analyzed with the R package BAMMtools ver. 2.1.6 [107] using the 835-taxon tree. BAMM implements the reversible jump Metropolis Coupled Markov Chain Monte Carlo method and allows for both time-dependent speciation rates as well as discrete shifts in the rate and pattern of diversification. Priors were configured using ‘setBAMMpriors’ function in ‘BAMMtools’ on the 835-taxon tree (Supplementary Appendix 2). To account for incomplete taxon sampling, sampling fraction data for each genus-level clade are provided (Supplementary Table S5).

The BAMM analysis was run for 20,000,000 generations, sampling every 10,000 generations. After removing 10% of the output as burn-in, the likelihood of all sampled generations was plotted in R, and effective sample size (ESS) values for the likelihood and the inferred numbers of shifts were calculated using the package coda ver. 0.19-1 to assess convergence [108]. The post burn-in output files were analyzed using ‘BAMMtools’ by assessing posterior probabilities for the selected best rate-shift configuration and the 95% credible rate shift configuration.

Rates-through-time plots were also generated for speciation (*λ*), extinction (*µ*), and net diversification (*r*) using ‘PlotRateThroughTime’ for the entire family as well as specific subfamilies or lineages.

We are well aware of the controversy surrounding BAMM [109,110,111; but see 112,113] (see Supplementary text “Potential Sources of Errors”). We accordingly compared our BAMM results with other methods, including turboMEDUSA and method-of-moments estimator.

### turboMEDUSA

Patterns of diversification were also analyzed in R [114] using the Modeling Evolutionary Diversification Using Stepwise Akaike Information Criterion (MEDUSA) package, ver. 0.952 ([27]; https://github.com/josephwb/turboMEDUSA).

The 835-taxon tree was condensed to a 101-tip genus-level tree using Newick utilities ver. 1.6 [115] (Supplementary Fig. S5). The species richness data (Supplementary Table S5) were included to assign unsampled taxa to each genus or genus-level clade to account for incomplete taxon sampling. The mixed model and Akaike information criterion (AIC) threshold criterion were used. MEDUSA selects the best model (pure birth or birth-death) for each shift and determines the significance of shifts based on Akaike information criterion (AICc) scores corrected for sample size. The net diversification rate (*r*) = speciation (λ) − extinction (μ), and relative extinction rate (ε) = μ/λ for each diversification rate shift were calculated.

### Method-of-moments estimator

We also estimated net diversification rates for each subfamily, tribe, and the genera or clades that show rate shifts in BAMM using the method-of-moments estimator for stem-group ages and crown-group ages [29] using the package geiger ver. 2.0.6 [116]. This method uses clade ages, species richness, and a correction for failing to sample extinct clades (relative extinction fraction, ɛ). Following standard practice, we used three different values of ε: two extreme values (0 and 0.9) and an intermediate value (0.45) [117,118].

### Museum model *vs* diversification rate hypotheses

To analyze relationships between species richness and age or diversification rate in the 101 genus-level clades in Annonaceae, we used phylogenetic generalized least-squares regression (PGLS) models [119] to account for phylogenetic non-independence of clades, using the R package caper ver. 1.0.1 [120]. Following standard practice, delta (δ) and kappa (κ) were set to 1 while the maximum-likelihood value of lambda (λ) was estimated for each analysis and used to transform branch lengths. Richness and clade age were ln-transformed to improve linearity [1].

The 101-tip genus-level tree (Supplementary Fig. S5) was used as the input tree. Stem age was used as clade age in the analysis as crown age could not be estimated for monotypic clades. Net diversification rates were estimated from method-of-moments estimators for stem-group ages following standard practice [28]. The richness, clade age, and net diversification rate data used in the analysis are provided in Supplementary Table S6.

### Character coding and ancestral state reconstruction

Based on data availability, we scored the following six traits for all 835 species in the data set. Trait (1): habit (self-supporting trees/shrubs *vs* lianas). Trait (2): occurrence of circadian trapping of pollinators (sensu [24]), scored in two ways, *viz.*: (a) a simple present *vs* absent coding; or (b) further discriminating between *Dasymaschalon*-type pollinator trapping (in which the trap chamber is formed by the three petals, homologous with the inner petals of other Annonaceae) and *Goniothalamus*-type pollinator trapping (in which the trap chamber is formed by all six petals). Trait (3): population-level floral sex expression, in which populations comprise exclusively bisexual flowers, unisexual flowers, or are androdioecious. Trait (4): anther septation (present *vs* absent). Trait (5): pollen dispersal unit, scored in two ways, *viz.*: (a) monad *vs* compound pollen coding; or (b) further discriminating among monads, tetrads, and larger polyads. Trait (6): seed dispersal unit (entire fruit, individual monocarps, single-seeded monocarp fragments, or single seed dispersal from dehiscent monocarps). For trait 6, there are very few reliable descriptions of seed dispersal in Annonaceae, and we therefore coded this trait based on fruit/seed morphology and sporadic field records from references and specimen labels.

The states for all traits (summarized in Table 1) were obtained from the literature on Annonaceae, available specimens, photographs, and the authors’ unpublished observations. Other traits that may have potential impact on diversification rate but with limited available data (*e.g.*, polyploidy, primary pollinators, and duration of anthesis) were not included in the analyses because of the limited power for statistical analysis for small sample sizes.

To reconstruct the ancestral state for each trait, we used stochastic character mapping of discrete traits via SIMMAP [121] using the package phytools ver. 06-44 [122] implemented in R. The best-fit model of character evolution was determined by fitting an equal-rates model and an all-rates-different (ARD) model to the data set using the function ‘fitDiscrete’ in geiger ver. 2.0.6 [116]. Model fit was compared using the AICc [123]. The ARD model had the best fit to all the data. We simulated 1000 stochastic character maps from the data set using the ARD model and obtained posterior probabilities for the nodes by averaging the state frequencies across all maps.

Uncertainties in character state coding were treated as missing data in the analyses. Species that did not have clearly defined character states were excluded from the appropriate analyses. For traits with both binary and multistate coding schemes, the multistate coding was used for ancestral state reconstruction analyses to better reflect current hypotheses of homology.

### Trait-dependent diversification

To test whether certain trait states are associated with increased speciation and diversification rates, we applied diverse models, with BiSSE, HiSSE, and FiSSE used to analyze binary traits (pollination trap, habit, pollen dispersal unit, and anther septation) and MuSSE used to study multistate traits (pollination trap, floral sexuality at population level, and seed dispersal unit). All analyses were performed on a pruned phylogeny excluding taxa with missing trait data.

The BiSSE model [124] was implemented in the R package diversitree ver. 0.9-10 [31]. For each trait, we evaluated four BiSSE diversification models using ML searches: a full model allowing all variables to change independently, a model constraining speciation rate values across states to be equal (λ_0_ ∼ λ_1_), a model constraining extinction rates to be equal (μ_0_ ∼ μ_1_), and one constraining transition rates to be equal (q_01_ ∼ q_10_). The best-fit model was selected based on the lowest AIC value. To obtain the posterior distributions of the parameters, we conducted a Bayesian BiSSE analysis with the best-fit model for each trait. This was run for 10,000 generations using an exponential prior with a rate of 1/(2*r*), where *r* is the diversification rate of the character [30]. To correct for non-random, incomplete sampling, we specified sampling fractions (*i.e.*, the proportion of species in state 0 and in state 1 that are included in the tree) [30]. The results were visualized as posterior probability distribution plots.

To avoid the possibility of type I errors in BiSSE (incorrectly finding neutral traits correlated with higher diversification rates: [125]), we also tested the hypothesis that shifts in the binary traits correlate with shifts in diversification rates by applying hidden state speciation and extinction analyses (HiSSE: [33]). HiSSE uses a hidden Markov model to allow an unmeasured alternative trait that is partially co-occurring with the observed trait. The hidden state may drive differences in diversification rates among taxa in addition to (or instead of) the observed state. For HiSSE, we tested the same 24 models proposed by Beaulieu and O’Meara [33], together with a HiSSE model with all possible rates varying independently (following Landis *et al.* [126]). These 25 models included four models corresponding to the BiSSE analysis with a variety of constrained parameters, 17 HiSSE models that assumed a hidden state associated with both observed character states with a variety of turnover rates, extinction rates, and transition rates constrained, and four trait-independent models. The model with the lowest AIC was preferred and considered strongly supported given ΔAIC ≥ 4 from the next value [123].

To reduce false positives and to help validate results from the other SSE analyses, we furthermore applied FiSSE to compare the speciation rates between the two states of binary traits [32]. FiSSE is a semi-parametric and likelihood-free method and is less susceptible to false positives than other SSE approaches [127].

To estimate speciation and extinction rates for multistate traits, we used the MuSSE model implemented in the R package diversitree ver. 0.9-10 [31]. Three parameters were included for each state: a speciation rate, λ, an extinction rate, μ, and a transition rate between different states, q. We compared the relative fit of the data with eight likelihood models, each with different combinations of parameters that were either free to vary among states or constrained to be equal among states: (1) λ free, μ free, and q free (*i.e.*, separate values for each parameter estimated for each state); (2) λ equal between all states, μ free, and q free; (3) λ equal, μ equal, and q free; (4) λ equal, μ equal, and q equal; (5) λ free, μ equal, and q equal; (6) λ equal, μ free, and q equal; (7) λ free, μ free, and q equal; and (8) λ free, μ equal, and q free. We compared their relative fit using AIC [123]. The Chi-squared value (ChiSq) and their significance (Pr) were calculated by comparison with the minimal model. We accounted for incomplete sampling in our tree for each state using a sampling fraction. We estimated the posterior density distribution of parameters for the best-fitting model with Bayesian MCMC analyses (100,000 steps) to estimate speciation, extinction, and transition rates. The results were visualized as posterior probability distribution plots.

## Acknowledgements

This research was supported by grants from National Natural Science Foundation of China (Grants 31872646 and 31400199) awarded to B. Xue, and the Hong Kong Research Grants Council (Grants 17112616 and 776713M) awarded to R.M.K. Saunders. B. Xue was supported by the China Scholarship Council as a visiting scholar for one year at the University of Florida, Gainesville, FL, USA. We thank Matt Gitzendanner for computational help with HiperGator; Daniel Thomas, Ryan Folk, Chaonan Fu, Lei Yang, Rebecca Stubbs, Haifei Yan, Tingting Duan, Zhonglai Luo, Zhongtao Zhao, Yuan Xu, Andrea Sanchez, Thomas Marcussen, Ranjit Sahoo, Evgeny Mavrodiev, Andres E. Ortiz-Rodriguez, and Joshua P. Scholl for sharing data alignment, scripts, or discussion.

## SUPPLEMENTARY MATERIAL

**Supplementary Text.** Potential sources of errors.

**Supplementary Table S1.** Estimation of diversification rates using method-of-moments estimator following [29].

**Supplementary Table S2.** Comparison of the fit of different BiSSE models. Gray boxes denote the best-fitting model. Abbreviations: Df = degrees of freedom; lnLik = log-likelihood; AIC = Akaike Information Criterion; ChiSq = Chi-squared value compared to the minimal model; Pr = *P*-value; λ = speciation rate; μ = extinction rate; q = transition rate; ΔAIC difference between the model and best model; NA = Not Applicable.

**Supplementary Table S3.** Comparison of the fit of different HiSSE models. Gray boxes denote the best-fitting model based on ΔAIC. Abbreviations: λ = speciation rate; μ = extinction rate; q = transition rate; τ = turnover rates; ε = extinction fraction.

**Supplementary Table S4.** Comparison of the fit of different MuSSE models. Gray boxes denote the best-fitting model. Abbreviations: Df = degrees of freedom for each model; lnLik = log-likelihood of the model; ChiSq = Chi-squared value compared to the minimal model; Pr = *P*-value; λ = speciation rate; μ = extinction rate; q = transition rate between two different regions; ΔAIC = AIC difference between the model and best model.

**Supplementary Table S5.** Species richness data and sampling fraction for BAMM and turboMEDUSA analyses.

**Supplementary Table S6.** Species richness, clade age, and net diversification rate for 101 genus-level clades used in PGLS analysis.

**Supplementary Fig. S1.** RaxML tree with 923 terminals.

**Supplementary Fig. S2.** Dated RaxML tree with 923 terminals.

**Supplementary Fig. S3.** Pruned ultrametric tree with 835 species for diversification analyses, showing family phylogeny and clade ages.

**Supplementary Fig. S4.** Ancestral character reconstruction for the six traits in Annonaceae using stochastic character mapping. (A) Habit. (B) Pollination trap. (C) Floral sexuality at population level. (D) Anther septa. (E) Pollen dispersal unit. (F) Seed dispersal unit.

**Supplementary Fig. S5.** 101-tip genus-level tree used for turboMEDUSA and PGLS analyses.

**Supplementary Appendix 1.** Species and GenBank accession numbers.

**Supplementary Appendix 2.** Priors for BAMM analyses.

