## Supplementary material for "Accelerated diversification correlated with functional traits shapes extant diversity of the early divergent angiosperm family Annonaceae"

#### Potential sources of errors

Additional sequence data—including *Monanthotaxis* [1], the newly described genus *Leoheo* Chaowasku, D.T. Ngo & H.T. Le [2], and newly described species and updated species richness estimates (*e.g.*, [3])—unfortunately could not be incorporated into our study since the data was available after we started this study. We repeated our analyses using different species richness estimates, however, including the original species richness data from Guo *et al.* [4] and Chatrou *et al*. [5], and found that minor difference in species numbers do not affect the results. Analyses running on the unpruned 923-taxon phylogeny also reveal the same rate shift patterns as the pruned tree, although the evolutionary rates were obviously inflated with duplicate taxa and subspecies. We are therefore confident that inclusion of additional recently published sequences and updated species richness data would not significantly impact our results.

Analyses were also repeated using global sampling fraction instead of clade-specific sampling fraction, and using different priors of expectedNumberOfshifts (1, 5, 10). As our phylogeny already had good coverage, the results from BAMM were largely insensitive to different estimates of species richness, different sampling fraction parameters, different priors of expectedNumberOfshifts, and even different ultrametric trees (pruned or not-pruned trees). Several versions of the SSE analyses were carried out for the same trait using different coding schemes. For the pollinator trapping trait, for example, different coding schemes (binary-state or multiple state; genera/species in which pollinator trapping lacks empirical field study—i.e. *Cyathocalyx*, *Drepananthus*, *Mitrella*, *Neostenanthera*, *Pseudartabotrys*, and *Trivalvaria macrophylla*—were coded as absent or present) were tried and applied to different SSE analyses. The results were invariably consistent.

BAMM

Recent simulations show that BAMM yields much weaker relationships between true and estimated diversification rates than Method-of-moment estimators [6] since it underestimates the number of distinct diversification rates present among subclades across a tree. Other simulations suggest that BAMM does not accurately estimate diversification, speciation, and/or extinction rates [7,8]. Meyer & Wiens [6] furthermore found that BAMM can yield misleading results for empirical data, with no significant relationship between rates estimated from BAMM for the same subclades in isolation and when estimated across a tree. In this study, we compared the result from BAMM using different methods, including turboMEDUSA and Method-of-moment estimators and found the most rate shifts identified by BAMM are supported by different methods as well.

BiSSE power

Although hugely popular, BiSSE has been shown to have low statistical power when the number of tips is low (fewer than 300 terminals), and when the tip-ratio bias is high (when less than 10% of species are of one character state) [9]. Since we have an extensive phylogeny and a balanced ratio of character states, type I errors seem less likely. Gamisch [10] has nevertheless shown that the power of the BiSSE method can be higher. A recent simulation study also showed that BiSSE is best suited for clades of intermediate age which have already undergone initial rapid diversification and are less likely to encounter biases which arise in older clades [11]. The use of SSE methods is suggested to still be of value, though care should be taken with the interpretation of results [12,13,14]. Although FiSSE had lower false-positive rates than SSE, FiSSE should generally be used alongside the model-based SSE approaches, and not as a replacement [14].

Supplementary Table S1. Estimation of diversification rates using method-of-moments estimator

| | Clade | stem_age | crown_age | N_species | s.r0 | s.r45 | s.r90 | c.r0 | c.r45 | c.r90 |
|---|---|---|---|---|---|---|---|---|---|---|
| family | Annonaceae | 108.64 | 93.93 | 2424 | 0.07 | 0.066 | 0.051 | 0.08 | 0.073 | 0.06 |
| subfam. | Anaxagoreoideae | 93.93 | 27.54 | 30 | 0.04 | 0.03 | 0.014 | 0.1 | 0.091 | 0.05 |
| subfam. | Ambavioideae | 87.72 | 77.61 | 56 | 0.05 | 0.039 | 0.021 | 0.04 | 0.04 | 0.02 |
| subfam. | Malmeoidaea | 86.63 | 41.94 | 823 | 0.08 | 0.071 | 0.051 | 0.14 | **0.138** | 0.1 |
| subfam. | Annonoideae | 86.63 | 82.71 | 1515 | 0.08 | 0.078 | 0.058 | 0.08 | 0.077 | 0.06 |
| tribe | Bocageeae | 82.71 | 45.58 | 63 | 0.05 | 0.043 | 0.024 | 0.08 | 0.071 | 0.04 |
| tribe | Guatterieae | 74.47 | 15.52 | 177 | 0.07 | 0.062 | 0.039 | 0.29 | **0.275** | 0.19 |
| tribe | Xylopieae | 73.25 | 62.75 | 269 | 0.08 | 0.068 | 0.045 | 0.08 | 0.075 | 0.05 |
| tribe | Duguetieae | 72.87 | 55.97 | 101 | 0.06 | 0.055 | 0.033 | 0.07 | 0.066 | 0.04 |
| tribe | Annoneae | 61.84 | 57.57 | 344 | 0.09 | **0.085** | 0.058 | 0.09 | 0.086 | 0.06 |
| tribe | Monodoreae | 53.18 | 51.44 | 86 | 0.08 | 0.073 | 0.042 | 0.07 | 0.069 | 0.04 |
| tribe | Uvariaeae | 53.18 | 35.34 | 474 | 0.12 | **0.105** | 0.073 | 0.15 | **0.148** | 0.11 |
| tribe | Piptostigmateae | 41.94 | 39.27 | 35 | 0.08 | 0.071 | 0.035 | 0.07 | 0.068 | 0.04 |
| tribe | Fenerivieae | 24.21 | 7.26 | 10 | 0.1 | 0.074 | 0.027 | 0.22 | **0.2** | 0.08 |
| tribe | Malmeeae | 24.87 | 23.55 | 180 | 0.21 | **0.185** | 0.118 | 0.19 | **0.182** | 0.12 |
| tribe | Maasieae | 24.57 | 15.4 | 6 | 0.07 | 0.054 | 0.017 | 0.07 | 0.063 | 0.02 |
| tribe | Miliuseae | 20.74 | 17.21 | 583 | 0.31 | **0.278** | 0.197 | 0.33 | **0.317** | 0.23 |
| tribe | Phoenicantheae | 21.88 | NA | 2 | 0.03 | 0.02 | 0.004 | NA | NA | NA |
| tribe | Dendrokingstonieae | 20.95 | NA | 3 | 0.05 | 0.035 | 0.009 | NA | NA | NA |
| tribe | Monocarpieae | 20.74 | NA | 4 | 0.07 | 0.047 | 0.013 | NA | NA | NA |
| genus | Annona | 47.57 | 25.91 | 170 | 0.11 | **0.095** | 0.061 | 0.17 | **0.163** | 0.11 |
| genus | Artabotrys | 62.75 | 20 | 105 | 0.07 | 0.065 | 0.039 | 0.2 | **0.187** | 0.12 |
| genus | Asimina | 41.59 | 8.53 | 17 | 0.07 | 0.055 | 0.023 | 0.25 | **0.229** | 0.11 |
| genus | Dielsiothamnus | 33.21 | NA | 1 | 0 | 0 | 0 | NA | NA | NA |
| genus | Disepalum | 41.59 | 15.55 | 9 | 0.05 | 0.041 | 0.014 | 0.1 | 0.087 | 0.03 |
| genus | Drepananthus | 15.51 | 5.93 | 26 | 0.21 | **0.174** | 0.081 | 0.43 | **0.399** | 0.2 |
| genus | Duguetia | 36.28 | 19.11 | 94 | 0.13 | **0.109** | 0.064 | 0.2 | **0.19** | 0.12 |
| genus | Goniothalamus | 52.37 | 15.47 | 134 | 0.09 | **0.082** | 0.051 | 0.27 | **0.258** | 0.17 |
| genus | Meiocarpidium | 77.61 | NA | 1 | 0 | 0 | 0 | NA | NA | NA |
| genus | Uvaria | 33.21 | 26.01 | 199 | 0.16 | **0.142** | 0.091 | 0.18 | **0.168** | 0.11 |
| genus | Xylopia | 62.75 | 26.8 | 164 | 0.08 | 0.072 | 0.045 | 0.16 | **0.156** | 0.1 |
| clade | Desmos-Dasymaschalon-Friesodielsia-Monanthotaxis | 25.98 | 16.81 | 182 | 0.2 | **0.177** | 0.114 | 0.27 | **0.255** | 0.17 |

rate in bold: higher than background rate

s.r0: rate estimated for stem group, $\varepsilon=0$

s.r45: rate estimated for stem group, $\varepsilon=0.45$

s.r90: rate estimated for stem group, $\varepsilon=0.9$

c.r0: rate estimated for crown group, $\varepsilon=0$

c.r45: rate estimated for crown group, $\varepsilon=0.45$

c.r90: rate estimated for crown group, $\varepsilon=0.9$

Supplementary Table S2. Comparison of the fit of different BiSSE models.

A. Pollination trap

| Model | Df | lnLik | AIC | ChiSq | Pr(>\|Chi\|) | ΔAIC |
|---|---|---|---|---|---|---|
| full | 6 | -2650.571 | 5313.143 | NA | NA | 1.955 |
| equal λ | 5 | -2675.890 | 5361.779 | 50.637 | 0.000 | 50.592 |
| equal μ | 5 | -2677.337 | 5364.673 | 53.530 | 0.000 | 53.485 |
| equal q | 5 | -2650.594 | 5311.188 | 0.045 | 0.832 | 0 |

B. Habit

| Model | Df | lnLik | AIC | ChiSq | Pr(>\|Chi\|) | ΔAIC |
|---|---|---|---|---|---|---|
| full | 6 | -2667.315 | 5346.631 | NA | NA | 0.002 |
| equal λ | 5 | -2676.045 | 5362.090 | 17.459 | 0.000 | 15.461 |
| equal μ | 5 | -2668.314 | 5346.629 | 1.998 | 0.158 | 0 |
| equal q | 5 | -2669.948 | 5349.896 | 5.265 | 0.022 | 3.267 |

C. Inner petal glands

| Model | Df | lnLik | AIC | ChiSq | Pr(>\|Chi\|) | ΔAIC |
|---|---|---|---|---|---|---|
| full | 6 | -2846.642 | 5705.283 | NA | NA | 0.719 |
| equal λ | 5 | -2847.282 | 5704.564 | 1.281 | 0.258 | 0 |
| equal μ | 5 | -2847.411 | 5704.821 | 1.538 | 0.215 | 0.257 |
| equal q | 5 | -2847.402 | 5704.805 | 1.522 | 0.217 | 0.241 |

D. Pollen dispersal unit

| Model | Df | lnLik | AIC | ChiSq | Pr(>\|Chi\|) | ΔAIC |
|---|---|---|---|---|---|---|
| full | 6 | -2677.745 | 5367.491 | NA | NA | 1.765 |
| equal λ | 5 | -2677.889 | 5365.777 | 0.286 | 0.593 | 0.051 |
| equal μ | 5 | -2679.720 | 5369.439 | 3.949 | 0.047 | 3.713 |
| equal q | 5 | -2677.863 | 5365.726 | 0.235 | 0.628 | 0 |

E. Anther septa

| Model | Df | lnLik | AIC | ChiSq | Pr(>\|Chi\|) | ΔAIC |
|---|---|---|---|---|---|---|
| full | 6 | -2655.945 | 5323.890 | NA | NA | 0.879 |
| equal λ | 5 | -2656.812 | 5323.624 | 1.734 | 0.188 | 0.614 |
| equal μ | 5 | -2656.505 | 5323.010 | 1.121 | 0.290 | 0 |
| equal q | 5 | -2663.565 | 5337.129 | 15.240 | 0.000 | 14.119 |

### Supplementary Table S3. Comparison of the fit of different HiSSE models.

A. Pollination trap

| Model | lnLik | AIC | ΔAIC |
|---|---|---|---|
| **HiSSE full model** | -2488.693 | 5009.386 | 0 |
| HiSSE: λ0A=λ1A, ε0A=ε1A, q's equal | -2573.69 | 5161.38 | 151.99 |
| HiSSE: q's and ε's equal | -2576.737 | 5165.473 | 156.083 |
| HiSSE: τ0A=τ1A, ε's equal, q0B1B=0, q1B0B=0, All other q's equal | -2580.625 | 5171.25 | 161.86 |
| HiSSE: ε's equal, q0B1B=0, q1B0B=0, All other q's equal | -2580.791 | 5173.583 | 164.193 |
| CID-2: q's equal | -2582.171 | 5174.343 | 164.953 |
| HiSSE: q's equal | -2578.189 | 5174.377 | 164.987 |
| HiSSE: λ0A=λ1A, ε's and q's equal | -2584.781 | 5179.561 | 170.171 |
| HiSSE: q0B1B=0, q1B0B=0, All other q's equal | -2589.916 | 5197.833 | 188.443 |
| HiSSE: τ0A=τ1A, ε0A=ε1A, q0B1B=0, q1B0B=0, All other q's equal | -2594.496 | 5202.992 | 193.602 |
| CID-2: q's equal and ε's equal | -2600.721 | 5209.441 | 200.051 |
| BiSSE model: q's equal | -2650.112 | 5310.223 | 300.833 |
| HiSSE: λ0A=λ0B, ε0A=ε0B | -2649.155 | 5312.309 | 302.919 |
| HiSSE: τ0A=τ0B, ε0A=ε0B, q0B1B=0, q1B0B=0, All other q's equal | -2663.003 | 5340.006 | 330.616 |
| HiSSE: λ0A=λ1A=λ0B, ε0A=ε1A=ε0B, q0B1B=0, q1B0B=0, All other q's equal | -2667.037 | 5344.075 | 334.685 |
| BiSSE model: q's equal, ε0=ε1 | -2668.765 | 5345.53 | 336.14 |
| HiSSE: λ0A=λ1A=λ0B, ε's and q's equal | -2668.978 | 5345.955 | 336.565 |
| HiSSE: τ0A=τ1A=τ0B, ε's equal, q0B1B=0, q1B0B=0, All other q's equal | -2669.554 | 5347.109 | 337.719 |
| HiSSE: τ0A=τ0B, ε's equal, q0B1B=0, q1B0B=0, All other q's equal | -2668.913 | 5347.827 | 338.437 |
| HiSSE: λ0A=λ0B, ε's and q's equal | -2668.955 | 5347.909 | 338.519 |
| BiSSE model: ε0=ε1 | -2668.992 | 5347.985 | 338.595 |
| HiSSE: τ0A=τ1A=τ0B, ε0A=ε1A=ε1B, q's equal | -2669.866 | 5349.733 | 340.343 |
| CID-4: ε's and q's equal | -2669.34 | 5350.68 | 341.29 |
| BiSSE model: All fτee | -2677.996 | 5367.991 | 358.601 |
| CID-4: q's equal | -2679.239 | 5376.479 | 367.089 |

B. Habit

| Model | lnLik | AIC | ΔAIC |
|---|---|---|---|
| HiSSE full model | -2497.284 | 5026.568 | 0 |
| HiSSE: τ0A=τ1A, ε0A=ε1A, q0B1B=0, q1B0B=0, All other q's equal | -2572.96 | 5159.92 | 133.352 |
| HiSSE: λ0A=λ1A, ε0A=ε1A, q's equal | -2576.052 | 5166.105 | 139.537 |
| HiSSE: λ0A=λ1A, ε's and q's equal | -2580.783 | 5171.566 | 144.998 |
| HiSSE: ε's equal, q0B1B=0, q1B0B=0, All other q's equal | -2581.434 | 5174.868 | 148.3 |
| CID-2: q's equal | -2585.277 | 5180.553 | 153.985 |
| HiSSE: q's and ε's equal | -2584.709 | 5181.417 | 154.849 |
| HiSSE: q0B1B=0, q1B0B=0, All other q's equal | -2583.685 | 5185.37 | 158.802 |
| HiSSE: q's equal | -2589.548 | 5197.095 | 170.527 |
| CID-2: q's equal and ε's equal | -2600.618 | 5209.236 | 182.668 |
| HiSSE: τ0A=τ1A, ε's equal, q0B1B=0, q1B0B=0, All other q's equal | -2606.975 | 5223.95 | 197.382 |
| HiSSE: τ0A=τ0B, ε0A=ε0B, q0B1B=0, q1B0B=0, All other q's equal | -2658.751 | 5331.503 | 304.935 |
| HiSSE: λ0A=λ1A=λ0B, ε0A=ε1A=ε0B, q0B1B=0, q1B0B=0, All other q's equal | -2660.847 | 5331.694 | 305.126 |
| HiSSE: τ0A=τ0B, ε's equal, q0B1B=0, q1B0B=0, All other q's equal | -2664.687 | 5339.374 | 312.806 |

| | | | |
|---|---|---|---|
| CID-4: ε's and q's equal | -2664.893 | 5341.785 | 315.217 |
| BiSSE model: q's equal, ε0=ε1 | -2671.251 | 5350.502 | 323.934 |
| HiSSE: λ0A=λ1A=λ0B, ε's and q's equal | -2671.389 | 5350.777 | 324.209 |
| HiSSE: λ0A=λ0B, ε0A=ε0B | -2668.772 | 5351.544 | 324.976 |
| HiSSE: λ0A=λ0B, ε's and q's equal | -2670.91 | 5351.82 | 325.252 |
| BiSSE model: q's equal | -2670.969 | 5351.937 | 325.369 |
| BiSSE model: All fτee | -2670.013 | 5352.026 | 325.458 |
| HiSSE: τ0A=τ1A=τ0B, ε's equal, q0B1B=0, q1B0B=0, All other q's equal | -2674.687 | 5357.374 | 330.806 |
| HiSSE: τ0A=τ1A=τ0B, ε0A=ε1A=ε1B, q's equal | -2675.067 | 5360.134 | 333.566 |
| CID-4: q's equal | -2671.569 | 5361.139 | 334.571 |
| BiSSE model: ε0=ε1 | -2678.648 | 5367.297 | 340.729 |

C. Inner petal glands

| Model | lnLik | AIC | ΔAIC |
|---|---|---|---|
| HiSSE full model | -2656.307 | 5344.615 | 0 |
| HiSSE: q0B1B=0, q1B0B=0, All other q's equal | -2766.833 | 5551.665 | 207.05 |
| HiSSE: τ0A=τ1A, ε0A=ε1A, q's equal | -2779.504 | 5573.008 | 228.393 |
| HiSSE: q's and ε's equal | -2782.431 | 5576.863 | 232.248 |
| HiSSE: q's equal | -2781.397 | 5580.793 | 236.178 |
| CID-2: q's equal | -2787.939 | 5585.878 | 241.263 |
| CID-2: q's equal and ε's equal | -2796.145 | 5600.29 | 255.675 |
| HiSSE: τ0A=τ1A, ε's and q's equal | -2795.868 | 5601.737 | 257.122 |
| HiSSE: ε's equal, q0B1B=0, q1B0B=0, All other q's equal | -2802.675 | 5617.35 | 272.735 |
| HiSSE: τ0A=τ1A, ε0A=ε1A, q0B1B=0, q1B0B=0, All other q's equal | -2806.766 | 5627.532 | 282.917 |
| HiSSE: τ0A=τ1A, ε's equal, q0B1B=0, q1B0B=0, All other q's equal | -2823.567 | 5657.135 | 312.52 |
| BiSSE model: ε0=ε1 | -2827.696 | 5665.391 | 320.776 |
| HiSSE: τ0A=τ0B, ε0A=ε0B, q0B1B=0, q1B0B=0, All other q's equal | -2826.849 | 5667.699 | 323.084 |
| HiSSE: τ0A=τ1A=τ0B, ε's equal, q0B1B=0, q1B0B=0, All other q's equal | -2834.491 | 5676.983 | 332.368 |
| HiSSE: τ0A=τ1A=τ0B, ε0A=ε1A=ε0B, q0B1B=0, q1B0B=0, All other q's equal | -2834.374 | 5678.749 | 334.134 |
| HiSSE: τ0A=τ0B, ε's equal, q0B1B=0, q1B0B=0, All other q's equal | -2836.252 | 5682.504 | 337.889 |
| HiSSE: τ0A=τ0B, ε0A=ε0B | -2835.869 | 5685.738 | 341.123 |
| HiSSE: τ0A=τ1A=τ0B, ε0A=ε1A=ε1B, q's equal | -2843.73 | 5697.459 | 352.844 |
| CID-4: q's equal | -2841.791 | 5701.581 | 356.966 |
| BiSSE model: All fτee | -2848.419 | 5708.837 | 364.222 |
| BiSSE model: q's equal | -2850.79 | 5711.58 | 366.965 |
| HiSSE: τ0A=τ0B, ε's and q's equal | -2857.511 | 5725.022 | 380.407 |
| HiSSE: τ0A=τ1A=τ0B, ε's and q's equal | -2862.98 | 5733.961 | 389.346 |
| BiSSE model: q's equal, ε0=ε1 | -2875.592 | 5759.185 | 414.57 |
| CID-4: ε's and q's equal | -2887.177 | 5786.354 | 441.739 |

D. Pollen dispersal unit

| Model | lnLik | AIC | ΔAIC |
|---|---|---|---|
| HiSSE full model | -2538.647 | 5109.294 | 0 |
| HiSSE: ε's equal, q0B1B=0, q1B0B=0, All other q's equal | -2601.966 | 5215.932 | 106.638 |

| | | | |
|---|---|---|---|
| HiSSE: τ0A=τ1A, ε's and q's equal | -2612.056 | 5234.113 | 124.819 |
| HiSSE: q's and ε's equal | -2613.622 | 5239.245 | 129.951 |
| HiSSE: q0B1B=0, q1B0B=0, All other q's equal | -2612.826 | 5243.652 | 134.358 |
| CID-2: q's equal | -2617.286 | 5244.573 | 135.279 |
| HiSSE: τ0A=τ1A, ε0A=ε1A, q0B1B=0, q1B0B=0, All other q's equal | -2621.751 | 5257.502 | 148.208 |
| HiSSE: q's equal | -2621.54 | 5261.079 | 151.785 |
| HiSSE: τ0A=τ1A, ε0A=ε1A, q's equal | -2625.112 | 5264.224 | 154.93 |
| CID-2: q's equal and ε's equal | -2638.282 | 5284.565 | 175.271 |
| HiSSE: τ0A=τ1A, ε's equal, q0B1B=0, q1B0B=0, All other q's equal | -2649.119 | 5308.238 | 198.944 |
| HiSSE: τ0A=τ0B, ε's and q's equal | -2657.275 | 5324.549 | 215.255 |
| HiSSE: τ0A=τ0B, ε0A=ε0B | -2658.986 | 5331.971 | 222.677 |
| CID-4: q's equal | -2659.502 | 5337.004 | 227.71 |
| HiSSE: τ0A=τ1A=τ0B, ε0A=ε1A=ε1B, q's equal | -2667.646 | 5345.293 | 235.999 |
| HiSSE: τ0A=τ0B, ε0A=ε0B, q0B1B=0, q1B0B=0, All other q's equal | -2670.818 | 5355.635 | 246.341 |
| BiSSE model: q's equal | -2680.061 | 5370.122 | 260.828 |
| BiSSE model: All fτee | -2679.666 | 5371.333 | 262.039 |
| HiSSE: τ0A=τ0B, ε's equal, q0B1B=0, q1B0B=0, All other q's equal | -2681.288 | 5372.576 | 263.282 |
| HiSSE: τ0A=τ1A=τ0B, ε0A=ε1A=ε0B, q0B1B=0, q1B0B=0, All other q's equal | -2688.611 | 5387.223 | 277.929 |
| BiSSE model: q's equal, ε0=ε1 | -2689.782 | 5387.564 | 278.27 |
| BiSSE model: ε0=ε1 | -2688.935 | 5387.87 | 278.576 |
| HiSSE: τ0A=τ1A=τ0B, ε's equal, q0B1B=0, q1B0B=0, All other q's equal | -2693.13 | 5394.26 | 284.966 |
| HiSSE: τ0A=τ1A=τ0B, ε's and q's equal | -2698.985 | 5405.97 | 296.676 |
| CID-4: ε's and q's equal | -2716.057 | 5444.114 | 334.82 |

E. Anther septa

| Model | lnLik | AIC | ΔAIC |
|---|---|---|---|
| HiSSE full model | -2503.619 | 5039.238 | 0 |
| HiSSE: q0B1B=0, q1B0B=0, All other q's equal | -2562.071 | 5142.143 | 102.905 |
| HiSSE: ε's equal, q0B1B=0, q1B0B=0, All other q's equal | -2572.842 | 5157.685 | 118.447 |
| HiSSE: τ0A=τ1A, ε0A=ε1A, q0B1B=0, q1B0B=0, All other q's equal | -2571.884 | 5157.768 | 118.53 |
| HiSSE: τ0A=τ1A, ε0A=ε1A, q's equal | -2573.826 | 5161.651 | 122.413 |
| CID-2: q's equal | -2588.44 | 5186.88 | 147.642 |
| HiSSE: τ0A=τ1A, ε's and q's equal | -2593.465 | 5196.93 | 157.692 |
| HiSSE: q's and ε's equal | -2593.351 | 5198.703 | 159.465 |
| HiSSE: q's equal | -2596.073 | 5210.146 | 170.908 |
| CID-2: q's equal and ε's equal | -2606.278 | 5220.556 | 181.318 |
| HiSSE: τ0A=τ1A, ε's equal, q0B1B=0, q1B0B=0, All other q's equal | -2610.864 | 5231.728 | 192.49 |
| HiSSE: τ0A=τ0B, ε0A=ε0B | -2641.923 | 5297.845 | 258.607 |
| HiSSE: τ0A=τ0B, ε's equal, q0B1B=0, q1B0B=0, All other q's equal | -2647.708 | 5305.415 | 266.177 |
| HiSSE: τ0A=τ0B, ε's and q's equal | -2653.598 | 5317.196 | 277.958 |
| HiSSE: τ0A=τ0B, ε0A=ε0B, q0B1B=0, q1B0B=0, All other q's equal | -2655.619 | 5325.238 | 286 |
| BiSSE model: All fτee | -2658.332 | 5328.665 | 289.427 |
| CID-4: q's equal | -2655.635 | 5329.269 | 290.031 |
| HiSSE: τ0A=τ1A=τ0B, ε's and q's equal | -2664.87 | 5337.74 | 298.502 |

| | | | |
|---|---|---|---|
| HiSSE: τ0A=τ1A=τ0B, ε0A=ε1A=ε1B, q's equal | -2664.075 | 5338.151 | 298.913 |
| HiSSE: τ0A=τ1A=τ0B, ε's equal, q0B1B=0, q1B0B=0, All other q's equal | -2665.653 | 5339.306 | 300.068 |
| HiSSE: τ0A=τ1A=τ0B, ε0A=ε1A=ε0B, q0B1B=0, q1B0B=0, All other q's equal | -2664.765 | 5339.53 | 300.292 |
| BiSSE model: q's equal | -2665.535 | 5341.07 | 301.832 |
| BiSSE model: q's equal, ε0=ε1 | -2667.282 | 5342.563 | 303.325 |
| BiSSE model: ε0=ε1 | -2667.43 | 5344.859 | 305.621 |
| CID-4: ε's and q's equal | -2671.888 | 5355.776 | 316.538 |

Supplementary Table S4. Comparison of the fit of different MuSSE models.

A. Pollination trap

| Models | Df | lnLik | AIC | ChiSq | Pr(>\|Chi\|) | ΔAIC |
|---|---|---|---|---|---|---|
| Model0 (all free) | 12 | -2675.619 | 5375.238 | NA | NA | 57.621 |
| Model1 ( μ and q free) | 10 | -2677.600 | 5375.201 | 3.962 | 0.138 | 57.583 |
| Model2 (q free) | 8 | -2671.966 | 5359.931 | -7.307 | 1.000 | 42.314 |
| Model3 (all constrained) | 3 | -2673.209 | 5352.418 | -4.820 | 1.000 | 34.801 |
| Model4 (λ free) | 5 | -2668.295 | 5346.590 | -14.649 | 1.000 | 28.972 |
| Model5 (μ free) | 5 | -2661.462 | 5332.923 | -28.315 | 1.000 | 15.306 |
| Model6 (λ and μ free) | 7 | -2651.809 | 5317.617 | -47.621 | 1.000 | 0.000 |
| model7 (λ and q free) | 10 | -2680.983 | 5381.967 | 10.728 | 0.005 | 64.349 |

B. Floral sexuality at population level

| Models | Df | lnLik | AIC | ChiSq | Pr(>\|Chi\|) | ΔAIC |
|---|---|---|---|---|---|---|
| Model0 (all free) | 12 | -2726.580 | 5477.160 | NA | NA | 0.887 |
| Model1 ( μ and q free) | 10 | -2741.549 | 5503.098 | 29.938 | 0.000 | 26.825 |
| Model2 (q free) | 8 | -2768.895 | 5553.789 | 84.630 | 0.000 | 77.517 |
| Model3 (all constrained) | 3 | -2821.838 | 5649.676 | 190.516 | 0.000 | 173.404 |
| Model4 (λ free) | 5 | -2813.499 | 5636.999 | 173.839 | 0.000 | 160.726 |
| Model5 (μ free) | 5 | -2797.469 | 5604.938 | 141.779 | 0.000 | 128.666 |
| Model6 (λ and μ free) | 7 | -2787.080 | 5588.160 | 121.000 | 0.000 | 111.888 |
| model7 (λ and q free) | 10 | -2728.136 | 5476.272 | 3.113 | 0.211 | 0 |

C. Pollen dispersal unit

| Models | Df | lnLik | AIC | ChiSq | Pr(>\|Chi\|) | ΔAIC |
|---|---|---|---|---|---|---|
| Model0 (all free) | 12 | -2675.657 | 5375.314 | NA | NA | 2.192 |
| Model1 ( μ and q free) | 10 | -2689.123 | 5398.246 | 26.932 | 0.000 | 25.124 |
| Model2 (q free) | 8 | -2730.600 | 5477.199 | 109.885 | 0.000 | 104.077 |
| Model3 (all constrained) | 3 | -2735.865 | 5477.729 | 120.415 | 0.000 | 104.607 |
| Model4 (λ free) | 5 | -2696.708 | 5403.416 | 42.102 | 0.000 | 30.294 |
| Model5 (μ free) | 5 | -2697.071 | 5404.143 | 42.829 | 0.000 | 31.021 |
| Model6 (λ and μ free) | 7 | -2690.741 | 5395.482 | 30.168 | 0.000 | 22.360 |
| model7 (λ and q free) | 10 | -2676.561 | 5373.122 | 1.808 | 0.405 | 0 |

D. seed dispersal unit

| Models | Df | lnLik | AIC | ChiSq | Pr(>\|Chi\|) | ΔAIC |
|---|---|---|---|---|---|---|
| Model0 (all free) | 20 | -2656.755 | 5353.509 | NA | NA | 47.578 |
| Model1 ( μ and q free) | 17 | -2638.730 | 5311.460 | -36.049 | 1.000 | 5.529 |
| Model2 (q free) | 14 | -2660.366 | 5348.733 | 7.224 | 0.301 | 42.802 |
| Model3 (all constrained) | 3 | -2686.291 | 5378.583 | 59.073 | 0.000 | 72.651 |
| Model4 (λ free) | 6 | -2677.825 | 5367.650 | 42.140 | 0.000 | 61.718 |
| Model5 (μ free) | 6 | -2677.433 | 5366.866 | 41.356 | 0.000 | 60.934 |
| Model6 (λ and μ free) | 9 | -2670.442 | 5358.884 | 27.375 | 0.004 | 52.953 |
| model7 (λ and q free) | 17 | -2635.966 | 5305.931 | -41.578 | 1.000 | 0 |

Supplementary Table S5. Species richness data and sampling fraction for BAMM and turboMEDUSA analyses.

| Genus | Sampled species | Known diversity | Sampling fraction |
|---|---|---|---|
| *Afroguatteria* | 2 | 3 | 0.67 |
| *Alphonsea* | 9 | 29 | 0.31 |
| *Ambavia* | 1 | 2 | 0.5 |
| *Anaxagorea* | 6 | 30 | 0.2 |
| *Annickia* | 3 | 8 | 0.38 |
| *Annona* | 14 | 170 | 0.08 |
| *Anodidium* | 1 | 5 | 0.2 |
| *Artabotrys* | 16 | 105 | 0.15 |
| *Asimina* | 10 | 17 | 0.59 |
| *Asteranthe* | 1 | 3 | 0.33 |
| *Bocagea* | 2 | 2 | 1 |
| *Bocageopsis* | 3 | 4 | 0.75 |
| *Brieya* | 1 | 2 | 0.5 |
| *Cananga* | 2 | 2 | 1 |
| *Cardiopetalum* | 2 | 3 | 0.67 |
| *Cleistochlamys* | 1 | 1 | 1 |
| *Cleistopholis* | 1 | 4 | 0.25 |
| *Cremastosperma* | 21 | 29 | 0.72 |
| *Cyathocalyx* | 7 | 9 | 0.78 |
| *Cymbopetalum* | 2 | 27 | 0.07 |
| *Dasymaschalon* | 18 | 27 | 0.67 |
| *Dendrokingstonia* | 2 | 3 | 0.67 |
| *DesmopStena* | 18 | 28 | 0.64 |
| *Desmos* | 8 | 22 | 0.36 |
| *Diclinanona* | 1 | 3 | 0.33 |
| *Dielsiothamnus* | 1 | 1 | 1 |
| *Disepalum* | 8 | 9 | 0.89 |
| *Drepananthus* | 16 | 26 | 0.62 |
| *Duckeanthus* | 1 | 1 | 1 |
| *Duguetia* | 32 | 94 | 0.34 |
| *Ephedranthus* | 2 | 7 | 0.29 |
| *Fenerivia* | 10 | 10 | 1 |
| *Fissistigma* | 3 | 59 | 0.05 |
| *Friesodielsia* | 15 | 39 | 0.38 |
| *Fusaea* | 2 | 2 | 1 |
| *Goniothalamus* | 65 | 134 | 0.49 |
| *Greenwayodendron* | 2 | 2 | 1 |
| *Guatteria* | 72 | 177 | 0.41 |
| *Hexalobus* | 2 | 5 | 0.4 |
| *Hornschuchia* | 1 | 10 | 0.1 |
| *Huberantha* | 11 | 35 | 0.31 |
| *Isolona* | 15 | 20 | 0.75 |
| *Letestudoxa* | 2 | 3 | 0.67 |
| *Lettowianthus* | 1 | 1 | 1 |
| *Maasia* | 5 | 6 | 0.83 |
| *Malmea* | 3 | 7 | 0.43 |
| *Marsypopetalum* | 4 | 6 | 0.67 |
| *Meiocarpidium* | 1 | 1 | 1 |
| *Meiogyne* | 21 | 29 | 0.72 |
| *Mezzettia* | 1 | 4 | 0.25 |
| *Miliusa* | 18 | 60 | 0.3 |
| *Mischogyne* | 1 | 2 | 0.5 |
| *Mitrella* | 1 | 9 | 0.11 |
| *Mitrephora* | 6 | 49 | 0.12 |
| *Mkilua* | 1 | 1 | 1 |

| Genus | Sampled species | Known diversity | Sampling fraction |
|---|---|---|---|
| *Monanthotaxis* | 36 | 94 | 0.38 |
| *Monocarpia* | 2 | 4 | 0.5 |
| *Monocyclanthus* | 1 | 1 | 1 |
| *Monodora* | 13 | 14 | 0.93 |
| *Monoon* | 21 | 66 | 0.32 |
| *Mosannona* | 6 | 14 | 0.43 |
| *Mwasumbia* | 1 | 1 | 1 |
| *Neostenanthera* | 2 | 6 | 0.33 |
| *Neouvaria* | 3 | 7 | 0.43 |
| *Onychopetalum* | 1 | 2 | 0.5 |
| *Ophrypetalum* | 1 | 1 | 1 |
| *Orophea* | 7 | 57 | 0.12 |
| *Phaeanthus* | 2 | 8 | 0.25 |
| *Phoenicanthus* | 1 | 2 | 0.5 |
| *Piptostigma* | 3 | 13 | 0.23 |
| *Platymitra* | 1 | 2 | 0.5 |
| *Polyalthia* | 27 | 91 | 0.3 |
| *Polyalthiopsis* | 1 | 1 | 1 |
| *Polyceratocarpus* | 4 | 8 | 0.5 |
| *Popowia* | 3 | 29 | 0.1 |
| *Porcelia* | 1 | 8 | 0.13 |
| *Pseudartabotrys* | 1 | 1 | 1 |
| *Pseudoxandra* | 4 | 24 | 0.17 |
| *Pseuduvaria* | 52 | 54 | 0.96 |
| *PseuOxanKla* | 20 | 45 | 0.44 |
| *Pyramidanthe* | 1 | 1 | 1 |
| *Ruizodendron* | 1 | 1 | 1 |
| *Sageraea* | 1 | 9 | 0.11 |
| *Sanrafaelia* | 1 | 1 | 1 |
| *Sapranthus* | 3 | 6 | 0.5 |
| *Sirdavidia* | 1 | 1 | 1 |
| *Sphaerocoryne* | 4 | 4 | 1 |
| *Stelechocarpus* | 3 | 4 | 0.75 |
| *Tetrameranthus* | 2 | 7 | 0.29 |
| *Toussaintia* | 1 | 4 | 0.25 |
| *Tridimeris* | 2 | 2 | 1 |
| *Trigynaea* | 2 | 9 | 0.22 |
| *Trivalvaria* | 2 | 8 | 0.25 |
| *Unonopsis* | 2 | 48 | 0.04 |
| *Uvaria* | 59 | 199 | 0.3 |
| *Uvariastrum* | 3 | 5 | 0.6 |
| *Uvariodendron* | 3 | 15 | 0.2 |
| *Uvariopsis* | 4 | 19 | 0.21 |
| *Wangia* | 2 | 2 | 1 |
| *Wuodendron* | 1 | 1 | 1 |
| *Xylopia* | 48 | 164 | 0.29 |

Genus-level clades:
*DesmopStena: Stenanona* and *Desmopsis*
*PseuOxanKla: Klarobelia-Oxandra-Pseudephedranthus-Pseudomalmea*
*Friesodielsia: Friesodielsia-Schefferomitra*

Supplementary Table S6. Species richness, clade age, and net diversification rate for 101 genus-level clades used in PGLS analysis.

| tips | stem_age | N_species | stem_age_ln | N_species_ln | s.r45 | s.r0 | s.r90 |
|---|---|---|---|---|---|---|---|
| Cleistochlamys | 16.790 | 1.000 | 2.821 | 0.000 | 0.000 | 0.000 | 0.000 |
| Dielsiothamnus | 33.210 | 1.000 | 3.503 | 0.000 | 0.000 | 0.000 | 0.000 |
| Duckeanthus | 21.660 | 1.000 | 3.075 | 0.000 | 0.000 | 0.000 | 0.000 |
| Lettowianthus | 28.230 | 1.000 | 3.340 | 0.000 | 0.000 | 0.000 | 0.000 |
| Meiocarpidium | 77.610 | 1.000 | 4.352 | 0.000 | 0.000 | 0.000 | 0.000 |
| Mkilua | 45.580 | 1.000 | 3.819 | 0.000 | 0.000 | 0.000 | 0.000 |
| Monocyclanthus | 9.960 | 1.000 | 2.299 | 0.000 | 0.000 | 0.000 | 0.000 |
| Mwasumbia | 18.510 | 1.000 | 2.918 | 0.000 | 0.000 | 0.000 | 0.000 |
| Ophrypetalum | 17.670 | 1.000 | 2.872 | 0.000 | 0.000 | 0.000 | 0.000 |
| Polyalthiopsis | 15.090 | 1.000 | 2.714 | 0.000 | 0.000 | 0.000 | 0.000 |
| Pseudartabotrys | 21.680 | 1.000 | 3.076 | 0.000 | 0.000 | 0.000 | 0.000 |
| Pyramidanthe | 9.190 | 1.000 | 2.218 | 0.000 | 0.000 | 0.000 | 0.000 |
| Ruizodendron | 16.040 | 1.000 | 2.775 | 0.000 | 0.000 | 0.000 | 0.000 |
| Sanrafaelia | 17.670 | 1.000 | 2.872 | 0.000 | 0.000 | 0.000 | 0.000 |
| Sirdavidia | 18.510 | 1.000 | 2.918 | 0.000 | 0.000 | 0.000 | 0.000 |
| Wuodendron | 14.840 | 1.000 | 2.697 | 0.000 | 0.000 | 0.000 | 0.000 |
| Greenwayodendron | 30.980 | 2.000 | 3.433 | 0.693 | 0.014 | 0.022 | 0.003 |
| Diclinanona | 47.610 | 3.000 | 3.863 | 1.099 | 0.016 | 0.023 | 0.004 |
| Brieya | 24.410 | 2.000 | 3.195 | 0.693 | 0.018 | 0.028 | 0.004 |
| Phoenicanthus | 24.210 | 2.000 | 3.187 | 0.693 | 0.018 | 0.029 | 0.004 |
| Fusaea | 21.660 | 2.000 | 3.075 | 0.693 | 0.020 | 0.032 | 0.004 |
| Cananga | 21.480 | 2.000 | 3.067 | 0.693 | 0.020 | 0.032 | 0.004 |
| Mischogyne | 19.150 | 2.000 | 2.952 | 0.693 | 0.023 | 0.036 | 0.005 |
| Anodidium | 46.970 | 5.000 | 3.850 | 1.609 | 0.025 | 0.034 | 0.007 |
| Cardiopetalum | 28.350 | 3.000 | 3.345 | 1.099 | 0.026 | 0.039 | 0.006 |
| Neostenanthera | 46.970 | 6.000 | 3.850 | 1.792 | 0.028 | 0.038 | 0.009 |
| Ambavia | 15.400 | 2.000 | 2.734 | 0.693 | 0.028 | 0.045 | 0.006 |
| Bocagea | 14.960 | 2.000 | 2.705 | 0.693 | 0.029 | 0.046 | 0.006 |
| Anaxagorea | 93.930 | 30.000 | 4.543 | 3.401 | 0.030 | 0.036 | 0.014 |
| Asteranthe | 23.380 | 3.000 | 3.152 | 1.099 | 0.032 | 0.047 | 0.008 |
| Platymitra | 13.400 | 2.000 | 2.595 | 0.693 | 0.033 | 0.052 | 0.007 |
| Wangia | 13.190 | 2.000 | 2.579 | 0.693 | 0.033 | 0.053 | 0.007 |
| Dendrokingstonia | 21.880 | 3.000 | 3.086 | 1.099 | 0.034 | 0.050 | 0.008 |
| Letestudoxa | 21.680 | 3.000 | 3.076 | 1.099 | 0.034 | 0.051 | 0.008 |
| Tridimeris | 11.840 | 2.000 | 2.471 | 0.693 | 0.037 | 0.059 | 0.008 |
| Cleistopholis | 24.830 | 4.000 | 3.212 | 1.386 | 0.039 | 0.056 | 0.011 |
| Annickia | 39.270 | 8.000 | 3.670 | 2.079 | 0.040 | 0.053 | 0.014 |
| Disepalum | 41.590 | 9.000 | 3.728 | 2.197 | 0.041 | 0.053 | 0.014 |
| Tetrameranthus | 31.950 | 7.000 | 3.464 | 1.946 | 0.046 | 0.061 | 0.015 |

| | | | | | | | |
|---|---|---|---|---|---|---|---|
| Monocarpia | 20.950 | 4.000 | 3.042 | 1.386 | 0.047 | 0.066 | 0.013 |
| Porcelia | 30.960 | 8.000 | 3.433 | 2.079 | 0.051 | 0.067 | 0.017 |
| Onychopetalum | 8.510 | 2.000 | 2.141 | 0.693 | 0.051 | 0.081 | 0.011 |
| Maasia | 24.570 | 6.000 | 3.202 | 1.792 | 0.054 | 0.073 | 0.017 |
| Asimina | 41.590 | 17.000 | 3.728 | 2.833 | 0.055 | 0.068 | 0.023 |
| Sphaerocoryne | 16.790 | 4.000 | 2.821 | 1.386 | 0.058 | 0.083 | 0.016 |
| Guatteria | 74.470 | 177.000 | 4.310 | 5.176 | 0.062 | 0.070 | 0.039 |
| Mezzettia | 15.400 | 4.000 | 2.734 | 1.386 | 0.063 | 0.090 | 0.017 |
| Bocageopsis | 15.300 | 4.000 | 2.728 | 1.386 | 0.064 | 0.091 | 0.017 |
| Hexalobus | 18.040 | 5.000 | 2.893 | 1.609 | 0.064 | 0.089 | 0.019 |
| Uvariastrum | 18.040 | 5.000 | 2.893 | 1.609 | 0.064 | 0.089 | 0.019 |
| Artabotrys | 62.750 | 105.000 | 4.139 | 4.654 | 0.065 | 0.074 | 0.039 |
| Malmea | 21.260 | 7.000 | 3.057 | 1.946 | 0.069 | 0.092 | 0.022 |
| Xylopia | 62.750 | 164.000 | 4.139 | 5.100 | 0.072 | 0.081 | 0.045 |
| Fenerivia | 24.210 | 10.000 | 3.187 | 2.303 | 0.074 | 0.095 | 0.027 |
| Goniothalamus | 52.370 | 134.000 | 3.958 | 4.898 | 0.082 | 0.094 | 0.051 |
| Marsypopetalum | 15.060 | 6.000 | 2.712 | 1.792 | 0.088 | 0.119 | 0.027 |
| Afroguatteria | 8.400 | 3.000 | 2.128 | 1.099 | 0.088 | 0.131 | 0.022 |
| Hornschuchia | 19.540 | 10.000 | 2.972 | 2.303 | 0.091 | 0.118 | 0.033 |
| Ephedranthus | 15.930 | 7.000 | 2.768 | 1.946 | 0.092 | 0.122 | 0.030 |
| Stelechocarpus | 10.450 | 4.000 | 2.347 | 1.386 | 0.093 | 0.133 | 0.025 |
| Phaeanthus | 16.910 | 8.000 | 2.828 | 2.079 | 0.093 | 0.123 | 0.031 |
| Polyceratocarpus | 16.720 | 8.000 | 2.817 | 2.079 | 0.094 | 0.124 | 0.032 |
| Annona | 47.570 | 170.000 | 3.862 | 5.136 | 0.095 | 0.108 | 0.061 |
| Cymbopetalum | 28.350 | 27.000 | 3.345 | 3.296 | 0.096 | 0.116 | 0.045 |
| Monodora | 21.710 | 14.000 | 3.078 | 2.639 | 0.097 | 0.122 | 0.038 |
| Piptostigma | 20.010 | 13.000 | 2.996 | 2.565 | 0.101 | 0.128 | 0.039 |
| Trivalvaria | 15.060 | 8.000 | 2.712 | 2.079 | 0.105 | 0.138 | 0.035 |
| Cyathocalyx | 15.510 | 9.000 | 2.741 | 2.197 | 0.109 | 0.142 | 0.038 |
| Mosannona | 19.280 | 14.000 | 2.959 | 2.639 | 0.109 | 0.137 | 0.043 |
| Duguetia | 36.280 | 94.000 | 3.591 | 4.543 | 0.109 | 0.125 | 0.064 |
| Sapranthus | 11.840 | 6.000 | 2.471 | 1.792 | 0.112 | 0.151 | 0.034 |
| Isolona | 21.710 | 20.000 | 3.078 | 2.996 | 0.112 | 0.138 | 0.049 |
| Trigynaea | 14.960 | 9.000 | 2.705 | 2.197 | 0.113 | 0.147 | 0.039 |
| Toussaintia | 8.400 | 4.000 | 2.128 | 1.386 | 0.116 | 0.165 | 0.031 |
| Neouvaria | 11.380 | 7.000 | 2.432 | 1.946 | 0.128 | 0.171 | 0.041 |
| Uvaria | 33.210 | 199.000 | 3.503 | 5.293 | 0.142 | 0.159 | 0.091 |
| Uvariodendron | 14.130 | 15.000 | 2.648 | 2.708 | 0.153 | 0.192 | 0.062 |
| Sageraea | 10.450 | 9.000 | 2.347 | 2.197 | 0.161 | 0.210 | 0.056 |
| Fissistigma | 21.590 | 59.000 | 3.072 | 4.078 | 0.162 | 0.189 | 0.089 |
| Desmos | 14.690 | 22.000 | 2.687 | 3.091 | 0.172 | 0.210 | 0.077 |
| Drepananthus | 15.510 | 26.000 | 2.741 | 3.258 | 0.174 | 0.210 | 0.081 |
| Meiogyne | 16.020 | 29.000 | 2.774 | 3.367 | 0.175 | 0.210 | 0.083 |

| | | | | | | | |
|---|---|---|---|---|---|---|---|
| Huberantha | 16.910 | 35.000 | 2.828 | 3.555 | 0.176 | 0.210 | 0.088 |
| Pseudoxandra | 14.820 | 24.000 | 2.696 | 3.178 | 0.176 | 0.214 | 0.081 |
| Mitrella | 9.190 | 9.000 | 2.218 | 2.197 | 0.184 | 0.239 | 0.064 |
| Dasymaschalon | 14.750 | 27.000 | 2.691 | 3.296 | 0.185 | 0.223 | 0.087 |
| Cremastosperma | 14.820 | 29.000 | 2.696 | 3.367 | 0.189 | 0.227 | 0.090 |
| Popowia | 14.340 | 29.000 | 2.663 | 3.367 | 0.195 | 0.235 | 0.093 |
| Mitrephora | 16.740 | 49.000 | 2.818 | 3.892 | 0.198 | 0.232 | 0.105 |
| Orophea | 17.210 | 57.000 | 2.845 | 4.043 | 0.201 | 0.235 | 0.110 |
| DesmopStena | 13.670 | 28.000 | 2.615 | 3.332 | 0.202 | 0.244 | 0.096 |
| PseuOxanKla | 15.930 | 45.000 | 2.768 | 3.807 | 0.203 | 0.239 | 0.106 |
| Alphonsea | 13.400 | 29.000 | 2.595 | 3.367 | 0.209 | 0.251 | 0.100 |
| Friesodielsia | 14.690 | 39.000 | 2.687 | 3.664 | 0.210 | 0.249 | 0.107 |
| Pseuduvaria | 16.170 | 54.000 | 2.783 | 3.989 | 0.211 | 0.247 | 0.114 |
| Unonopsis | 15.300 | 48.000 | 2.728 | 3.871 | 0.215 | 0.253 | 0.114 |
| Miliusa | 15.090 | 60.000 | 2.714 | 4.094 | 0.233 | 0.271 | 0.128 |
| Monanthotaxis | 16.810 | 94.000 | 2.822 | 4.543 | 0.235 | 0.270 | 0.139 |
| Uvariopsis | 9.960 | 19.000 | 2.299 | 2.944 | 0.240 | 0.296 | 0.103 |
| Polyalthia | 14.340 | 91.000 | 2.663 | 4.511 | 0.273 | 0.315 | 0.161 |
| Monoon | 11.380 | 66.000 | 2.432 | 4.190 | 0.317 | 0.368 | 0.177 |

s.r0: rate estimated for stem group, ε=0

s.r45: rate estimated for stem group, ε=0.45

s.r90: rate estimated for stem group, ε=0.9

stem_age_ln: ln-transformed stem age

N_species_ln: ln-transformed  richness

Supplementary Fig. S1. RaxML tree with 923 terminals.

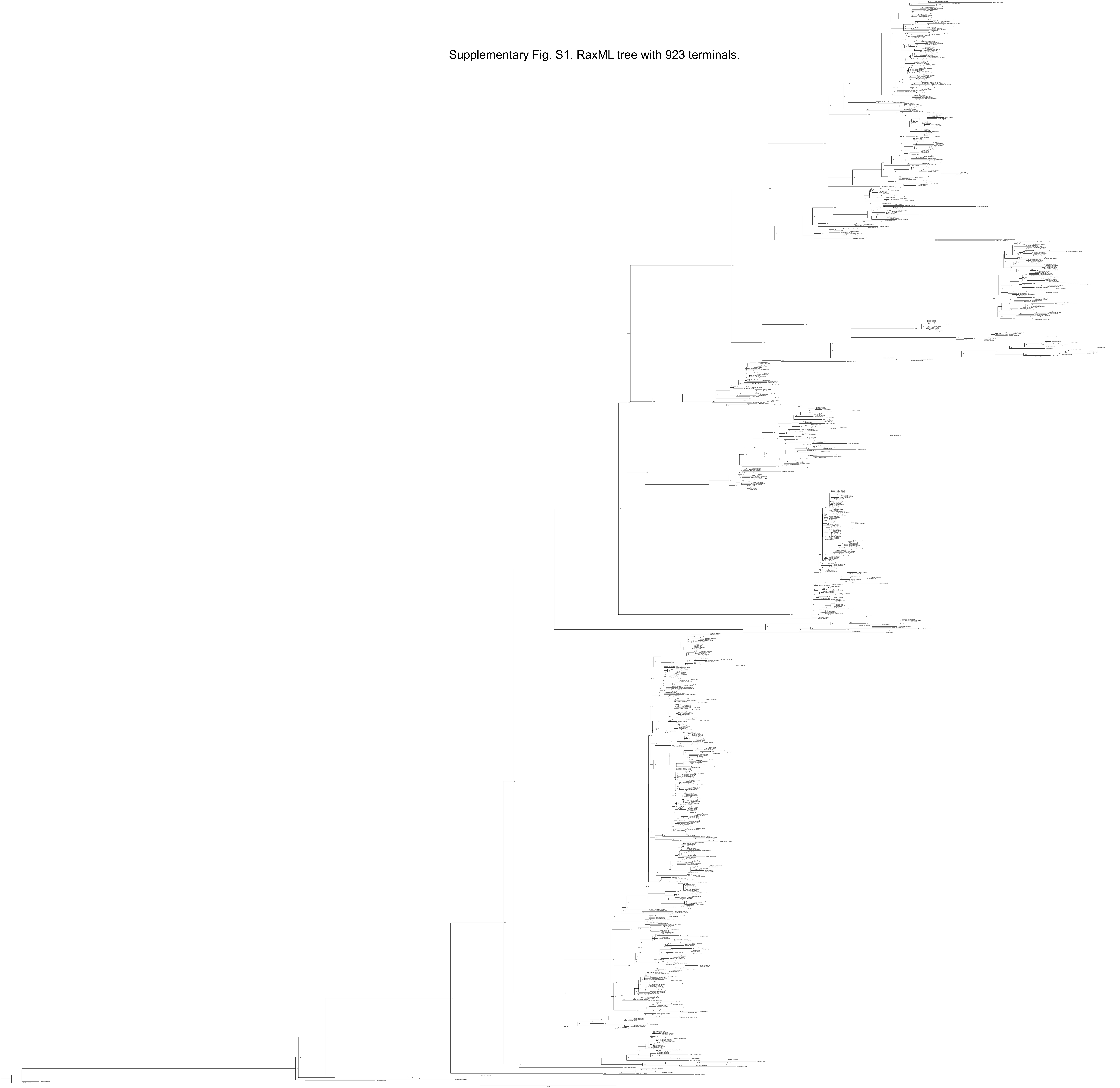

Supplementary Fig. S2. Dated RaxML tree with 923 terminals.

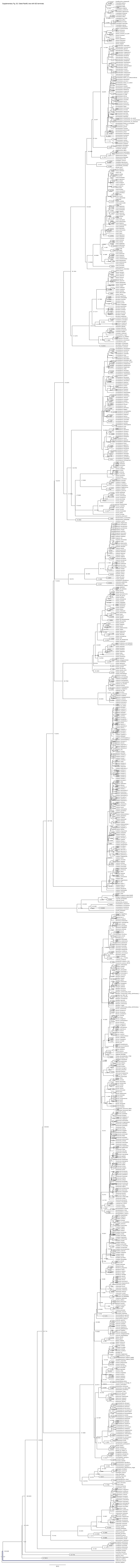

Supplementary Fig. S3. Pruned ultrametric tree with 835 species for diversification analyses, showing family phylogeny and clade ages.

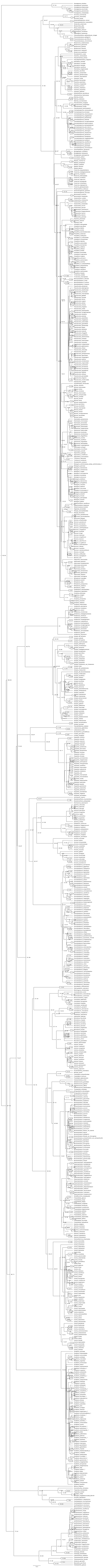

Supplementary Fig. S4. Ancestral character reconstruction for the six traits in Annonaceae using stochastic character mapping.

(A) Habit

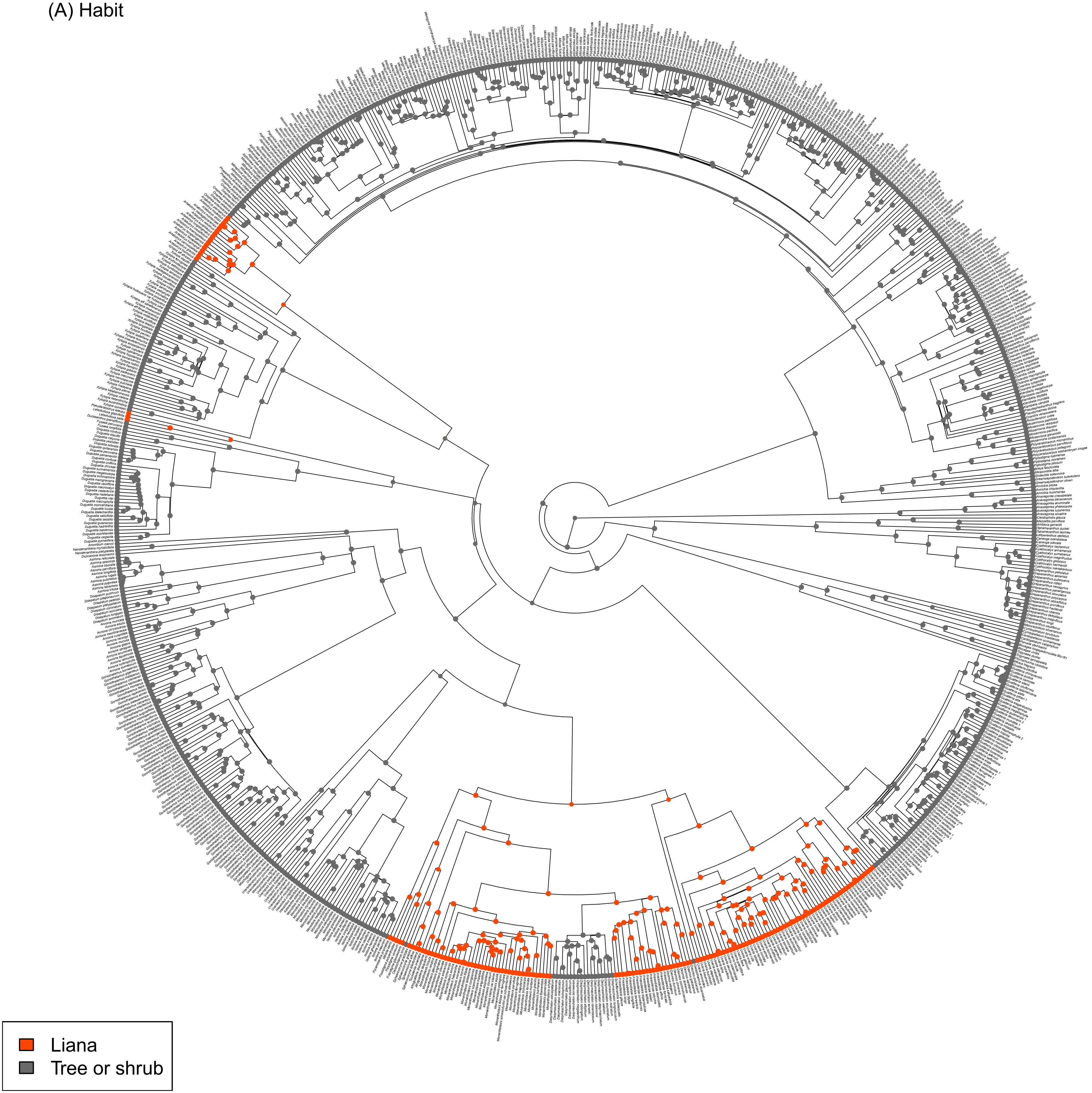

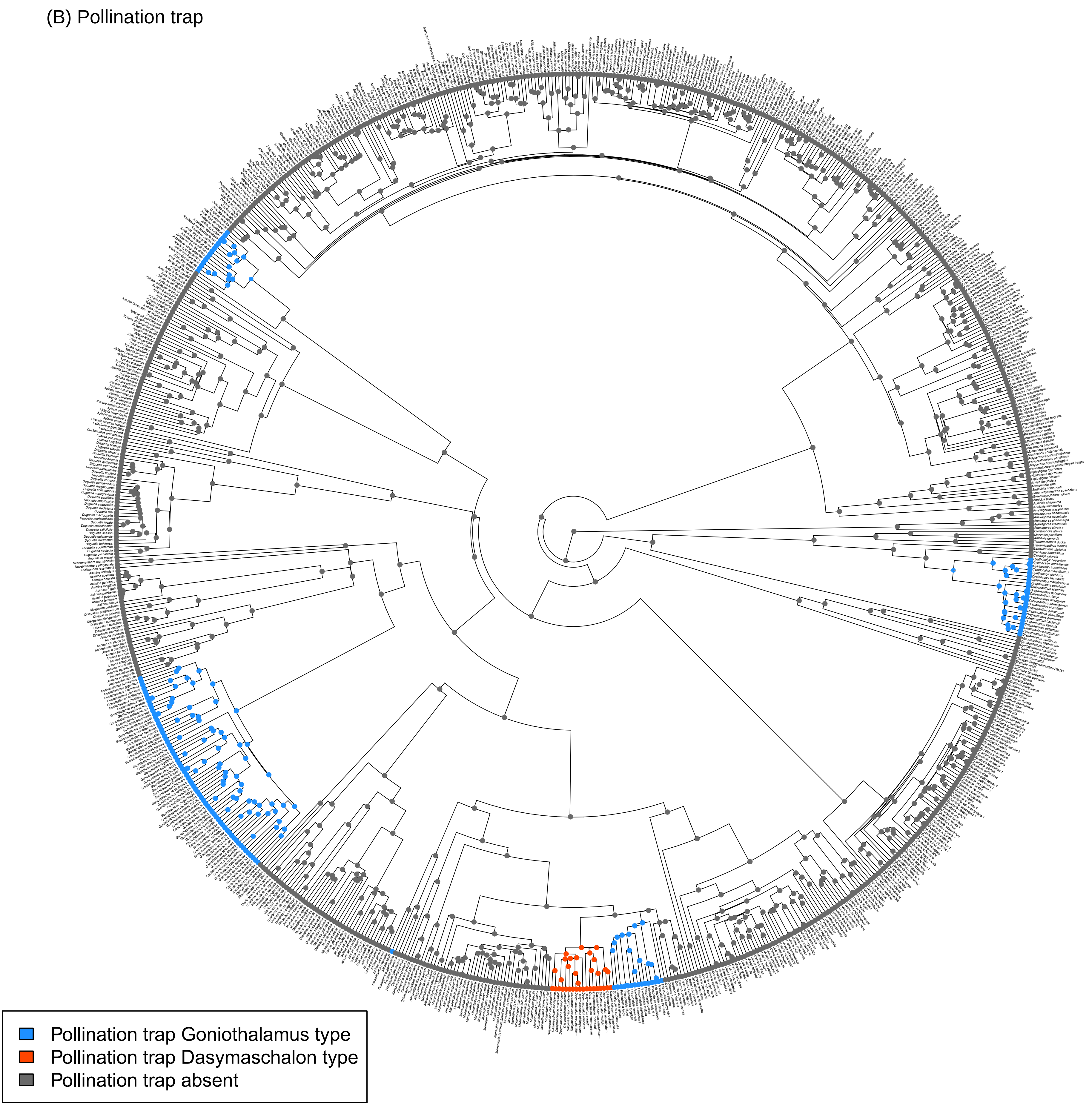
(B) Pollination trap
Pollination trap Goniothalamus type
Pollination trap Dasymaschalon type
Pollination trap absent

(C) Floral sexuality at population level

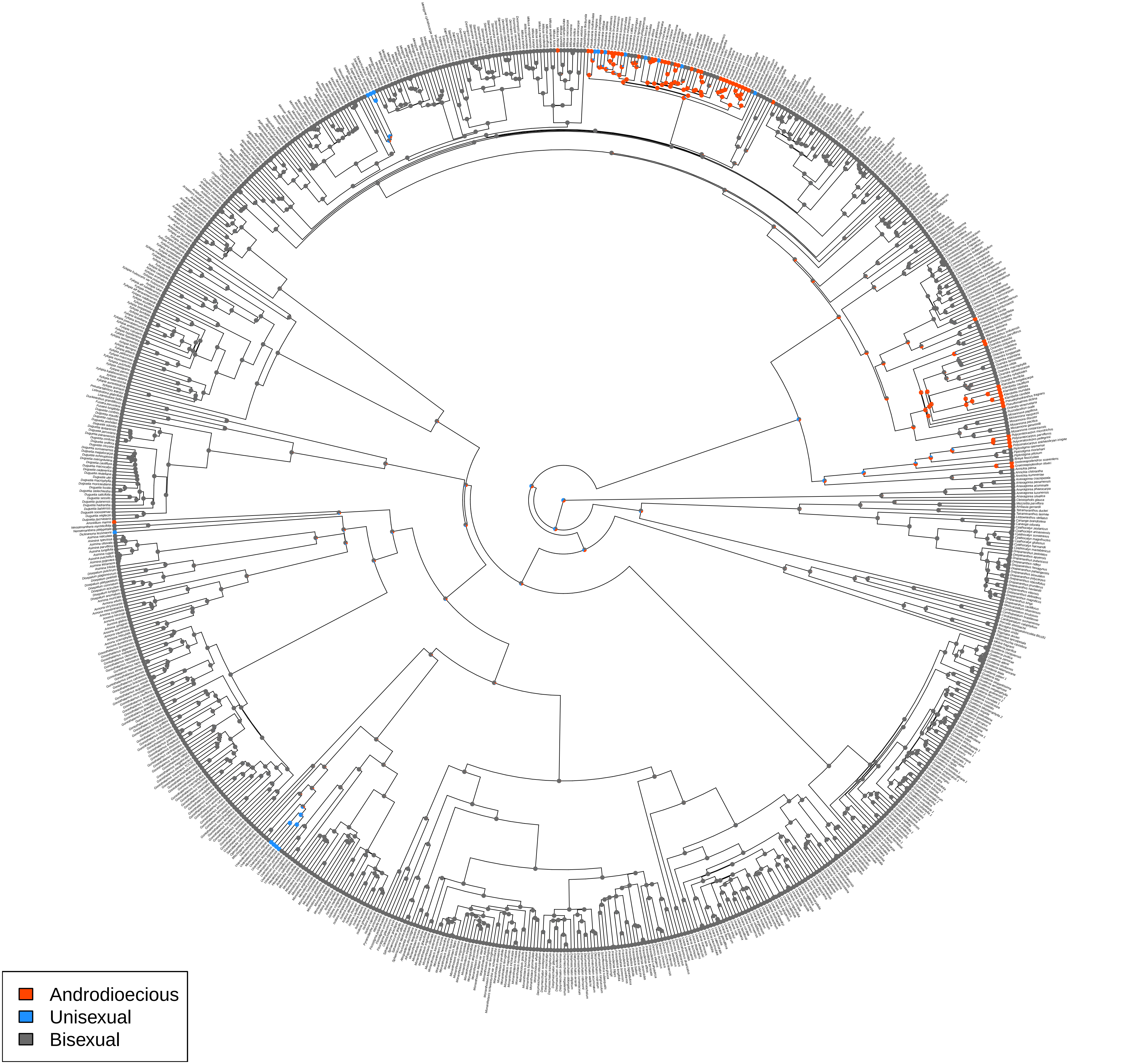

(D) Anther septa

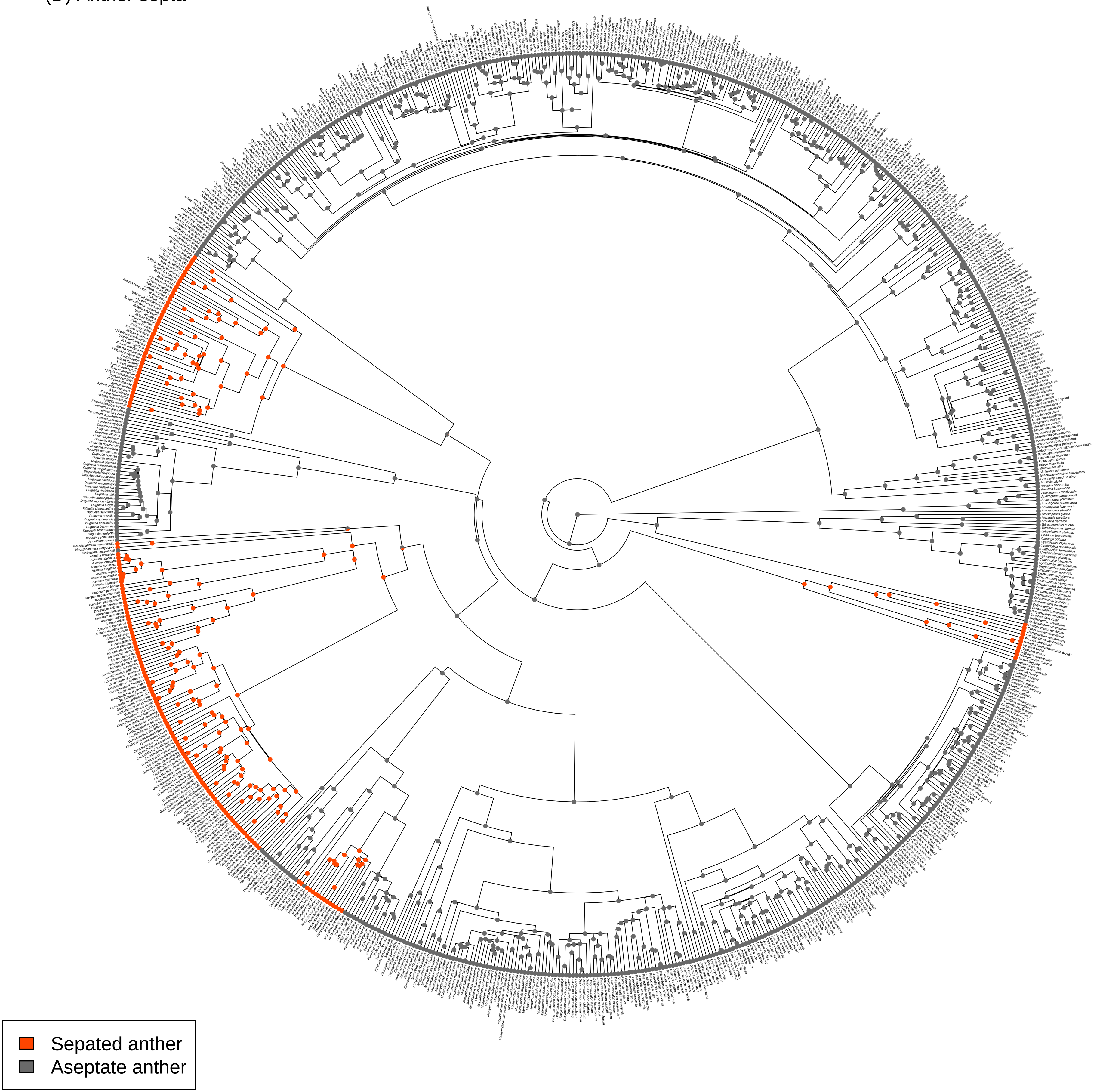

(E) Pollen dispersal unit

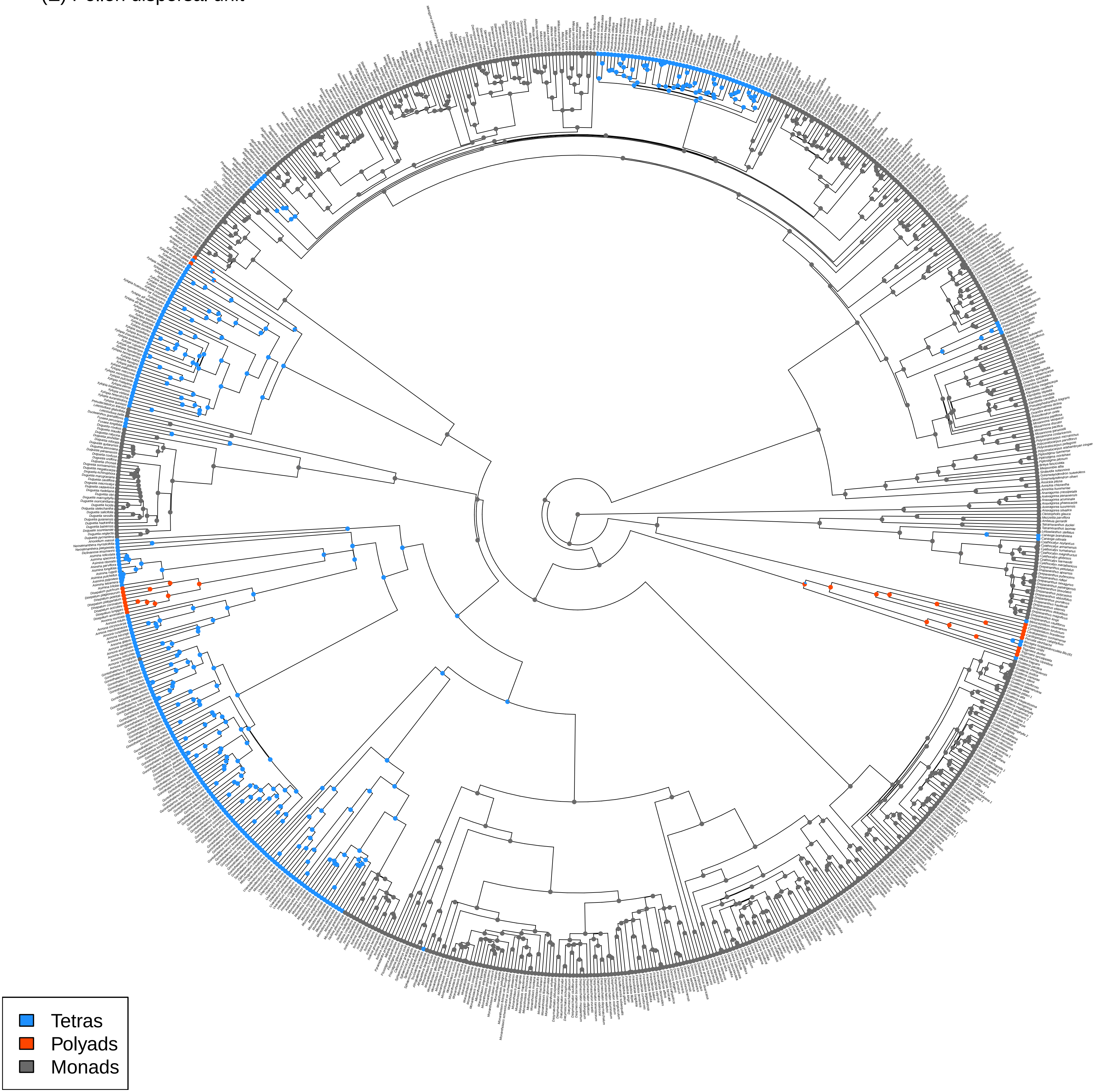

(F) Seed dispersal unit

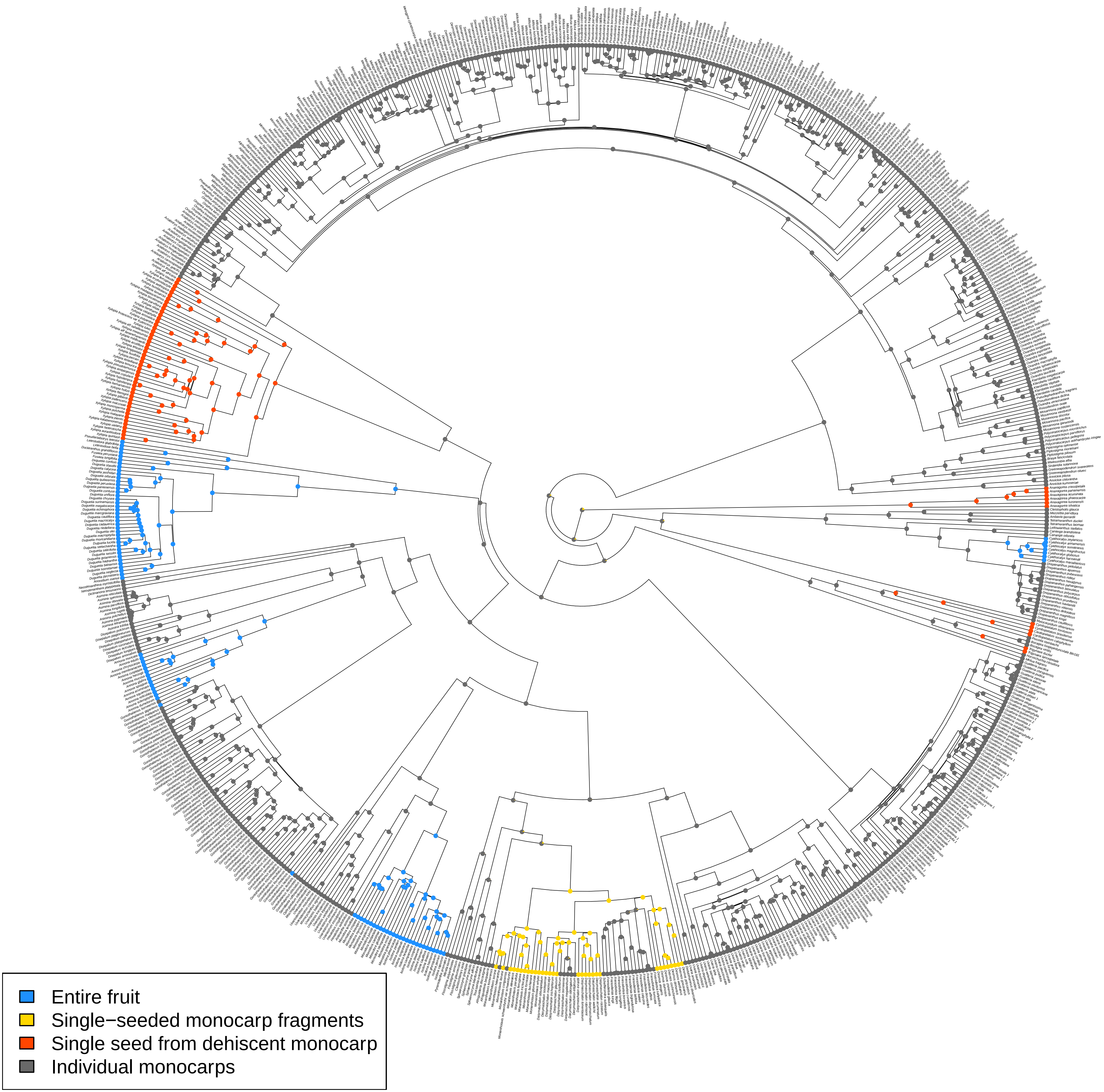

Supplementary Fig. S5. 101-tip genus-level tree used for turboMEDUSA and PGLS analyses.

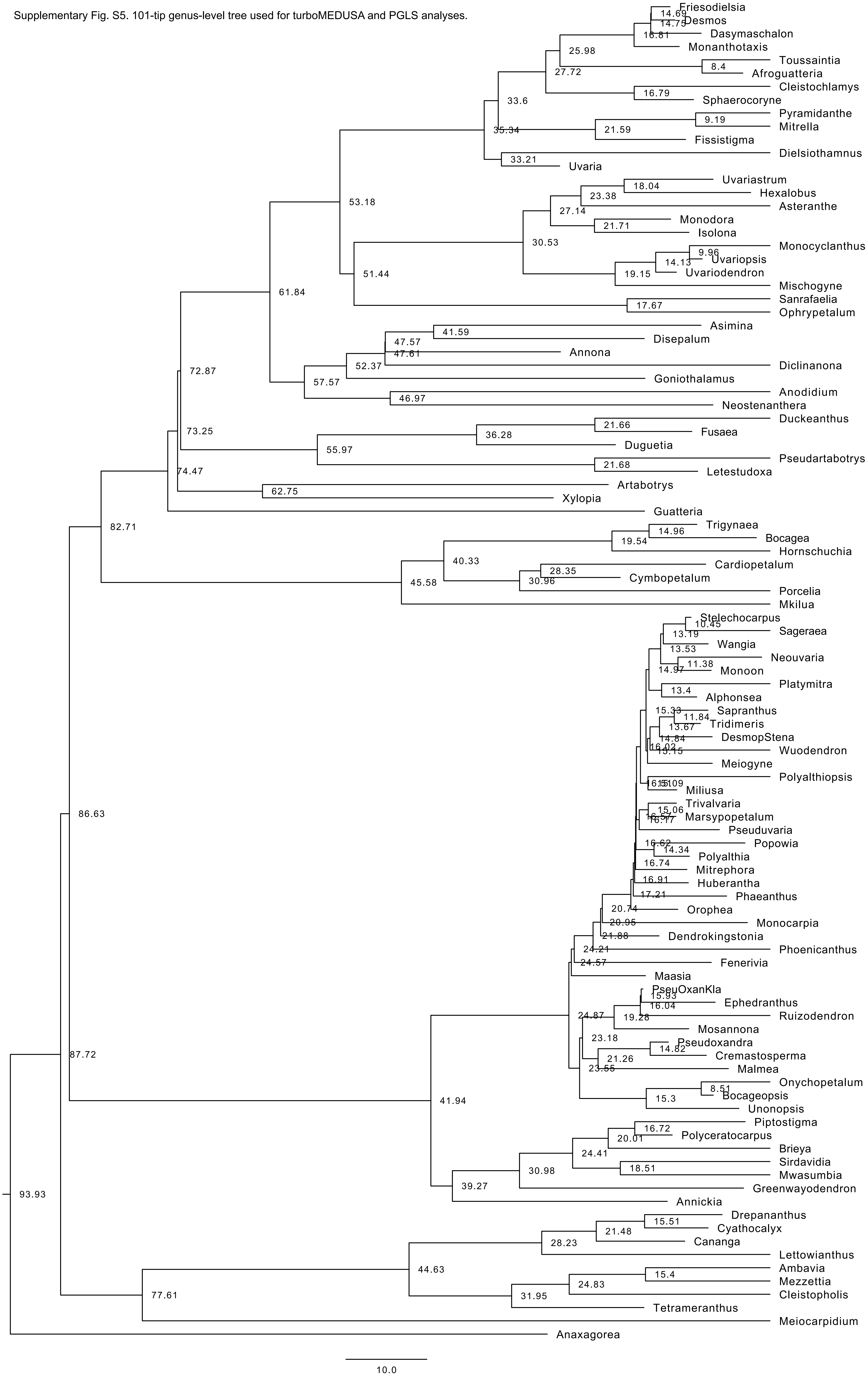

**+ Supplementary Appendix 1.** Species names and GenBank accession numbers of DNA sequences used in this study. -: missing data.

| No. | Voucher information | | GenBank accession numbers | | | | | | | |
|---|---|---|---|---|---|---|---|---|---|---|
| | Tip name in the tree | Taxa name | *atpB-rbcL* | *matK* | *ndhF* | *psbA-trnH* | *rbcL* | *trnL-F* | *trnS-G* | *ycf1* |
| 1 | Afroguatteria_bequaertii | *Afroguatteria bequaertii* (De Wild.) Boutique | - | KX786588 | - | - | KX786627 | KX786629 | - | - |
| 2 | Afroguatteria_discostigma | *Afroguatteria discostigma (Diels) X. Guo & R.M.K. Saunders= Friesodielsia discostigma* (Diels) Steenis | - | KX786595 | - | KX786623 | - | KX786635 | | |
| 3 | Alphonsea_boniana | *Alphonsea boniana* Finet & Gagnep. | - | AY518809 | JQ723785 | - | - | AY319077 | - | - |
| 4 | Alphonsea_elliptica | *Alphonsea elliptica* Hook. f. & Thomson | - | AY518807 | JQ690401 | JQ690402 | - | AY319078 | - | JQ690403 |
| 5 | Alphonsea_glandulosa | *Alphonsea glandulosa* Tan & Xue | - | KY054909 | KY054912 | KY054915 | KY054906 | KY054918 | | KY054921 |
| 6 | Alphonsea_hainanensis | *Alphonsea hainanensis* Merr. & Chun | - | KR530320 | - | KR533235 | KR528675 | - | | - |
| 7 | Alphonsea_javanica | *Alphonsea javanica* Scheff. | - | AY518810 | - | - | - | AY319079 | - | - |
| 8 | Alphonsea_johorensis | *Alphonsea johorensis* J. Sinclair | - | KJ708819 | - | - | - | - | | - |
| 9 | Alphonsea_kinabaluensis | *Alphonsea kinabaluensis* J. Sinclair | - | AY518811 | - | - | AY318968 | AY319080 | | - |
| 10 | Alphonsea_mollis | *Alphonsea mollis* Dunn | - | KY054907 | KY054910 | KY054913 | KY054904 | KY054916 | | KY054919 |
| 11 | Alphonsea_monogyna | *Alphonsea monogyna* Merr. & Chun | - | KY054908 | KY054911 | KY054914 | KY054905 | KY054917 | | KY054920 |
| 12 | Ambavia_gerrardii | *Ambavia gerrardii* (Baill.) Le Thomas | AY578118 | AY220435 | AY218168 | - | JQ513886 | JQ513889 | - | - |
| 13 | Anaxagorea_acuminata | *Anaxagorea acuminata* (Dunal) A. DC. | AY578121 | AY220436 | AY218169 | - | - | - | - | - |
| 14 | Anaxagorea_crassipetala | *Anaxagorea crassipetala* Hemsl. | AY578126 | JQ586481 | - | - | JQ590151 | AY580043 | - | - |
| 15 | Anaxagorea_luzonensis | *Anaxagorea luzonensis* A. Gray | AY578132 | AY518883 | - | - | - | AY319188 | - | - |

| No. | Voucher information | | GenBank accession numbers | | | | | | | |
|---|---|---|---|---|---|---|---|---|---|---|
| | Tip name in the tree | Taxa name | *atpB-rbcL* | *matK* | *ndhF* | *psbA-trnH* | *rbcL* | *trnL-F* | *trnS-G* | *ycf1* |
| 16 | Anaxagorea_panamensis | *Anaxagorea panamensis* Standl. | AY578135 | GQ981933 | - | GQ982144 | GQ981662 | AY580052 | - | - |
| 17 | Anaxagorea_phaeocarpa | *Anaxagorea phaeocarpa* Mart. | EF179244 | AY238960 | EF179279 | AY841426 | AY238952 | EF179316 | EF179321 | - |
| 18 | Anaxagorea_silvatica | *Anaxagorea silvatica* R. E. Fr. | EF179245 | AY743477 | EF179280 | AY841427 | AY743439 | AY743458 | EF179322 | - |
| 19 | Annickia_chlorantha | *Annickia affinis* (Exell) Versteegh & Sosef | AY841370 | AY841393 | AY841401 | KC667934 | AY841594 | AY841671 | AY841550 | - |
| 20 | Annickia_kummeriae | *Annickia kummeriae* (Engl. & Diels) Setten & Maas | - | AY238961 | - | AY841443 | AY238959 | AY319171 | - | - |
| 21 | Annickia_pilosa | *Annickia pilosa* (Exell) Setten & Maas | AY841371 | AY743488 | AY841402 | AY841444 | AY743450 | AY743469 | AY841551 | - |
| 22 | Annona_acuminata | *Annona acuminata* Saff. | - | GQ981934 | - | GQ982146 | GQ981664 | - | - | - |
| 23 | Annona_chrysocarpa | *Annona chrysocarpa* Lepr. ex A. Rich. | - | - | - | - | EU420868 | EU420850 | - | - |
| 24 | Annona_cuspidata | *Annona cuspidata* (Mart.) H. Rainer | - | - | - | - | EU420869 | EU420851 | - | - |
| 25 | Annona_dumetorum | *Annona dumetorum* R. E. Fr. | - | GQ139704 | - | GQ139844 | EU420856 | EU420838 | - | GU937352 |
| 26 | Annona_edulis | *Annona edulis* (Triana & Planch.) H. Rainer | - | - | - | - | AY841655 | AY841733 | - | - |
| 27 | Annona_glabra | *Annona glabra* L. | EF179246 | GQ139717 | EF179281 | DQ125116 | AY841596 | AY841673 | EF179323 | GU937365 |
| 28 | Annona_herzogii | *Annona herzogii* (R. E. Fr.) H. Rainer | EF179273 | DQ125062 | EF179308 | DQ125132 | AY841656 | AY841734 | EF179350 | - |
| 29 | Annona_membranacea | *Annona membranacea* R. E. Fr. | - | JQ586514 | - | - | JQ590184 | - | - | - |
| 30 | Annona_mucosa | *Annona mucosa* Jacq. | - | GQ139705 | - | GQ139845 | EU420870 | EU420852 | - | GU937353 |
| 31 | Annona_muricata | *Annona muricata* L. | EF179247 | AF543722 | EF179282 | AY841428 | AY743440 | AY743459 | EF179324 | - |
| 32 | Annona_reticulata | *Annona reticulata* L. | - | JQ586491 | - | JX856821 | EU420863 | EU420845 | - | - |
| 33 | Annona_sclerophylla | *Annona sclerophylla* Saff. | - | GQ139718 | - | GQ139858 | - | GQ139892 | - | GU937366 |

| No. | Voucher information | | GenBank accession numbers | | | | | | | |
|---|---|---|---|---|---|---|---|---|---|---|
| | Tip name in the tree | Taxa name | *atpB-rbcL* | *matK* | *ndhF* | *psbA-trnH* | *rbcL* | *trnL-F* | *trnS-G* | *ycf1* |
| 34 | Annona_spraguei | *Annona spraguei* Saff. | - | GQ981935 | - | GQ982147 | GQ981665 | - | - | - |
| 35 | Annona_squamosa | *Annona squamosa* L. | - | EU715064 | - | EU715086 | EU420865 | EU420847 | - | - |
| 36 | Anodidium_mannii | *Anonidium mannii* (Oliv.) Engl. & Diels | EF179248 | DQ125051 | EF179283 | DQ125117 | AY841598 | AY841675 | EF179325 | - |
| 37 | Artabotrys_aurantiacus | *Artabotrys aurantiacus* | - | KM924834 | KM924864 | KM924965 | KM924904 | KM924932 | - | - |
| 38 | Artabotrys_brachypetalus | *Artabotrys brachypetalus* Benth. | - | - | - | - | JF265293 | - | - | - |
| 39 | Artabotrys_carnosipetala | *Artabotrys carnosipetalus* | - | KM924835 | KM924865 | KM924966 | KM924905 | KM924933 | - | - |
| 40 | Artabotrys_crassifolium | *Artabotrys crassifolius* | - | KM924836 | KM924866 | KM924967 | KM924906 | KM924934 | - | - |
| 41 | Artabotrys_gracilis | *Artabotrys gracilis* | - | KM924837 | KM924867 | KM924968 | KM924907 | KM924935 | - | - |
| 42 | Artabotrys_harmandii | *Artabotrys harmandii* | - | KM924838 | KM924868 | KM924969 | KM924908 | KM924936 | - | - |
| 43 | Artabotrys_hexapetalus | *Artabotrys hexapetalus* (L. f.) Bhandari | EF179249 | AY238962 | EF179284 | AY841429 | AY238953 | EF179317 | EF179326 | - |
| 44 | Artabotrys_hongkongensis | *Artabotrys honkongensis* | - | KM924839 | KM924869 | KM924970 | KM924909 | KM924937 | - | - |
| 45 | Artabotrys_longistigmatus | *Artabotrys longistigmatus* | - | KM924840 | KM924870 | KM924971 | KM924910 | KM924938 | - | - |
| 46 | Artabotrys_madagascariensis | *Artabotrys madagascariensis* | - | - | KM924871 | KM924972 | - | KM924939 | - | - |
| 47 | Artabotrys_modestus | *Artabotrys modestus* | - | KM924841 | KM924872 | KM924973 | KM924911 | KM924940 | - | - |
| 48 | Artabotrys_monteiroae | *Artabotrys monteiroae* | - | KM924842 | KM924873 | KM924974 | KM924912 | KM924941 | - | - |
| 49 | Artabotrys_pierreanus | *Artabotrys pierreanus* | - | KM924843 | KM924874 | KM924975 | KM924913 | KM924942 | - | - |
| 50 | Artabotrys_sp_Gabon | *Artabotrys spec.* Gabon | EF179250 | DQ125052 | EF179285 | DQ125118 | AY841599 | AY841676 | EF179327 | - |
| 51 | Artabotrys_sp_PNG | *Artabotrys spec.* PNG | - | KM924844 | KM924875 | KM924976 | KM924914 | KM924943 | - | - |

| No. | Voucher information | | GenBank accession numbers | | | | | | | |
|---|---|---|---|---|---|---|---|---|---|---|
| | Tip name in the tree | Taxa name | *atpB-rbcL* | *matK* | *ndhF* | *psbA-trnH* | *rbcL* | *trnL-F* | *trnS-G* | *ycf1* |
| 52 | Artabotrys_sp_Sumatra | *Artabotrys spec.* Sumatra | - | KM924845 | KM924876 | KM924977 | KM924915 | KM924944 | - | - |
| 53 | Artabotrys_stolzii | *Artabotrys stolzii* | - | KM924846 | KM924877 | KM924978 | KM924916 | KM924945 | - | - |
| 54 | Artabotrys_thomsonii | *Artabotrys thomsonii* | - | KM924847 | KM924878 | KM924979 | KM924917 | KM924946 | - | - |
| 55 | Artabotrys_velutinus | *Artabotrys velutinus* | - | KM924848 | KM924879 | KM924980 | KM924918 | KM924947 | - | - |
| 56 | Asimina_incana | *Asimina incana* (W. Bartram) Exell | - | GQ139730 | - | GQ139873 | - | GQ139903 | - | GU937377 |
| 57 | Asimina_longifolia | *Asimina longifolia* Kral | EF179251 | GQ139707 | EF179286 | DQ125119 | DQ124939 | GQ139885 | EF179328 | GU937355 |
| 58 | Asimina_obovata | *Asimina obovata* (Willd.) Nash | - | GQ139716 | - | GQ139855 | - | GQ139908 | - | GU937363 |
| 59 | Asimina_parviflora | *Asimina parviflora* (Michx.) Dunal | - | GQ139703 | - | GQ139843 | - | GQ139878 | - | GU937351 |
| 60 | Asimina_pulchellus | *Asimina pulchellus* (Small) Rehder & Dayton | - | GQ139714 | - | GQ139854 | - | GQ139889 | - | GU937362 |
| 61 | Asimina_pygmaea | *Asimina pygmaea* (W. Bartram) Dunal | - | GQ139726 | - | GQ139868 | - | GQ139899 | - | GU937356 |
| 62 | Asimina_reticulata | *Asimina reticulata* Shuttlew. ex Chapm. | - | GQ139713 | - | GQ139863 | - | GQ139906 | - | GU937361 |
| 63 | Asimina_rugelii | *Asimina rugelii* B. L. Rob. | - | GQ139706 | - | GQ139846 | JQ513887 | GQ139881 | - | GU937354 |
| 64 | Asimina_tetramera | *Asimina tetramera* Small | - | GQ139702 | - | GQ139870 | - | GQ139877 | - | GU937350 |
| 65 | Asimina_triloba | *Asimina triloba* (L.) Dunal | EF179252 | GQ139711 | AY218171 | AY841430 | AY743441 | AY743460 | EF179329 | GU937349 |
| 66 | Asteranthe_asterias | *Asteranthe asterias* (S. Moore) Engl. & Diels | - | - | EU169711 | EU169734 | EU169757 | EU169779 | EU169801 | - |
| 67 | Bocagea_longepedunculata_Blo181 | *Bocagea longepedunculata* Mart. Blo181 | MF322668 | MF322630 | MF322640 | MF322655 | MF322645 | MF322659 | - | - |
| 68 | Bocagea_longepedunculata_Blo460 | *Bocagea longepedunculata* Mart. Blo460 | MF322669 | MF322631 | MF322641 | - | MF322646 | - | - | - |

| No. | Voucher information | | GenBank accession numbers | | | | | | | |
|---|---|---|---|---|---|---|---|---|---|---|
| | Tip name in the tree | Taxa name | *atpB-rbcL* | *matK* | *ndhF* | *psbA-trnH* | *rbcL* | *trnL-F* | *trnS-G* | *ycf1* |
| 69 | Bocagea_viridis | *Bocagea viridis* A. St.-Hil. | MF322670 | MF322632 | MF322642 | - | MF322647 | MF322660 | - | - |
| 70 | Bocageopsis_canescens | *Bocageopsis canescens* (Spruce ex Benth.) R. E. Fr. | - | JQ690409 | JQ690410 | JQ690411 | JQ690407 | JQ690408 | - | JQ690412 |
| 71 | Bocageopsis_multiflora | *Bocageopsis multiflora* (Mart.) R. E. Fr. | - | DQ018262 | - | AY841445 | AY841600 | AY841678 | - | - |
| 72 | Bocageopsis_pleiosperma | *Bocageopsis pleiosperma* Maas | - | - | - | AY841446 | AY841601 | AY841679 | - | - |
| 73 | Brieya_fasciculata | *Brieya fasciculata* De Wild. | - | - | - | AY841497 | AY841647 | AY841725 | - | - |
| 74 | Cananga_brandisiana | *Cananga brandisiana* (Pierre) I. M. Turner | - | HM173744 | - | HM173715 | HM173801 | HM173772 | - | - |
| 75 | Cananga_odorata | *Cananga odorata* (Lam.) Hook. f. & Thomson | AY841372 | AY220438 | AY841403 | AY841431 | AY841602 | AY841680 | AY841548 | - |
| 76 | Cardiopetalum_calophyllum | *Cardiopetalum calophyllum* Schltdl. | MF322672 | MF322634 | MF322644 | - | MF322649 | MF322662 | - | - |
| 77 | Cardiopetalum_surinamense | *Cardiopetalum surinamense* R. E. Fr. | MF322673 | MF322635 | - | - | MF322650 | MF322663 | - | - |
| 78 | Cleistochlamys_kirkii | *Cleistochlamys kirkii* (Benth.) Oliv. | - | - | KM924880 | KM924981 | - | KM924948 | - | - |
| 79 | Cleistopholis_glauca | *Cleistopholis glauca* Pierre ex Engl. & Diels | AY841373 | AY841395 | AY841404 | AY841432 | AY841603 | AY841681 | AY841549 | - |
| 80 | Coelocaryon_preussii | *Coelocaryon preussii* Warb. | - | AY743475 | JQ437546 | - | AY743437 | AY743456 | - | - |
| 81 | Cremastosperma_brevipes | *Cremastosperma brevipes* (DC.) R. E. Fr. | AY841374 | AY743550 | AY841405 | AY841447 | AY743527 | AY743573 | AY841552 | - |
| 82 | Cremastosperma_bullatum | *Cremastosperma bullatum* Pirie | - | AY743560 | DQ018140 | AY841459 | DQ018235 | DQ018188 | - | - |

| No. | Voucher information | | GenBank accession numbers | | | | | | | |
|---|---|---|---|---|---|---|---|---|---|---|
| | Tip name in the tree | Taxa name | *atpB-rbcL* | *matK* | *ndhF* | *psbA-trnH* | *rbcL* | *trnL-F* | *trnS-G* | *ycf1* |
| 83 | Cremastosperma_cauliflorum | *Cremastosperma cauliflorum* R. E. Fr. | AY841375 | AY743542 | AY841406 | DQ018240 | AY743519 | AY743565 | AY841553 | - |
| 84 | Cremastosperma_cenepense | *Cremastosperma cenepense* Pirie & Zapata | - | DQ018277 | - | DQ018257 | DQ018236 | - | - | - |
| 85 | Cremastosperma_gracilipes | *Cremastosperma gracilipes* R. E. Fr. | - | AY743544 | - | - | AY743521 | AY743567 | - | - |
| 86 | Cremastosperma_leiophyllum | *Cremastosperma leiophyllum* (Diels) R. E. Fr. | - | AY743546 | DQ018123 | AY841449 | AY743523 | AY743569 | - | - |
| 87 | Cremastosperma_macrocarpum | *Cremastosperma macrocarpum* Maas | - | AY743551 | DQ018129 | AY841450 | AY743528 | AY743574 | - | - |
| 88 | Cremastosperma_magdalenae | *Cremastosperma magdalenae* Pirie | - | DQ018279 | DQ018143 | AY841460 | AY841521 | AY841535 | - | - |
| 89 | Cremastosperma_megalophyllum | *Cremastosperma megalophyllum* R. E. Fr. | - | AY743545 | DQ018121 | AY841451 | AY743517 | AY743563 | - | - |
| 90 | Cremastosperma_microcarpum | *Cremastosperma microcarpum* R. E. Fr. | - | AY743539 | DQ018120 | AY841452 | AY743516 | AY319172 | - | - |
| 91 | Cremastosperma_monospermum | *Cremastosperma monospermum* (Rusby) R. E. Fr. | - | AY743547 | DQ018124 | AY841453 | AY743524 | AY743570 | - | - |
| 92 | Cremastosperma_napoense | *Cremastosperma napoense* Pirie | - | DQ018265 | DQ018127 | DQ018242 | DQ018224 | DQ018177 | - | - |
| 93 | Cremastosperma_novogranatense | *Cremastosperma novogranatense* R. E. Fr. | - | AY743552 | - | - | AY743529 | AY743575 | - | - |
| 94 | Cremastosperma_oblongum | *Cremastosperma oblongum* R. E. Fr. | - | DQ018266 | DQ018128 | DQ018241 | DQ018225 | DQ018178 | - | - |

| No. | Voucher information | | GenBank accession numbers | | | | | | | |
|---|---|---|---|---|---|---|---|---|---|---|
| | Tip name in the tree | Taxa name | *atpB-rbcL* | *matK* | *ndhF* | *psbA-trnH* | *rbcL* | *trnL-F* | *trnS-G* | *ycf1* |
| 95 | Cremastosperma_panamense | *Cremastosperma panamense* Maas | - | AY743553 | DQ018131 | DQ018246 | AY743530 | AY743576 | - | - |
| 96 | Cremastosperma_pedunculatum | *Cremastosperma pedunculatum* (Diels) R. E. Fr. | - | DQ018269 | DQ018135 | DQ018248 | DQ018228 | DQ018181 | - | - |
| 97 | Cremastosperma_pendulum | *Cremastosperma pendulum* (Ruiz & Pav.) R. E. Fr. | - | AY743554 | DQ018132 | DQ018247 | DQ018233 | DQ018186 | - | - |
| 98 | Cremastosperma_peruvianum | *Cremastosperma peruvianum* R. E. Fr. | - | AY743557 | - | - | AY743534 | AY743580 | - | - |
| 99 | Cremastosperma_venezuelanum | *Cremastosperma venezuelanum* Pirie | - | AY743559 | DQ018134 | AY841457 | AY743536 | AY743582 | - | - |
| 100 | Cremastosperma_westrae | *Cremastosperma westrae* Pirie | - | DQ018272 | DQ018136 | DQ018252 | DQ018231 | DQ018184 | - | - |
| 101 | Cremastosperma_yamayakatense | *Cremastosperma yamayakatense* Pirie | - | DQ018267 | DQ018139 | DQ018244 | DQ018226 | DQ018179 | - | - |
| 102 | Cyathocalyx_annamensis | *Cyathocalyx annamensis* Ast | - | HM173748 | - | HM173719 | HM173805 | HM173776 | - | - |
| 103 | Cyathocalyx_globosus | *Cyathocalyx globosus* Merr. | - | HM173725 | - | HM173696 | HM173782 | HM173754 | - | - |
| 104 | Cyathocalyx_harmandii | *Cyathocalyx harmandii* (Finet & Gagnep.) R. J. Wang & R. M. K. Saunders | - | HM173726 | - | HM173697 | HM173783 | HM173755 | - | - |
| 105 | Cyathocalyx_magnifructus | *Cyathocalyx magnifructus* R. J. Wang & R. M. K. Saunders | - | HM173730 | - | HM173701 | HM173787 | HM173759 | - | - |
| 106 | Cyathocalyx_martabanicus | *Cyathocalyx martabanicus* Hook. f. & Thomson | EF179253 | DQ125054 | EF179288 | DQ125120 | AY841605 | AY841683 | EF179330 | - |
| 107 | Cyathocalyx_sumatranus | *Cyathocalyx sumatranus* Scheff. | - | HM173737 | - | HM173708 | HM173794 | HM173766 | - | - |

| No. | Voucher information | | GenBank accession numbers | | | | | | | |
|---|---|---|---|---|---|---|---|---|---|---|
| | Tip name in the tree | Taxa name | *atpB-rbcL* | *matK* | *ndhF* | *psbA-trnH* | *rbcL* | *trnL-F* | *trnS-G* | *ycf1* |
| 108 | Cyathocalyx_zeylanicus | *Cyathocalyx zeylanicus* Champ. ex Hook. f. & Thomson | - | HM173739 | - | HM173710 | HM173796 | HM173768 | - | - |
| 109 | Cymbopetalum_brasiliense | *Cymbopetalum brasiliense* (Vell.) Benth. ex Baill. | EF179254 | DQ125055 | EF179289 | DQ125121 | AY841608 | AY841686 | EF179331 | - |
| 110 | Cymbopetalum_torulosum | *Cymbopetalum torulosum* G. E. Schatz | - | GQ139720 | - | GQ139859 | AY743442 | GQ139894 | - | - |
| 111 | Dasymaschalon_acuminatum | *Dasymaschalon acuminatum* Jing Wang & R. M. K. Saunders | - | JQ768546 | JQ768587 | JQ768625 | JQ768666 | JQ768706 | - | - |
| 112 | Dasymaschalon_borneense | *Dasymaschalon borneense* Nurmawati | - | JQ768547 | - | JQ768626 | JQ768667 | JQ768707 | - | - |
| 113 | Dasymaschalon_clusiflorum | *Dasymaschalon clusiflorum* (Merr.) Merr. | - | JQ768548 | - | JQ768627 | JQ768668 | JQ768708 | - | - |
| 114 | Dasymaschalon_dasymaschalum | *Dasymaschalon dasymaschalum* (Blume) I. M. Turner | - | JQ768549 | JQ768588 | JQ768628 | JQ768669 | JQ768709 | - | - |
| 115 | Dasymaschalon_ellipticum | *Dasymaschalon ellipticum* Nurmawati | - | JQ768550 | JQ768589 | JQ768629 | JQ768670 | JQ768710 | - | - |
| 116 | Dasymaschalon_evrardii | *Dasymaschalon evrardii* Ast | - | JQ768551 | JQ768590 | JQ768630 | JQ768671 | JQ768711 | - | - |
| 117 | Dasymaschalon_glaucum | *Dasymaschalon glaucum* Merr. & Chun | - | JQ768553 | JQ768592 | JQ768632 | JQ768673 | JQ768713 | - | - |
| 118 | Dasymaschalon_lomentaceum | *Dasymaschalon lomentaceum* Finet & Gagnep. | - | EU715065 | JQ768593 | EU715087 | JQ768674 | JQ768714 | - | - |
| 119 | Dasymaschalon_longiusculum | *Dasymaschalon longiusculum* (Bân) Jing Wang & R. M. K. Saunders | - | JQ768556 | - | JQ768635 | JQ768676 | JQ768716 | - | - |

| No. | Voucher information | | GenBank accession numbers | | | | | | | |
|---|---|---|---|---|---|---|---|---|---|---|
| | Tip name in the tree | Taxa name | *atpB-rbcL* | *matK* | *ndhF* | *psbA-trnH* | *rbcL* | *trnL-F* | *trnS-G* | *ycf1* |
| 120 | Dasymaschalon_macrocalyx | *Dasymaschalon macrocalyx* Finet & Gagnep. | EF179255 | EF179277 | EF179290 | EF179313 | AY841610 | AY841688 | EF179332 | - |
| 121 | Dasymaschalon_megalanthum | *Dasymaschalon megalanthum* (Merr.) Jing Wang & R.M.K. Saunders | - | JQ768558 | JQ768596 | JQ768637 | JQ768678 | JQ768718 | - | - |
| 122 | Dasymaschalon_oblongatum | *Dasymaschalon oblongatum* Merr. | - | JQ768559 | JQ768597 | JQ768638 | JQ768679 | JQ768719 | - | - |
| 123 | Dasymaschalon_robinsonii | *Dasymaschalon robinsonii* Ast | - | JQ768561 | - | JQ768640 | JQ768681 | JQ768721 | - | - |
| 124 | Dasymaschalon_rostratum | *Dasymaschalon rostratum* Merr. & Chun | - | JQ768562 | JQ768599 | JQ768641 | JQ768682 | JQ768722 | - | - |
| 125 | Dasymaschalon_sootepense | *Dasymaschalon sootepense* Craib | - | AY743481 | JQ768600 | JQ768642 | AY743443 | AY743462 | - | - |
| 126 | Dasymaschalon_trichophorum | *Dasymaschalon trichophorum* Merr. | - | JQ768565 | JQ768602 | JQ768644 | JQ768685 | JQ768725 | - | - |
| 127 | Dasymaschalon_wallichii | *Dasymaschalon wallichii* (Hook. f. & Thomson) Jing Wang & R. M. K. Saunders | - | JQ768566 | - | JQ768645 | JQ768686 | JQ768726 | - | - |
| 128 | Dasymaschalon_obtusipetalum | *Dasymaschalon yunnanense* (Hu) Bân | - | JQ768560 | JQ768598 | JQ768639 | JQ768680 | JQ768720 | - | - |
| 129 | Degeneria_vitiensis | *Denegeria roseiflora* J.M. Miller *(in Genebank as Degeneria vitiensis I.W.Bailey & A.C.Sm.)* | AB332086 (include in 15-atpB-rbcL | AB055549 | AY394736 | AB332084 | L12643 | | | |

| No. | Voucher information | | GenBank accession numbers | | | | | | | |
|---|---|---|---|---|---|---|---|---|---|---|
| | Tip name in the tree | Taxa name | *atpB-rbcL* | *matK* | *ndhF* | *psbA-trnH* | *rbcL* | *trnL-F* | *trnS-G* | *ycf1* |
| 130 | Dendrokingstonia_gardneri | *Dendrokingstonia gardneri* Chaowasku | - | KJ418391 | KJ418385 | KJ418399 | KJ418381 | KJ418406 | - | KJ418378 |
| 131 | Dendrokingstonia_nervosa | *Dendrokingstonia nervosa* (Hook. f. & Thomson) Rauschert | - | KJ418392 | KJ418386 | KJ418400 | KJ418382 | KJ418407 | - | - |
| 132 | Desmopsis_bibracteata | *Desmopsis bibracteata* (B. L. Rob.) Saff. | - | JQ586480 | - | - | JQ590148 | - | - | - |
| 133 | Desmopsis_lanceolata | Desmopsis lanceolata Lundell | | KU727296 | - | KU727337 | KU727378 | KU727414 | - | - |
| 134 | Desmopsis_microcarpa | *Desmopsis microcarpa* R. E. Fr. | - | AY518804 | JX544771 | AY841461 | AY319059 | AY319173 | - | JX544758 |
| 135 | Desmopsis_panamensis | *Desmopsis panamensis* (B. L. Rob.) Saff. | - | GQ981981 | - | GQ982207 | GQ981723 | - | - | - |
| 136 | Desmopsis_schippii | *Desmopsis schippii* Standl. | - | AY518805 | JQ723786 | - | JQ590166 | AY319174 | - | - |
| 137 | Desmopsis_sp_A | *Desmopsis* sp. A | - | KU727293 | - | KU727334 | KU727375 | KU727411 | - | - |
| 138 | Desmopsis_sp_B | *Desmopsis* sp. B | - | KU727297 | - | KU727338 | KU727379 | KU727415 | - | - |
| 139 | Desmopsis_sp_C | *Desmopsis* sp. C | - | KU727323 | - | KU727365 | KU727403 | KU727442 | - | - |
| 140 | Desmopsis_trunciflora | *Desmopsis trunciflora* (Schltdl. & Cham.) G.E. Schatz | - | - | - | KU727340 | - | KU727417 | - | - |
| 141 | Desmopsis_uxpanapensis | *Desmopsis uxpanapensis* | - | KU727300 | - | KU727342 | KU727382 | KU727419 | - | - |
| 142 | Desmopsis_wendtii | *Desmopsis wendtii* G.E. Schatz ined. | - | KU727302 | - | KU727344 | KU727384 | KU727421 | - | - |
| 143 | Desmos_chinensis | *Desmos chinensis* Lour. | - | JQ768567 | JQ768603 | JQ768646 | JQ762414 | JQ762415 | - | - |
| 144 | Desmos_chinensis_var_lawii | *Desmos chinensis* var. *lawii* (Hook.f. & Thomson) Bân | - | KC933937 | KC933943 | KC933941 | KC933935 | KC933939 | | |
| 145 | Desmos_cochinchinensis | *Desmos cochinchinensis* Lour. | - | JQ768568 | JQ768604 | JQ768647 | JQ768688 | JQ768728 | - | - |

| No. | Voucher information | | GenBank accession numbers | | | | | | | |
|---|---|---|---|---|---|---|---|---|---|---|
| | Tip name in the tree | Taxa name | *atpB-rbcL* | *matK* | *ndhF* | *psbA-trnH* | *rbcL* | *trnL-F* | *trnS-G* | *ycf1* |
| 146 | Desmos_dinhensis | *Desmos dinhensis* (Finet & Gagnep.) Merr. | - | JQ768569 | JQ768605 | JQ768648 | - | JQ768729 | - | - |
| 147 | Desmos_dumosus | *Desmos dumosus* (Roxb.) Saff. | - | JQ768570 | JQ768606 | JQ768649 | JQ768689 | JQ768730 | - | - |
| 148 | Desmos_elegans | *Desmos elegans* (Thwaites) Saff. | - | JQ768571 | - | JQ768650 | JQ768690 | JQ768731 | - | - |
| 149 | Desmos_goezeanus | *Desmos goezeanus* (F. Muell.) Jessup | - | JQ768572 | JQ768607 | JQ768651 | JQ768691 | JQ768732 | - | - |
| 150 | Desmos_polycarpus | *Desmos polycarpus* Jessup | - | KX786589 | KX786607 | KX786613 | - | KX786630 | | |
| 151 | Desmos_sp | *Desmos* sp. | - | JQ768573 | - | JQ768652 | JQ768692 | JQ768733 | | |
| 152 | Desmos_wardianus | *Desmos wardianus* (F. M. Bailey) Jessup | - | JQ768574 | JQ768608 | JQ768653 | JQ768693 | JQ768734 | - | - |
| 153 | Diclinanona_tessmannii | *Diclinanona tessmannii* Diels | - | DQ125056 | - | - | AY841611 | AY841689 | - | - |
| 154 | Dielsiothamnus_divaricatus | *Dielsiothamnus divaricatus* (Diels) R. E. Fr. | - | - | - | EU169736 | EU169759 | EU169781 | EU169803 | - |
| 155 | Disepalum_aciculare | *Disepalum aciculare* D. M. Johnson | - | KT452821 | KT452832 | - | - | KT452843 | - | KT452854 |
| 156 | Disepalum_anomalum | *Disepalum anomalum* Hook. f. | - | KT452819 | KT452830 | - | - | KT452841 | - | KT452852 |
| 157 | Disepalum_coronatum | *Disepalum coronatum* Becc. | - | KT452820 | KT452831 | - | - | KT452842 | - | KT452853 |
| 158 | Disepalum_longipes | *Disepalum longipes* King | - | KT452822 | KT452833 | - | - | KT452844 | - | KT452855 |
| 159 | Disepalum_petelotii | *Disepalum petelotii* (Merr.) D. M. Johnson | - | KT452813 | KT452824 | - | - | KT452835 | - | KT452846 |
| 160 | Disepalum_plagioneurum | *Disepalum plagioneurum* (Diels) D. M. Johnson | - | KT452814 | KT452825 | - | - | KT452836 | - | KT452847 |
| 161 | Disepalum_platypetalum | *Disepalum platypetalum* Merr. | EF179257 | DQ125057 | EF179292 | DQ125122 | - | - | EF179334 | - |
| 162 | Disepalum_pulchrum | *Disepalum pulchrum* (King) J. Sinclair | - | GQ139736 | - | GQ139875 | JQ513888 | GQ139909 | - | GU937383 |

| No. | Voucher information | | GenBank accession numbers | | | | | | | |
|---|---|---|---|---|---|---|---|---|---|---|
| | Tip name in the tree | Taxa name | *atpB-rbcL* | *matK* | *ndhF* | *psbA-trnH* | *rbcL* | *trnL-F* | *trnS-G* | *ycf1* |
| 163 | Drepananthus_apoensis | *Drepananthus apoensis* Elmer | - | HM173721 | - | HM173692 | HM173778 | HM173750 | - | - |
| 164 | Drepananthus_biovulatus | *Drepananthus biovulatus* (Boerl.) Survesw. & R. M. K. Saunders | - | HM173722 | - | HM173693 | AY841604 | AY841682 | - | - |
| 165 | Drepananthus_cauliflorus | *Drepananthus cauliflorus* (Lauterb. & K. Schum.) Survesw. & R. M. K. Saunders | - | HM173723 | - | HM173694 | HM173780 | HM173752 | - | - |
| 166 | Drepananthus_deltoideus | *Drepananthus deltoideus* (Airy Shaw) Survesw. & R. M. K. Saunders | - | HM173724 | - | HM173695 | HM173781 | HM173753 | - | - |
| 167 | Drepananthus_havilandii | *Drepananthus havilandii* (Boerl.) Survesw. & R. M. K. Saunders | - | HM173727 | - | HM173698 | HM173784 | HM173756 | - | - |
| 168 | Drepananthus_hexagynus | *Drepananthus hexagynus* (Miq.) Survesw. & R. M. K. Saunders | - | HM173740 | - | HM173711 | HM173797 | HM173769 | - | - |
| 169 | Drepananthus_kingii | *Drepananthus kingii* (Boerl. ex Koord.) Survesw. & R. M. K. Saunders | - | HM173728 | - | HM173699 | HM173785 | HM173757 | - | - |
| 170 | Drepananthus_magnificus | *Drepananthus magnificus* (Diels) Survesw. & R. M. K. Saunders | - | HM173729 | - | HM173700 | HM173786 | HM173758 | - | - |
| 171 | Drepananthus_obtusifolius | *Drepananthus obtusifolius* (Becc. & Scheff.) Survesw. & R. M. K. Saunders | - | HM173732 | - | HM173703 | HM173789 | HM173761 | - | - |
| 172 | Drepananthus_pahangensis | *Drepananthus pahangensis* M. R. Hend. | - | HM173741 | - | HM173712 | HM173798 | - | - | - |
| 173 | Drepananthus_petiolatus | *Drepananthus petiolatus* (Diels) Survesw. & R. M. K. Saunders | - | HM173743 | - | HM173714 | HM173800 | HM173771 | - | - |

| No. | Voucher information | | GenBank accession numbers | | | | | | | |
|---|---|---|---|---|---|---|---|---|---|---|
| | Tip name in the tree | Taxa name | *atpB-rbcL* | *matK* | *ndhF* | *psbA-trnH* | *rbcL* | *trnL-F* | *trnS-G* | *ycf1* |
| 174 | Drepananthus_polycarpus | *Drepananthus polycarpus* (C. T. White & W. D. Francis) Survesw. & R. M. K. Saunders | - | HM173733 | - | HM173704 | HM173790 | HM173762 | - | - |
| 175 | Drepananthus_pruniferus | *Drepananthus pruniferus* Maingay ex Hook. f. & Thomson | - | HM173734 | - | HM173705 | HM173791 | HM173763 | - | - |
| 176 | Drepananthus_pubescens | *Drepananthus pubescens* (Scheff.) Survesw. & R. M. K. Saunders | - | HM173735 | - | HM173706 | HM173792 | HM173764 | - | - |
| 177 | Drepananthus_ridleyi | *Drepananthus ridleyi* (King) Survesw. & R. M. K. Saunders | - | HM173736 | - | HM173707 | HM173793 | HM173765 | - | - |
| 178 | Drepananthus_vitiensis | *Drepananthus vitiensis* (A. C. Sm.) Survesw. & R. M. K. Saunders | - | HM173738 | - | HM173709 | HM173795 | HM173767 | - | - |
| 179 | Duckeanthus_grandiflorus | *Duckeanthus grandiflorus* R. E. Fr. | MF322674 | MF322636 | - | - | MF322651 | MF322664 | - | - |
| 180 | Duguetia_bahiensis | *Duguetia bahiensis* Maas | - | AY740532 | - | - | AY738152 | AY740564 | - | - |
| 181 | Duguetia_cadaverica | *Duguetia cadaverica* Huber | - | AY740533 | - | - | AY738153 | AY740565 | - | - |
| 182 | Duguetia_calycina | *Duguetia calycina* Benoist | - | AY740534 | - | - | AY738154 | AY740566 | - | - |
| 183 | Duguetia_cauliflora | *Duguetia cauliflora* R. E. Fr. | - | AY740535 | - | - | AY738155 | AY740567 | - | - |
| 184 | Duguetia_chrysea | *Duguetia chrysea* Maas | - | AY740536 | - | AY841435 | AY738156 | AY841691 | - | - |
| 185 | Duguetia_confinis | *Duguetia confinis* (Engl. & Diels) Chatrou | - | AY740537 | - | - | AY738157 | AY740569 | - | - |
| 186 | Duguetia_confusa | *Duguetia confusa* Maas | - | AY740538 | - | - | AY738158 | AY740570 | - | - |
| 187 | Duguetia_echinophora | *Duguetia echinophora* R. E. Fr. | - | AY740539 | - | - | AY738159 | AY740571 | - | - |
| 188 | Duguetia_guianensis | *Duguetia guianensis* R. E. Fr. | - | AY740540 | - | - | AY738160 | AY740572 | - | - |
| 189 | Duguetia_hadrantha | *Duguetia hadrantha* (Diels) R. E. Fr. | EF179258 | AY740541 | EF179293 | DQ125123 | AY738161 | AY740573 | EF179335 | - |

| No. | Voucher information | | GenBank accession numbers | | | | | | | |
|---|---|---|---|---|---|---|---|---|---|---|
| | Tip name in the tree | Taxa name | *atpB-rbcL* | *matK* | *ndhF* | *psbA-trnH* | *rbcL* | *trnL-F* | *trnS-G* | *ycf1* |
| 190 | Duguetia_lucida | *Duguetia lucida* Urb. | - | AY740542 | - | - | AY738162 | AY740574 | - | - |
| 191 | Duguetia_macrocalyx | *Duguetia macrocalyx* R. E. Fr. | - | AY740543 | - | - | AY738163 | AY740575 | - | - |
| 192 | Duguetia_macrophylla | *Duguetia macrophylla* R. E. Fr. | - | AY740544 | - | - | AY738164 | AY740576 | - | - |
| 193 | Duguetia_marcgraviana | *Duguetia marcgraviana* Mart. | - | AY740545 | - | - | AY738165 | AY740577 | - | - |
| 194 | Duguetia_megalocarpa | *Duguetia megalocarpa* Maas | - | AY740546 | - | - | AY738166 | AY740578 | - | - |
| 195 | Duguetia_moricandiana | *Duguetia moricandiana* Mart. | - | AY740547 | - | - | AY738167 | AY740579 | - | - |
| 196 | Duguetia_neglecta | *Duguetia neglecta* Sandwith | - | AY740548 | - | - | AY738168 | AY740580 | - | - |
| 197 | Duguetia_odorata | *Duguetia odorata* (Diels) J. F. Macbr. | - | AY740549 | - | - | AY738169 | AY740581 | - | - |
| 198 | Duguetia_panamensis | *Duguetia panamensis* Standl. | - | AY740550 | - | - | AY738170 | AY740582 | - | - |
| 199 | Duguetia_peruviana | *Duguetia peruviana* (R. E. Fr.) J. F. Macbr. | - | AY740551 | - | - | AY738171 | AY740583 | - | - |
| 200 | Duguetia_pycnastera | *Duguetia pycnastera* Sandwith | - | AY740552 | - | - | AY738172 | AY740584 | - | - |
| 201 | Duguetia_quitarensis | *Duguetia quitarensis* Benth. | - | AY740553 | - | - | AY738173 | AY740585 | - | - |
| 202 | Duguetia_riedeliana | *Duguetia riedeliana* R. E. Fr. | - | AY740554 | - | - | AY738174 | AY740586 | - | - |
| 203 | Duguetia_salicifolia | *Duguetia salicifolia* R. E. Fr. | - | AY740555 | - | - | AY738175 | AY740587 | - | - |
| 204 | Duguetia_sessilis | *Duguetia sessilis* (Vell.) Maas | - | AY740556 | - | - | AY738176 | AY740588 | - | - |
| 205 | Duguetia_sooretamae | *Duguetia sooretamae* Maas | - | AY740557 | - | DQ861746 | AY738177 | AY740589 | - | - |
| 206 | Duguetia_staudtii | *Duguetia staudtii* (Engl. & Diels) Chatrou | EF179259 | AY740558 | EF179294 | DQ125124 | AY738178 | AY740590 | EF179336 | - |
| 207 | Duguetia_stelechantha | *Duguetia stelechantha* (Diels) R. E. Fr. | - | AY740559 | - | - | AY738179 | AY740591 | - | - |
| 208 | Duguetia_surinamensis | *Duguetia surinamensis* R. E. Fr. | - | AY740560 | - | FJ038842 | AY738180 | AY740592 | - | - |
| 209 | Duguetia_ulei | *Duguetia ulei* (Diels) R. E. Fr. | - | AY740561 | - | - | AY738181 | AY740593 | - | - |

| No. | Voucher information | | GenBank accession numbers | | | | | | | |
|---|---|---|---|---|---|---|---|---|---|---|
| | Tip name in the tree | Taxa name | *atpB-rbcL* | *matK* | *ndhF* | *psbA-trnH* | *rbcL* | *trnL-F* | *trnS-G* | *ycf1* |
| 210 | Duguetia_uniflora | *Duguetia uniflora* (DC.) Mart. | - | AY740562 | - | - | AY738182 | AY740594 | - | - |
| 211 | Duguetia_yeshidan | *Duguetia yeshidan* Sandwith | - | AY740563 | - | - | AY738183 | AY740595 | - | - |
| 212 | Ephedranthus_boliviensis | *Ephedranthus boliviensis* Chatrou & Pirie | - | - | - | - | AY841614 | AY841692 | - | - |
| 213 | Ephedranthus_parviflorus | *Ephedranthus parviflorus* S. Moore | - | - | - | AY841462 | AY841615 | AY841693 | - | - |
| 214 | Ephedranthus_sp_Maas_8826 | *Ephedranthus sp Maas et al 8826* | AY841376 | AY841396 | AY841407 | AY841463 | AY841616 | AY841694 | AY841554 | - |
| 215 | Eupomatia_bennettii | *Eupomatia bennettii* F.Müll. | - | JQ437547 | - | - | DQ861790 | DQ861842 | JQ513885 | - |
| 216 | Fenerivia_angustielliptica | *Fenerivia angustielliptica* (G. E. Schatz & Le Thomas) R. M. K. Saunders | - | JF810373 | - | - | JF810385 | JF810397 | - | - |
| 217 | Fenerivia_capuronii | *Fenerivia capuronii* (Cavaco & Keraudren) R. M. K. Saunders | - | JF810374 | - | - | JF810386 | JF810398 | - | - |
| 218 | Fenerivia_chapelieri | *Fenerivia chapelieri* (Baill.) R. M. K. Saunders | - | JF810375 | JQ723788 | - | JF810387 | JF810399 | - | - |
| 219 | Fenerivia_emarginata | *Fenerivia emarginata* (Diels) R. M. K. Saunders | - | JF810376 | - | - | JF810388 | JF810400 | - | - |
| 220 | Fenerivia_ghesquiereana | *Fenerivia ghesquiereana* (Cavaco & Keraudren) R. M. K. Saunders | - | JF810377 | - | - | JF810389 | JF810401 | - | - |
| 221 | Fenerivia_heteropetala | *Fenerivia heteropetala* Diels | - | JF810378 | - | - | JF810390 | JF810402 | - | - |
| 222 | Fenerivia_humbertii | *Fenerivia humbertii* (Diels) R. M. K. Saunders | - | JF810379 | - | - | JF810391 | JF810403 | - | - |
| 223 | Fenerivia_madagascariensis | *Fenerivia madagascariensis* (Cavaco & Keraudren) R. M. K. Saunders | - | JF810380 | - | - | JF810392 | JF810404 | - | - |

| No. | Voucher information | | GenBank accession numbers | | | | | | | |
|---|---|---|---|---|---|---|---|---|---|---|
| | Tip name in the tree | Taxa name | *atpB-rbcL* | *matK* | *ndhF* | *psbA-trnH* | *rbcL* | *trnL-F* | *trnS-G* | *ycf1* |
| 224 | Fenerivia_oligosperma | *Fenerivia oligosperma* (Danguy) R. M. K. Saunders | - | JF810381 | - | - | JF810393 | JF810405 | - | - |
| 225 | Fenerivia_richardiana | *Fenerivia richardiana* (Baill.) R. M. K. Saunders | - | JF810382 | - | - | JF810394 | JF810406 | - | - |
| 226 | Fissistigma_glaucescens | *Fissistigma glaucescens* (Hance) Merr. | - | AY743482 | - | - | AY743444 | AY743463 | - | - |
| 227 | Fissistigma_polyanthoide s | *Fissistigma polyanthoides* (Aug. DC.) Merr. | - | JQ768575 | JQ768609 | JQ768654 | JQ768694 | JQ768735 | - | - |
| 228 | Fissistigma_uonicum | *Fissistigma uonicum* (Dunn) Merr. | - | - | - | - | AY841617 | AY841695 | - | - |
| 229 | Friesodielsia_affinis | *Friesodielsia affinis* (Hook.f. & Thomson) D.Das | - | KX786590 | - | KX786614 | - | KX786631 | | |
| 230 | Friesodielsia_bakeri | *Friesodielsia bakeri* (Merr.) Steenis | - | KX786591 | - | KX786615 | - | KX786632 | | |
| 231 | Friesodielsia_biglandulos a | *Friesodielsia biglandulosa* (Blume) Steenis | - | KX786592 | JQ768610 | JQ768655 | - | JQ768736 | | |
| 232 | Friesodielsia_boreensis | *Friesodielsia borneensis* (Miq.) Steenis | - | KX786593 | KX786608 | KX786616 | - | KX786633 | | |
| 233 | Friesodielsia_calycina | *Friesodielsia calycina* (King) Steenis | - | KX786594 | - | KX786617 | - | KX786634 | | |
| 234 | Friesodielsia_cuneiformis | *Friesodielsia cuneiformis* (Blume) Steenis | - | JQ768576 | JQ768611 | - | JQ768695 | JQ768737 | | |
| 235 | Friesodielsia_desmoides | *Friesodielsia desmoides* (Craib) Steenis | - | JQ768577 | JQ768612 | JQ768656 | JQ768696 | JQ768738 | | |
| 236 | Friesodielsia_filipes | *Friesodielsia filipes* (Hook.f. & Thomson) Steenis | - | KX786596 | - | - | - | KX786636 | | |
| 237 | Friesodielsia_fornicata | *Friesodielsia fornicata* (Roxb.) D.Das | - | KX786597 | - | KX786618 | - | KX786637 | | |

| No. | Voucher information | | GenBank accession numbers | | | | | | | |
|---|---|---|---|---|---|---|---|---|---|---|
| | Tip name in the tree | Taxa name | *atpB-rbcL* | *matK* | *ndhF* | *psbA-trnH* | *rbcL* | *trnL-F* | *trnS-G* | *ycf1* |
| 238 | Friesodielsia_glauca | *Friesodielsia glauca* (Hook.f. &Thomson) Steenis | - | KX786598 | - | KX786619 | - | KX786638 | | |
| 239 | Friesodielsia_kingii | *Friesodielsia kingii* (J.Sinclair) Steenis | - | KX786601 | - | - | - | KX786641 | | |
| 240 | Friesodielsia_latifolia | *Friesodielsia latifolia* (Hook.f. & Thomson) Steenis | - | KX786602 | - | KX786622 | - | KX786642 | | |
| 241 | Friesodielsia_longifolia | *Friesodielsia longiflora* (Merr.) Steenis | - | KX786603 | - | - | - | - | | |
| 242 | Friesodielsia_sahyadrica | *Friesodielsia sahyadrica* N.V.Page & S.Surveswaran | - | KC933936 | KC933942 | KC933940 | KC933934 | KC933938 | | |
| 243 | Friesodielsia_sp_12278 | *Friesodielsia* sp. 12278 | - | KX761305 | KX787025 | KX786964 | KX761336 | KX786994 | | |
| 244 | Fusaea_longifolia | *Fusaea longifolia* (Aubl.) Saff. | - | FJ514750 | - | FJ038844 | AY841620 | AY841698 | - | - |
| 245 | Fusaea_peruviana | *Fusaea peruviana* R. E. Fr. | EF179260 | AY743483 | EF179295 | AY841436 | AY743445 | AY743464 | EF179337 | - |
| 246 | Galbulimima_belgraveana | *Galbulimima belgraveana* (F.Muell.) Sprague | - | AY220441 | AY218176 | - | L12646 | AY220415 | | |
| 247 | Goniothalamus_amuyon | *Goniothalamus amuyon* (Blanco) Merr. | KM818567 | KM818518 | KM818648 | KM818728 | KM818839 | KM818898 | KM818916 | KM818979 |
| 248 | Goniothalamus_andersonii | *Goniothalamus andersonii* J. Sinclair | KM818519 | KM818568 | - | KM818711 | KM818789 | KM818867 | KM818949 | - |
| 249 | Goniothalamus_aruensis | *Goniothalamus angustifolius* (A. C. Sm.) B. Xue & R. M. K. Saunders | - | KM818569 | KM818632 | KM818732 | KM818797 | KM818878 | KM818937 | KM818983 |
| 250 | Goniothalamus_augustifolia | *Goniothalamus aruensis* Scheff. | KM818520 | KM818570 | KM818640 | KM818706 | KM818791 | KM818868 | KM818918 | - |
| 251 | Goniothalamus_australis | *Goniothalamus australis* Jessup | KM818521 | KM818571 | KM818638 | KM818709 | KM818836 | KM818887 | KM818910 | KM818973 |

| No. | Voucher information | | GenBank accession numbers | | | | | | | |
|---|---|---|---|---|---|---|---|---|---|---|
| | Tip name in the tree | Taxa name | *atpB-rbcL* | *matK* | *ndhF* | *psbA-trnH* | *rbcL* | *trnL-F* | *trnS-G* | *ycf1* |
| 252 | Goniothalamus_borneensis | *Goniothalamus borneensis* Mat-Salleh | KM818522 | KM818572 | KM818673 | - | KM818826 | KM818893 | KM818952 | - |
| 253 | Goniothalamus_bracteosus | *Goniothalamus bracteosus* Bân | - | KM818573 | - | KM818730 | KM818796 | KM818906 | KM818967 | - |
| 254 | Goniothalamus_calcareus | *Goniothalamus calcareus* Mat-Salleh | - | - | - | KM818717 | KM818810 | - | KM818927 | KM818994 |
| 255 | Goniothalamus_calvicarpus | *Goniothalamus calvicarpus* Craib | KM818523 | KM818574 | KM818647 | KM818702 | KM818809 | KM818874 | KM818934 | KM819005 |
| 256 | Goniothalamus_cardiopetalus | *Goniothalamus cardiopetalus* (Dalzell) Hook. f. & Thomson | KM818524 | KM818575 | KM818654 | KM818692 | KM818799 | KM818879 | KM818912 | - |
| 257 | Goniothalamus_cauiflorus | *Goniothalamus cauliflorus* K. Schum. | KM818525 | KM818576 | KM818663 | KM818696 | KM818807 | KM818869 | KM818919 | - |
| 258 | Goniothalamus_cheliensis | *Goniothalamus cheliensis* Hu | KM818526 | KM818577 | KM818661 | KM818678 | KM818831 | KM818901 | KM818926 | KM818992 |
| 259 | Goniothalamus_clemensii | *Goniothalamus clemensii* Bân | - | KM818578 | - | KM818736 | KM818780 | KM818844 | KM818915 | - |
| 260 | Goniothalamus_costulatus | *Goniothalamus costulatus* Miq. | - | KM818579 | - | KM818737 | KM818805 | KM818865 | KM818945 | - |
| 261 | Goniothalamus_dumontetii | *Goniothalamus dumontetii* R. M. K. Saunders & Munzinger | - | KM818580 | - | KM818729 | KM818840 | KM818861 | KM818954 | - |
| 262 | Goniothalamus_elegans | *Goniothalamus elegans* Ast | KM818527 | KM818581 | KM818676 | KM818707 | KM818817 | KM818850 | KM818955 | KM818997 |
| 263 | Goniothalamus_elmeri | *Goniothalamus elmeri* Merr. | - | KM818582 | KM818639 | KM818677 | KM818811 | KM818882 | KM818924 | KM819003 |
| 264 | Goniothalamus_expansus | *Goniothalamus expansus* Craib | - | KM818583 | KM818634 | KM818714 | KM818829 | KM818853 | KM818931 | KM818987 |
| 265 | Goniothalamus_fasciculatus | *Goniothalamus fasciculatus* Boerl. | KM818528 | KM818584 | KM818636 | - | - | KM818890 | KM818950 | - |
| 266 | Goniothalamus_gardneri | *Goniothalamus gardneri* Hook. f. & Thomson | KM818529 | KM818585 | KM818656 | KM818704 | KM818784 | KM818871 | KM818923 | KM819001 |
| 267 | Goniothalamus_giganteus | *Goniothalamus giganteus* Wall. ex Hook. f. & Thomson | KM818530 | KM818586 | KM818655 | KM818698 | KM818837 | KM818892 | KM818963 | KM818996 |

| No. | Voucher information | | GenBank accession numbers | | | | | | | |
|---|---|---|---|---|---|---|---|---|---|---|
| | Tip name in the tree | Taxa name | *atpB-rbcL* | *matK* | *ndhF* | *psbA-trnH* | *rbcL* | *trnL-F* | *trnS-G* | *ycf1* |
| 268 | Goniothalamus_grandiflorus | *Goniothalamus grandiflorus* (Warb.) Boerl. | KM818531 | KM818587 | KM818637 | KM818691 | KM818802 | KM818851 | KM818930 | - |
| 269 | Goniothalamus_griffithii | *Goniothalamus griffithii* Hook. f. & Thomson | KM818532 | KM818588 | KM818651 | KM818701 | KM818798 | KM818894 | KM818939 | KM819000 |
| 270 | Goniothalamus_hookeri | *Goniothalamus hookeri* Thwaites | KM818533 | KM818589 | KM818657 | KM818734 | KM818814 | KM818872 | KM818956 | - |
| 271 | Goniothalamus_howii | *Goniothalamus howii* Merr. & Chun | KM818534 | KM818590 | - | KM818689 | KM818833 | KM818886 | KM818938 | KM818986 |
| 272 | Goniothalamus_imbricatus | *Goniothalamus imbricatus* (Blume) Koord. | KM818535 | KM818591 | - | KM818722 | KM818806 | KM818847 | KM818946 | KM818998 |
| 273 | Goniothalamus_kinabaluensis | *Goniothalamus kinabaluensis* Bân ex Mat-Salleh | KM818536 | KM818592 | KM818672 | KM818684 | KM818787 | KM818876 | KM818935 | - |
| 274 | Goniothalamus_laoticus | *Goniothalamus laoticus* (Fin. & Gagnep.) Tien Ban | KM818537 | KM818593 | KM818666 | KM818699 | KM818808 | KM818881 | KM818959 | KM818993 |
| 275 | Goniothalamus_loerzingii | *Goniothalamus loerzingii* R. M. K. Saunders | - | KM818594 | - | KM818724 | KM818782 | KM818902 | KM818947 | - |
| 276 | Goniothalamus_macranthus | *Goniothalamus macranthus* (Kurz) Boerl. | KM818538 | KM818595 | KM818643 | KM818695 | KM818792 | KM818873 | KM818928 | KM818995 |
| 277 | Goniothalamus_macrophyllus | *Goniothalamus macrophyllus* (Blume) Hook. f. & Thomson | KM818539 | KM818596 | KM818665 | KM818688 | KM818843 | KM818897 | KM818940 | KM819002 |
| 278 | Goniothalamus_maewongensis | *Goniothalamus maewongensis* R. M. K. Saunders & Chalermglin | KM818540 | KM818597 | KM818659 | KM818725 | KM818838 | KM818888 | KM818962 | KM818977 |
| 279 | Goniothalamus_majestatis | *Goniothalamus majestatis* Keßler | KM818541 | KM818598 | - | KM818713 | KM818788 | KM818903 | KM818958 | - |
| 280 | Goniothalamus_malayanus | *Goniothalamus malayanus* Hook. f. & Thomson | KM818542 | KM818599 | KM818650 | KM818718 | KM818835 | KM818891 | KM818914 | KM819006 |

| No. | Voucher information | | GenBank accession numbers | | | | | | | |
|---|---|---|---|---|---|---|---|---|---|---|
| | Tip name in the tree | Taxa name | *atpB-rbcL* | *matK* | *ndhF* | *psbA-trnH* | *rbcL* | *trnL-F* | *trnS-G* | *ycf1* |
| 281 | Goniothalamus_megalocalyx | *Goniothalamus megalocalyx* I. M. Turner & R. M. K. Saunders | KM818543 | KM818600 | KM818645 | KM818726 | KM818822 | KM818885 | KM818960 | KM819007 |
| 282 | Goniothalamus_monospermus | *Goniothalamus monospermus* (A. Gray) R. M. K. Saunders | - | KM818601 | - | KM818735 | KM818790 | - | KM818969 | - |
| 283 | Goniothalamus_montanus | *Goniothalamus montanus* J. Sinclair | KM818544 | KM818602 | KM818674 | KM818710 | KM818813 | KM818856 | KM818932 | - |
| 284 | Goniothalamus_obtusatus | *Goniothalamus obtusatus* (Baill.) R. M. K. Saunders | KM818545 | KM818603 | KM818660 | KM818687 | KM818815 | KM818883 | KM818911 | KM818981 |
| 285 | Goniothalamus_palawanensis | *Goniothalamus palawanensis* C.C. Tang & R. M. K. Saunders | - | KM818604 | - | KM818716 | KM818793 | KM818855 | KM818925 | KM818976 |
| 286 | Goniothalamus_parallelivenius | *Goniothalamus parallelivenius* Ridl. | KM818546 | KM818605 | KM818635 | KM818683 | KM818801 | KM818880 | KM818941 | - |
| 287 | Goniothalamus_repevensis | *Goniothalamus repevensis* Pierre ex Finet & Gagnep. | KM818547 | KM818606 | KM818664 | KM818723 | KM818795 | KM818877 | KM818936 | - |
| 288 | Goniothalamus_reticulatus | *Goniothalamus reticulatus* Thwaites | KM818548 | KM818607 | - | - | KM818786 | - | KM818913 | - |
| 289 | Goniothalamus_ridleyi | *Goniothalamus ridleyi* King | KM818549 | KM818608 | - | KM818739 | KM818830 | KM818860 | KM818951 | KM818985 |
| 290 | Goniothalamus_rotundisepalus | *Goniothalamus rotundisepalus* M. R. Hend. | KM818550 | KM818609 | KM818649 | KM818693 | KM818794 | KM818857 | KM818908 | - |
| 291 | Goniothalamus_rufus | *Goniothalamus rufus* Miq. | KM818551 | KM818610 | - | KM818727 | KM818819 | KM818848 | KM818943 | - |
| 292 | Goniothalamus_sawtehii | *Goniothalamus sawtehii* C. E. C. Fisch. | KM818552 | KM818611 | KM818646 | KM818680 | KM818785 | KM818895 | KM818942 | KM819004 |
| 293 | Goniothalamus_scortechinii | *Goniothalamus scortechinii* King | KM818553 | KM818612 | KM818670 | KM818712 | KM818781 | KM818845 | KM818929 | KM818988 |

| No. | Voucher information | | GenBank accession numbers | | | | | | | |
|---|---|---|---|---|---|---|---|---|---|---|
| | Tip name in the tree | Taxa name | *atpB-rbcL* | *matK* | *ndhF* | *psbA-trnH* | *rbcL* | *trnL-F* | *trnS-G* | *ycf1* |
| 294 | Goniothalamus_sesquipedalis | *Goniothalamus sesquipedalis* (Wall.) Hook. f. & Thomson | KM818554 | KM818613 | KM818667 | KM818719 | KM818825 | KM818904 | KM818907 | KM818984 |
| 295 | Goniothalamus_sp_TCC10 | *Goniothalamus* sp. tcc10 | - | KM818614 | KM818675 | KM818715 | KM818821 | KM818864 | KM818944 | KM818980 |
| 296 | Goniothalamus_suaveness_TCC32 | *Goniothalamus suaveolens* Becc. 466 | - | KM818615 | - | KM818681 | KM818818 | KM818884 | KM818968 | KM818999 |
| 297 | Goniothalamus_suaveolens_466 | *Goniothalamus suaveolens* Becc. tcc32 | KM818555 | KM818616 | - | KM818682 | KM818800 | KM818858 | KM818933 | KM818982 |
| 298 | Goniothalamus_tamirensis | *Goniothalamus tamirensis* Pierre ex Finet & Gagnep. | KM818556 | KM818617 | KM818662 | KM818700 | KM818832 | KM818866 | KM818917 | KM818990 |
| 299 | Goniothalamus_tapis | *Goniothalamus tapis* Miq. | EF179262 | DQ125058 | EF179297 | AY841622 | AY841700 | EF179339 | - | AY841622 |
| 300 | Goniothalamus_tapisoides | *Goniothalamus tapisoides* Mat-Salleh | KM818557 | KM818618 | KM818641 | KM818686 | KM818823 | KM818899 | KM818920 | - |
| 301 | Goniothalamus_tavoyensis | *Goniothalamus tavoyensis* Chatterjee | KM818558 | KM818619 | KM818633 | KM818690 | KM818841 | KM818854 | KM818961 | - |
| 302 | Goniothalamus_tenuifolius | *Goniothalamus tenuifolius* King | KM818559 | KM818620 | KM818669 | KM818694 | KM818842 | KM818889 | KM818909 | KM818974 |
| 303 | Goniothalamus_thomsoni | *Goniothalamus thomsonii* Thwaites | - | KM818621 | - | KM818733 | KM818834 | KM818875 | KM818971 | - |
| 304 | Goniothalamus_thwaitesii | *Goniothalamus thwaitesii* Hook. f. & Thomson | KM818560 | KM818622 | KM818653 | KM818703 | - | KM818849 | KM818922 | - |
| 305 | Goniothalamus_tomentosus | *Goniothalamus tomentosus* R. M. K. Saunders | KM818561 | KM818623 | - | KM818738 | KM818783 | KM818846 | KM818964 | - |
| 306 | Goniothalamus_tortilipetalus | *Goniothalamus tortilipetalus* M. R. Hend. | - | KM818624 | KM818642 | KM818708 | KM818828 | KM818905 | KM818948 | - |

| No. | Voucher information | | GenBank accession numbers | | | | | | | |
|---|---|---|---|---|---|---|---|---|---|---|
| | Tip name in the tree | Taxa name | *atpB-rbcL* | *matK* | *ndhF* | *psbA-trnH* | *rbcL* | *trnL-F* | *trnS-G* | *ycf1* |
| 307 | Goniothalamus_touranensis | *Goniothalamus touranensis* Ast | - | KM818625 | - | KM818731 | KM818804 | KM818870 | KM818965 | - |
| 308 | Goniothalamus_undulatus | *Goniothalamus undulatus* Ridl. | KM818562 | KM818626 | KM818652 | KM818679 | KM818820 | KM818896 | KM818921 | KM818978 |
| 309 | Goniothalamus_uvarioides | *Goniothalamus uvarioides* King | - | KM818627 | KM818658 | KM818685 | KM818827 | KM818852 | KM818966 | KM818975 |
| 310 | Goniothalamus_velutinus | *Goniothalamus velutinus* Airy Shaw | KM818563 | KM818628 | KM818644 | KM818705 | KM818812 | KM818900 | KM818953 | KM818989 |
| 311 | Goniothalamus_woodii | *Goniothalamus woodii* Merr. ex Mat Salleh | KM818564 | KM818629 | KM818668 | KM818720 | KM818824 | KM818862 | KM818972 | - |
| 312 | Goniothalamus_wrayi | *Goniothalamus wrayi* King | KM818565 | KM818630 | KM818671 | KM818721 | KM818803 | KM818859 | KM818957 | - |
| 313 | Goniothalamus_wynaadensis | *Goniothalamus wynaadensis* (Bedd.) Bedd. | KM818566 | KM818631 | - | KM818697 | KM818816 | KM818863 | KM818970 | KM818991 |
| 314 | Greenwayodendron_oliveri | *Greenwayodendron oliveri* (Engl.) Verdc. | AY841377 | AY743489 | AY841408 | AY841465 | AY743451 | AY743470 | AY841555 | - |
| 315 | Greenwayodendron_suaveolens | *Greenwayodendron suaveolens* (Engl. & Diels) Verdc. | - | - | - | AY841466 | AY841524 | AY319090 | - | - |
| 316 | Guatteria_aeruginosa | *Guatteria aeruginosa* Standl. | EF179264 | AY740909 | EF179299 | DQ125136 | AY740958 | AY741007 | EF179341 | - |
| 317 | Guatteria_alata | *Guatteria alata* Maas & van Setten | - | AY740910 | - | DQ125137 | AY740959 | AY741008 | - | - |
| 318 | Guatteria_allenii | *Guatteria allenii* R. E. Fr. | - | AY740911 | - | DQ125138 | DQ861791 | DQ861843 | - | - |
| 319 | Guatteria_alta | *Guatteria alta* R. E. Fr. | - | DQ125065 | - | DQ125139 | DQ124941 | DQ124999 | - | - |
| 320 | Guatteria_hirsuta_2 | *Guatteria alutacea* Diels = *Guatteria hirsuta* Ruiz & Pav. | - | AY740912 | - | DQ125140 | AY740961 | AY741010 | - | - |
| 321 | Guatteria_amplifolia_1 | *Guatteria amplifolia* Triana & Planch. | - | DQ125066 | - | DQ125141 | DQ124942 | DQ125000 | - | - |
| 322 | Guatteria_anomala | *Guatteria anomala* R. E. Fr. | EF179263 | AY740913 | EF179298 | AY841437 | AY740962 | AY741011 | EF179340 | - |

| No. | Voucher information | | GenBank accession numbers | | | | | | | |
|---|---|---|---|---|---|---|---|---|---|---|
| | Tip name in the tree | Taxa name | *atpB-rbcL* | *matK* | *ndhF* | *psbA-trnH* | *rbcL* | *trnL-F* | *trnS-G* | *ycf1* |
| 323 | Guatteria_liesneri_2 | *Guatteria anthracina* Scharf & Maas = *Guatteria liesneri* D. M. Johnson & N. A. Murray | - | DQ861698 | - | DQ861748 | DQ861792 | DQ861844 | - | - |
| 324 | Guatteria_atra | *Guatteria atra* Sandwith = *Guatteria punctata* (Aubl.) R. A. Howard | - | AY740914 | - | DQ125142 | AY740963 | AY741012 | - | - |
| 325 | Guatteria_australis_1 | *Guatteria australis* A. St.-Hil. | - | AY740915 | - | - | AY740964 | AY741013 | - | - |
| 326 | Guatteria_blainii | *Guatteria blainii* (Griseb.) Urb. | - | AY740916 | - | DQ861758 | DQ861793 | DQ861845 | - | - |
| 327 | Guatteria_blepharophylla _1 | *Guatteria blepharophylla* Mart. | - | DQ861738 | - | DQ861782 | DQ861834 | DQ861888 | - | - |
| 328 | Guatteria_ucayalina_2 | *Guatteria boliviana* H. J. P. Winkl. = *Guatteria ucayalina* Huber | - | DQ125067 | - | DQ125144 | DQ124943 | DQ125001 | - | - |
| 329 | Guatteria_blepharophylla _2 | *Guatteria brevicuspis* R. E. Fr. = *Guatteria blepharophylla* Mart. | - | AY740917 | - | DQ125145 | AY740966 | AY741015 | - | - |
| 330 | Guatteria_hirsuta_1 | *Guatteria brevipedicellata* R. E. Fr. = *Guatteria hirsuta* Ruiz & Pav. | - | DQ125068 | - | DQ125146 | DQ124944 | DQ125002 | - | - |
| 331 | Guatteria_ferruginea_2 | *Guatteria burchellii* R. E. Fr. = *Guatteria ferruginea* A. St. -Hil. | - | DQ861701 | - | - | DQ861795 | DQ861847 | - | - |
| 332 | Guatteria_caribaea | *Guatteria caribaea* Urb. | - | AY740918 | - | DQ125149 | AY740967 | AY741016 | - | - |
| 333 | Guatteria_chiriquiensis | *Guatteria chiriquiensis* R. E. Fr. | - | AY740919 | - | DQ125150 | AY740968 | AY741017 | - | - |
| 334 | Guatteria_citriodora_1 | *Guatteria citriodora* Ducke | - | DQ861702 | - | - | DQ861796 | DQ861848 | - | - |
| 335 | Guatteria_decurrens_2 | *Guatteria decurrens* R. E. Fr. | - | DQ861703 | - | - | DQ861798 | DQ861850 | - | - |
| 336 | Guatteria_amplifolia_2 | *Guatteria diospyroides* Baill. = *Guatteria amplifolia* Triana & Planch. | - | AY740920 | - | EF179243 | AY740969 | AY741018 | - | - |

| No. | Voucher information | | GenBank accession numbers | | | | | | | |
|---|---|---|---|---|---|---|---|---|---|---|
| | Tip name in the tree | Taxa name | *atpB-rbcL* | *matK* | *ndhF* | *psbA-trnH* | *rbcL* | *trnL-F* | *trnS-G* | *ycf1* |
| 337 | Guatteria_discolor | *Guatteria discolor* R. E. Fr. | - | AY740921 | - | DQ125153 | AY740970 | AY741019 | - | - |
| 338 | Guatteria_dolichopoda_1 | *Guatteria dolichopoda* Donn. Sm. | - | DQ861704 | - | DQ861752 | DQ861800 | DQ861852 | - | - |
| 339 | Guatteria_lucens | *Guatteria dumetorum* R. E. Fr. = *Guatteria lucens* Standl. | - | AY740922 | - | DQ125154 | DQ861799 | DQ861851 | - | - |
| 340 | Guatteria_australis_9 | *Guatteria dusenii* R. E. Fr. = *Guatteria australis* A. St. -Hil. | - | DQ125072 | - | DQ125155 | DQ124948 | DQ125006 | - | - |
| 341 | Guatteria_hirsuta_4 | *Guatteria ecuadorensis* R. E. Fr. = *Guatteria hirsuta* Ruiz & Pav. | - | DQ861706 | - | DQ861753 | DQ861802 | DQ861854 | - | - |
| 342 | Guatteria_elata | *Guatteria elata* R. E. Fr. | - | AY740923 | - | DQ125156 | AY740972 | AY741021 | - | - |
| 343 | Guatteria_elegantissima | *Guatteria elegantissima* R. E. Fr. | - | AY740924 | - | DQ125157 | AY740973 | AY741022 | - | - |
| 344 | Guatteria_guianensis_2 | *Guatteria excellens R. E. Fr.* = *Guatteria guianensis* (Aubl.) R. E. Fr. | - | DQ861707 | - | DQ861754 | DQ861803 | DQ861855 | - | - |
| 345 | Guatteria_ferruginea_1 | *Guatteria ferruginea* A. St.-Hil. | - | DQ125073 | - | DQ125158 | DQ124949 | DQ125007 | - | - |
| 346 | Guatteria_foliosa | *Guatteria foliosa* Benth. | - | AY740925 | - | DQ125159 | AY740974 | AY741023 | - | - |
| 347 | Guatteria_friesiana | *Guatteria friesiana* (W. A. Rodrigues) Erkens & Maas | - | DQ861740 | - | DQ861784 | DQ861836 | DQ861889 | - | - |
| 348 | Guatteria_galeottiana | *Guatteria galeottiana* Baill. | - | DQ125074 | - | DQ125160 | DQ124950 | DQ125008 | - | - |
| 349 | Guatteria_australis_5 | *Guatteria glabrescens* R. E. Fr. = *Guatteria australis* A. St. -Hil. | EF179265 | AY740926 | EF179300 | DQ125161 | AY740975 | AY741024 | EF179342 | - |
| 350 | Guatteria_punctata_2 | *Guatteria gracilipes* R. E. Fr. = *Guatteria punctata* (Aubl.) R. A. Howard | - | DQ125075 | - | DQ125162 | DQ124951 | DQ125009 | - | - |
| 351 | Guatteria_guianensis_1 | *Guatteria guianensis* (Aubl.) R. E. Fr. | - | AY740927 | - | DQ125163 | AY740976 | AY741025 | - | - |

| No. | Voucher information | | GenBank accession numbers | | | | | | | |
|---|---|---|---|---|---|---|---|---|---|---|
| | Tip name in the tree | Taxa name | *atpB-rbcL* | *matK* | *ndhF* | *psbA-trnH* | *rbcL* | *trnL-F* | *trnS-G* | *ycf1* |
| 352 | Guatteria_heteropetala | *Guatteria heteropetala* Benth. [as *Heteropetalum spruceanum* in GenBank] | - | DQ861745 | - | DQ861789 | DQ861841 | DQ861894 | - | - |
| 353 | Guatteria_argentea | *Guatteria heterotricha* R. E. Fr. = *Guatteria argentea* Erkens & Maas | - | AY740928 | - | DQ125164 | AY740977 | AY741026 | - | - |
| 354 | Guatteria_australis_3 | *Guatteria hilariana* var. *verruculosa* R. E. Fr. = *Guatteria australis* A.St.-Hil. | - | DQ861710 | - | DQ861756 | DQ861806 | DQ861858 | - | - |
| 355 | Guatteria_hispida | *Guatteria hispida* (R. E. Fr.) Erkens & Maas | - | DQ861741 | - | DQ861785 | DQ861837 | DQ861890 | - | - |
| 356 | Guatteria_scytophylla_2 | *Guatteria hyposericea* Diels = *Guatteria scytophylla* Diels | - | AY740929 | - | DQ125166 | AY740978 | AY741027 | - | - |
| 357 | Guatteria_amplifolia_3 | *Guatteria inuncta* R. E. Fr. = *Guatteria amplifolia* Triana & Planch. | - | AY740930 | - | DQ125167 | AY740979 | AY741028 | - | - |
| 358 | Guatteria_inundata_1 | *Guatteria inundata* Mart. | - | AY740931 | - | DQ125168 | AY740980 | AY741029 | - | - |
| 359 | Guatteria_jefensis | *Guatteria jefensis* Barringer | - | AY740932 | - | DQ125169 | DQ861805 | DQ861857 | - | - |
| 360 | Guatteria_hirsuta_3 | *Guatteria juruensis* Diels = *Guatteria hirsuta* Ruiz & Pav. | - | DQ861708 | - | - | DQ861804 | DQ861856 | - | - |
| 361 | Guatteria_cryandra | *Guatteria kuhlmannii* R. E. Fr. = *Guatteria cryandra* Erkens & Maas | - | DQ861742 | - | DQ861786 | DQ861838 | DQ861891 | - | - |
| 362 | Guatteria_latifolia | *Guatteria latifolia* (Mart.) R. E. Fr. | - | AY740933 | - | DQ125170 | AY740982 | AY741031 | - | - |
| 363 | Guatteria_latisepala | *Guatteria latisepala* R. E. Fr. | - | DQ125077 | - | DQ125171 | DQ124953 | DQ125011 | - | - |

| No. | Voucher information | | GenBank accession numbers | | | | | | | |
|---|---|---|---|---|---|---|---|---|---|---|
| | Tip name in the tree | Taxa name | *atpB-rbcL* | *matK* | *ndhF* | *psbA-trnH* | *rbcL* | *trnL-F* | *trnS-G* | *ycf1* |
| 364 | Guatteria_goudotiana | *Guatteria lehmannii* R. E. Fr. = *Guatteria goudotiana* Triana & Planch. | - | DQ861711 | - | DQ861757 | DQ861807 | DQ861859 | - | - |
| 365 | Guatteria_liesneri_1 | *Guatteria liesneri* D. M. Johnson & N. A. Murray | - | AY740934 | - | DQ125172 | AY740983 | DQ861860 | - | - |
| 366 | Guatteria_macropus | *Guatteria macropus* Mart. | - | AY740935 | - | DQ125174 | AY740984 | AY741033 | - | - |
| 367 | Guatteria_maypurensis | *Guatteria maypurensis* Kunth | - | AY740936 | - | DQ125175 | AY740985 | DQ861882 | - | - |
| 368 | Guatteria_megalophylla | *Guatteria megalophylla* Diels | - | AY740937 | - | DQ125176 | AY740986 | AY741035 | - | - |
| 369 | Guatteria_sellowiana_2 | *Guatteria mexiae* R. E. Fr. = *Guatteria sellowiana* Schltdl. | - | DQ125080 | - | DQ125178 | DQ124956 | DQ125014 | - | - |
| 370 | Guatteria_modesta_1 | *Guatteria modesta* Diels | - | DQ125081 | - | DQ125179 | DQ124957 | DQ125015 | - | - |
| 371 | Guatteria_decurrens _1 | *Guatteria multivenia* Diels = *Guatteria decurrens* R. E. Fr. | - | AY740938 | - | DQ125180 | AY740987 | AY741036 | - | - |
| 372 | Guatteria_australis_4 | *Guatteria neglecta* (Griseb.) P. Wilson ex León & Alain = *Guatteria australis* A.St.-Hil. | - | DQ861713 | - | DQ861759 | DQ861810 | DQ861862 | - | - |
| 373 | Guatteria_notabilis | *Guatteria notabilis* Mello-Silva & Pirani | - | DQ125082 | - | DQ125181 | DQ124958 | DQ125016 | - | - |
| 374 | Guatteria_oligocarpa | *Guatteria oligocarpa* Mart. | - | AY740939 | - | DQ125182 | AY740988 | AY741037 | - | - |
| 375 | Guatteria_punctata_5 | *Guatteria olivacea* R. E. Fr. = *Guatteria punctata* (Aubl.) R. A. Howard | - | AY740940 | - | DQ125183 | AY740989 | AY741038 | - | - |
| 376 | Guatteria_oliviformis | *Guatteria oliviformis* Donn. Sm. | - | AY740941 | - | DQ125184 | AY740990 | AY741039 | - | - |
| 377 | Guatteria_ouregou | *Guatteria ouregou* (Aubl.) Dunal | - | AY740942 | - | DQ125185 | AY740991 | AY741040 | - | - |

| No. | Voucher information | | GenBank accession numbers | | | | | | | |
|---|---|---|---|---|---|---|---|---|---|---|
| | Tip name in the tree | Taxa name | *atpB-rbcL* | *matK* | *ndhF* | *psbA-trnH* | *rbcL* | *trnL-F* | *trnS-G* | *ycf1* |
| 378 | Guatteria_punctata_4 | *Guatteria ovalifolia* R. E. Fr. = *Guatteria punctata* (Aubl.) R. A. Howard | - | DQ861715 | - | DQ861761 | DQ861812 | DQ861864 | - | - |
| 379 | Guatteria_pacifica | *Guatteria pacifica* R. E. Fr. | - | DQ125083 | - | DQ125186 | DQ124960 | DQ125018 | - | - |
| 380 | Guatteria_citriodora_2 | *Guatteria paraensis* R. E. Fr. = *Guatteria citriodora* Ducke | - | DQ125085 | - | DQ125188 | DQ124961 | DQ125019 | - | - |
| 381 | Guatteria_australis_2 | *Guatteria parvifolia* R. E. Fr. = *Guatteria australis* A.St.-Hil. | - | AY740943 | - | DQ861762 | AY740992 | AY741041 | - | - |
| 382 | Guatteria_pittieri_1 | *Guatteria pittieri* R. E. Fr. | - | AY740944 | - | DQ125190 | AY740993 | AY741042 | - | - |
| 383 | Guatteria_punctata_3 | *Guatteria poeppigiana* Mart. = *Guatteria punctata* (Aubl.) R. A. Howard | - | DQ125086 | - | DQ125191 | DQ124962 | DQ125020 | - | - |
| 384 | Guatteria_pogonopus_1 | *Guatteria pogonopus* Mart. | - | DQ125087 | - | DQ125192 | DQ124963 | DQ125021 | - | - |
| 385 | Guatteria_pohliana | *Guatteria pohliana* Schltdl. | - | DQ125088 | - | DQ125193 | DQ124964 | DQ125022 | - | - |
| 386 | Guatteria_polyantha | *Guatteria polyantha* R. E. Fr. | - | DQ125089 | - | DQ125194 | DQ124965 | DQ125023 | - | - |
| 387 | Guatteria_australis_8 | *Guatteria polycarpa* R. E. Fr. = *Guatteria australis* A.St.-Hil. | - | DQ125090 | - | DQ125195 | DQ124966 | DQ125024 | - | - |
| 388 | Guatteria_sellowiana_3 | *Guatteria pubens* (Mart.) R. E. Fr. | - | DQ125091 | - | DQ125196 | DQ124967 | DQ125025 | - | - |
| 389 | Guatteria_pudica | *Guatteria pudica* N. Zamora & Maas | JQ513884 | AY740945 | JQ769093 | DQ125197 | AY740994 | AY741043 | - | - |
| 390 | Guatteria_punctata_1 | *Guatteria punctata* (Aubl.) R. A. Howard | - | AY740946 | - | DQ125198 | AY740995 | AY741044 | - | - |
| 391 | Guatteria_modesta_2 | *Guatteria puncticulata* R. E. Fr. = *Guatteria modesta* Diels | - | AY740947 | - | DQ125199 | AY740996 | AY741045 | - | - |

| No. | Voucher information | | GenBank accession numbers | | | | | | | |
|---|---|---|---|---|---|---|---|---|---|---|
| | Tip name in the tree | Taxa name | *atpB-rbcL* | *matK* | *ndhF* | *psbA-trnH* | *rbcL* | *trnL-F* | *trnS-G* | *ycf1* |
| 392 | Guatteria_ramiflora | *Guatteria ramiflora* (D. R. Simpson) Erkens & Maas | - | DQ125064 | - | DQ125135 | DQ124940 | DQ124998 | - | - |
| 393 | Guatteria_ucayalina_4 | *Guatteria recurvisepala* R. E. Fr. = *Guatteria ucayalina* Huber | - | AY740948 | - | DQ125200 | AY740997 | AY741046 | - | - |
| 394 | Guatteria_australis_7 | *Guatteria reflexa* R. E. Fr. = *Guatteria australis* A.St.-Hil. | - | DQ125092 | - | DQ125201 | DQ124968 | DQ125026 | - | - |
| 395 | Guatteria_rigida | *Guatteria rigida* R. E. Fr. | - | DQ861718 | - | - | DQ861814 | DQ861867 | - | - |
| 396 | Guatteria_dolichopoda_3 | *Guatteria rigidipes* R. E. Fr. = *Guatteria dolichopoda* Donn. Sm. | - | DQ125094 | - | DQ125203 | DQ124970 | DQ125028 | - | - |
| 397 | Guatteria_inundata_2 | *Guatteria riparia* R. E. Fr. = *Guatteria inundata* Mart. | - | DQ861719 | - | DQ861764 | DQ861815 | DQ861868 | - | - |
| 398 | Guatteria_rotundata | *Guatteria rotundata* Maas & van Setten | - | AY740949 | - | DQ125204 | AY740998 | AY741047 | - | - |
| 399 | Guatteria_rupestris | *Guatteria rupestris* Mello-Silva & Pirani | - | AY740950 | - | DQ125205 | AY740999 | AY741048 | - | - |
| 400 | Guatteria_sabuletorum | *Guatteria sabuletorum* R. E. Fr. | - | DQ861721 | - | DQ861766 | DQ861817 | DQ861870 | - | - |
| 401 | Guatteria_saffordiana | *Guatteria saffordiana* Pittier | - | DQ861720 | - | DQ861765 | DQ861816 | DQ861869 | - | - |
| 402 | Guatteria_salicifolia | *Guatteria salicifolia* R. E. Fr. | - | AY740951 | - | - | AY741000 | AY741049 | - | - |
| 403 | Guatteria_scandens | *Guatteria scandens* Ducke | - | DQ125095 | - | DQ125207 | DQ124971 | DQ125029 | - | - |
| 404 | Guatteria_pogonopus_2 | *Guatteria schlechtendaliana* Mart. = *Guatteria pogonopus* Mart. | - | DQ125096 | - | DQ125208 | DQ124972 | DQ125030 | - | - |
| 405 | Guatteria_schomburgkiana_1 | *Guatteria schomburgkiana* Mart. | - | AY740952 | - | DQ125209 | AY741001 | DQ861871 | - | - |

| No. | Voucher information | | GenBank accession numbers | | | | | | | |
|---|---|---|---|---|---|---|---|---|---|---|
| | Tip name in the tree | Taxa name | *atpB-rbcL* | *matK* | *ndhF* | *psbA-trnH* | *rbcL* | *trnL-F* | *trnS-G* | *ycf1* |
| 406 | Guatteria_ucayalina_3 | *Guatteria schunkevigoi* D. R. Simpson = *Guatteria ucayalina* Huber | - | DQ125097 | - | DQ125210 | DQ124973 | DQ125031 | - | - |
| 407 | Guatteria_scytophylla_1 | *Guatteria scytophylla* Diels | - | AY740953 | - | DQ125211 | AY741002 | AY741051 | - | - |
| 408 | Guatteria_sellowiana_1 | *Guatteria sellowiana* Schltdl. | - | AY740954 | - | DQ125212 | AY741003 | AY841702 | - | - |
| 409 | Guatteria_sessilicarpa | *Guatteria sessilicarpa* Maas & van Setten | - | AY740955 | - | DQ125214 | DQ124974 | DQ125032 | - | - |
| 410 | Guatteria_schomburgkiana_2 | *Guatteria sessilis* R. E. Fr. = *Guatteria schomburgkiana* Mart. | - | DQ125099 | - | DQ125215 | DQ124975 | DQ125033 | - | - |
| 411 | Guatteria_australis_6 | *Guatteria sordida* var. *ovalis* R. E. Fr. = *Guatteria australis* A.St.-Hil. | - | DQ125100 | - | DQ125216 | DQ124976 | DQ125034 | - | - |
| 412 | Guatteria_pittieri_2 | *Guatteria sphaerantha* R. E. Fr. = *Guatteria pittieri* R. E. Fr. | - | DQ125101 | - | DQ125217 | DQ124977 | DQ125035 | - | - |
| 413 | Guatteria_ucayalina_5 | *Guatteria stipitata* R. E. Fr. = *Guatteria ucayalina* Huber. | - | DQ125102 | - | DQ125218 | DQ124978 | DQ125036 | - | - |
| 414 | Guatteria_subsessilis | *Guatteria subsessilis* Mart. | - | DQ125103 | - | DQ125219 | DQ124979 | DQ125037 | - | - |
| 415 | Guatteria_talamancana | *Guatteria talamancana* N. Zamora & Maas | - | DQ861729 | - | DQ861773 | DQ861825 | DQ861878 | - | - |
| 416 | Guatteria_tomentosa_1 | *Guatteria tomentosa* Rusby 9521 | - | DQ861730 | - | DQ861774 | DQ861826 | DQ861879 | - | - |
| 417 | Guatteria_dolichopoda_2 | *Guatteria tonduzii* Diels = *Guatteria dolichopoda* Donn. Sm. | - | AY740956 | - | DQ125228 | AY741005 | AY741054 | - | - |
| 418 | Guatteria_tomentosa_2 | *Guatteria trichoclonia* Diels = *Guatteria tomentosa* Rusby | - | DQ125112 | - | DQ125229 | DQ861827 | DQ125046 | - | - |
| 419 | Guatteria_ucayaliana_1 | *Guatteria ucayaliana* Diels | - | DQ861732 | - | DQ861776 | DQ861828 | DQ861881 | - | - |

| No. | Voucher information | | GenBank accession numbers | | | | | | | |
|---|---|---|---|---|---|---|---|---|---|---|
| | Tip name in the tree | Taxa name | *atpB-rbcL* | *matK* | *ndhF* | *psbA-trnH* | *rbcL* | *trnL-F* | *trnS-G* | *ycf1* |
| 420 | Guatteria_venezuelana | *Guatteria venezuelana* R. E. Fr. | - | DQ125113 | - | DQ125230 | DQ861831 | DQ125047 | - | - |
| 421 | Guatteria_verruculosa | *Guatteria verruculosa* R. E. Fr. | - | DQ125114 | - | DQ125231 | DQ124990 | DQ125048 | - | - |
| 422 | Guatteria_villosissima | *Guatteria villosissima* A. St.-Hil. | - | AY740957 | - | DQ125232 | AY741006 | AY741055 | - | - |
| 423 | Guatteria_wachenheimii | *Guatteria wachenheimii* Benoist | - | DQ125115 | - | DQ125233 | DQ124991 | DQ125049 | - | - |
| 424 | Guatteria_zamorae | *Guatteria zamorae* Erkens & Maas | - | DQ861736 | - | DQ861780 | DQ861832 | DQ861885 | - | - |
| 425 | Guatteria_trichocarpa | *Guatteriella tomentosa* R.E.Fr. 4494 = *Guatteria trichocarpa* Erkens & Maas | - | DQ861737 | - | DQ861781 | DQ861833 | DQ861886 | - | - |
| 426 | Hexalobus_crispiflorus | *Hexalobus crispiflorus* A. Rich. | - | - | EU169713 | EU169737 | EU169760 | EU169782 | EU169804 | - |
| 427 | Hexalobus_salicifolius | *Hexalobus salicifolius* Engl. | - | - | EU169714 | EU169738 | EU169761 | EU169783 | EU169805 | - |
| 428 | Hornschuchia_citriodora | *Hornschuchia citriodora* D. M. Johnson | - | - | - | - | AY841625 | AY841703 | - | - |
| 429 | Huberantha_cerasoides | *Huberantha cerasoides* (Roxb.) Chaowasku | - | AY518854 | JQ889985 | JQ889980 | - | AY319131 | - | JQ723950 |
| 430 | Huberantha_decora | *Huberantha decora* (Diels) Chaowasku | - | - | JX544879 | JX544859 | - | JX544869 | - | JX544849 |
| 431 | Huberantha_henrici | *Huberantha henrici* (Diels) Chaowasku | - | - | JX544880 | JX544860 | - | JX544870 | - | JX544850 |
| 432 | Huberantha_jenkinsii | *Huberantha jenkinsii* (Hook. f. & Thomson) Chaowasku | - | - | JX544842 | JX544812 | - | JX544803 | - | JX544821 |
| 433 | Huberantha_korinti | *Huberantha korinti* (Dunal) Chaowasku | EU522345 | EU522234 | JX544877 | EU522124 | EU522289 | EU522179 | - | JX544847 |
| 434 | Huberantha_nitidissima | *Huberantha nitidissima* (Dunal) Chaowasku | - | JQ889989 | JQ889986 | JQ889981 | - | JQ889988 | - | JQ889976 |

| No. | Voucher information | | GenBank accession numbers | | | | | | | |
|---|---|---|---|---|---|---|---|---|---|---|
| | Tip name in the tree | Taxa name | *atpB-rbcL* | *matK* | *ndhF* | *psbA-trnH* | *rbcL* | *trnL-F* | *trnS-G* | *ycf1* |
| 435 | Huberantha_pendula | *Huberantha pendula* (Capuron ex G. E. Schatz & Le Thomas) Chaowasku | - | AY518852 | JQ889987 | JQ889982 | - | AY319144 | - | JQ889977 |
| 436 | Huberantha_perrieri | *Huberantha perrieri* (Cavaco & Keraudren) Chaowasku | - | - | JX544881 | JX544861 | - | JX544871 | - | JX544851 |
| 437 | Huberantha_rumphii | *Huberantha rumphii* (Blume ex Hensch.) Chaowasku | - | AY518791 | JX544841 | JX544811 | - | AY319145 | - | JX544820 |
| 438 | Huberantha_stuhlmannii | *Huberantha stuhlmannii* (Engl.) Chaowasku | - | AY518853 | JX544882 | JX544862 | - | AY319149 | - | JX544852 |
| 439 | Huberantha_tanganyikens is | *Huberantha tanganyikensis* (Vollesen) Chaowasku | - | - | JX544883 | JX544863 | - | JX544872 | - | JX544853 |
| 440 | Isolona_campanulata | *Isolona campanulata* Engl. & Diels | EF179266 | AY238963 | EF179301 | DQ125127 | AY238954 | EF179318 | EF179343 | - |
| 441 | Isolona_capuronii | *Isolona capuronii* Cavaco & Keraudren | - | - | - | EU216662 | - | EU216708 | EU216617 | - |
| 442 | Isolona_cauliflora | *Isolona cauliflora* Verdc. | - | - | EU169716 | EU169739 | EU169762 | EU169784 | EU169807 | - |
| 443 | Isolona_congolana | *Isolona congolana* (De Wild. & T. Durand) Engl. & Diels 3852 | - | - | EU216637 | EU216658 | - | EU216704 | EU216613 | - |
| 444 | Isolona_cooperi | *Isolona congolana* (De Wild. & T. Durand) Engl. & Diels 9550 | - | - | EU216644 | EU216668 | - | EU216714 | EU216623 | - |
| 445 | Isolona_dewevrei | *Isolona cooperi* Hutch. & Dalziel ex G. P. Cooper & Record | - | - | EU216636 | EU216657 | AY841626 | AY841704 | EU216612 | - |
| 446 | Isolona_ghesquierei | *Isolona dewevrei* (De Wild. & T. Durand) Engl. & Diels | - | - | EU216645 | EU216669 | - | EU216715 | EU216624 | - |
| 447 | Isolona_heinsenii | *Isolona ghesquierei* Cavaco & Keraudren | - | - | - | EU216663 | - | EU216709 | EU216618 | - |

| No. | Voucher information | | GenBank accession numbers | | | | | | | |
|---|---|---|---|---|---|---|---|---|---|---|
| | Tip name in the tree | Taxa name | *atpB-rbcL* | *matK* | *ndhF* | *psbA-trnH* | *rbcL* | *trnL-F* | *trnS-G* | *ycf1* |
| 448 | Isolona_hexaloba | *Isolona heinsenii* Engl. & Diels | - | - | EU216640 | EU216664 | - | EU216710 | EU216619 | - |
| 449 | Isolona_linearis | *Isolona hexaloba* Pierre ex Engl. & Diels | - | - | EU169717 | EU169740 | EU169763 | EU169785 | EU169808 | - |
| 450 | Isolona_maitlandii | *Isolona linearis* Couvreur | - | - | EU216641 | EU216665 | - | EU216711 | EU216620 | - |
| 451 | Isolona_perrieri | *Isolona perrieri* Diels | - | - | EU216639 | EU216661 | - | EU216707 | EU216616 | - |
| 452 | Isolona_pleurocarpa | *Isolona pleurocarpa* Diels | - | - | EU216642 | EU216666 | - | EU216712 | EU216621 | - |
| 453 | Isolona_thonneri | *Isolona thonneri* (De Wild. & T. Durand) Engl. & Diels | - | - | EU216643 | EU216667 | - | EU216713 | EU216622 | - |
| 454 | Isolona_zenkeri | *Isolona zenkeri* Engl. | - | - | EU216638 | EU216660 | - | EU216705 | EU216614 | - |
| 455 | Klarobelia_candida | *Klarobelia candida* Chatrou | - | - | - | AY841467 | AY841525 | AY841539 | - | - |
| 456 | Klarobelia_cauliflora | *Klarobelia cauliflora* Chatrou | - | - | - | AY841468 | AY841627 | AY841705 | - | - |
| 457 | Klarobelia_inundata | *Klarobelia inundata* Chatrou | AY841378 | AY743490 | AY841409 | AY841469 | AY743452 | AY743471 | AY841556 | - |
| 458 | Klarobelia_megalocarpa | *Klarobelia megalocarpa* Chatrou | - | AY518866 | - | AY841470 | - | AY319176 | - | - |
| 459 | Klarobelia_sp | *Klarobelia* sp. | - | - | - | AY841471 | AY841526 | AY841540 | - | - |
| 460 | Klarobelia_stipitata | *Klarobelia stipitata* Chatrou | - | - | - | AY841472 | AY841628 | AY841706 | - | - |
| 461 | Letestudoxa_bella | *Letestudoxa bella* Pellegr. | EF179267 | DQ125059 | EF179302 | DQ125128 | AY841629 | AY841707 | EF179344 | - |
| 462 | Letestudoxa_glabrifolia | *Letestudoxa glabrifolia* Chatrou & Repetur | - | - | - | - | AY841630 | AY841708 | - | - |
| 463 | Lettowianthus_stellatus | *Lettowianthus stellatus* Diels | - | - | - | EU169730 | EU169753 | EU169775 | EU169797 | - |
| 464 | Liriodendron_chinense | *Liriodendron chinense* (Hemsl.) Sarg. | - | - | - | AY841424 | AY841593 | AY841670 | - | - |
| 465 | Maasia_discolor | *Maasia discolor* (Diels) Mols, Keßler & Rogstad | AY841385 | AY518872 | AY841416 | AY841500 | - | AY319135 | AY841563 | - |

| No. | Voucher information | | GenBank accession numbers | | | | | | | |
|---|---|---|---|---|---|---|---|---|---|---|
| | Tip name in the tree | Taxa name | *atpB-rbcL* | *matK* | *ndhF* | *psbA-trnH* | *rbcL* | *trnL-F* | *trnS-G* | *ycf1* |
| 466 | Maasia_glauca | *Maasia glauca* (Hassk.) Mols, Keßler & Rogstad | - | AY518871 | - | AY841501 | GQ248676 | AY319137 | - | - |
| 467 | Maasia_multinervis | *Maasia multinervis* (Diels) Mols, Keßler & Rogstad | - | JF810383 | - | - | JF810395 | JF810407 | - | - |
| 468 | Maasia_ovalifolia | *Maasia ovalifolia* (Rogstad) Mols, Keßler & Rogstad | - | JF810384 | - | - | JF810396 | JF810408 | - | - |
| 469 | Maasia_sumatrana | *Maasia sumatrana* (Miq.) Mols, Keßler & Rogstad | AY841387 | AY518873 | AY841418 | AY841503 | - | AY319153 | AY841565 | - |
| 470 | Magnolia_kobus | *Magnolia kobus* DC. | - | AY743476 | - | AY841425 | AY743438 | AY743457 | - | - |
| 471 | Malmea_dielsiana | *Malmea dielsiana* R. E. Fr. | AY841379 | AY238964 | AY841410 | AY841473 | AY238955 | AY319177 | AY841557 | - |
| 472 | Malmea_dimera | *Malmea dimera* Chatrou | - | - | - | AY841474 | AY841631 | AY841709 | - | - |
| 473 | Malmea_sp_Chatrou_8 | *Malmea* sp. Chatrou et al 8 | AY841380 | AY841397 | AY841411 | AY841475 | AY841527 | AY841541 | AY841558 | - |
| 474 | Malmea_surinamensis | *Malmea surinamensis* Chatrou | - | AY743491 | - | AY841476 | AY743453 | AY743472 | - | - |
| 475 | Marsypopetalum_crassum | *Marsypopetalum crassum* (R. Parker) B. Xue & R. M. K. Saunders | - | HQ286571 | JQ723792 | - | HQ286577 | HQ286583 | - | JQ723929 |
| 476 | Marsypopetalum_littorale | *Marsypopetalum littorale* (Blume) B. Xue & R. M. K. Saunders | - | AY518835 | JX544827 | JX544804 | - | AY319140 | - | JX544813 |
| 477 | Marsypopetalum_lucidum | *Marsypopetalum lucidum* (Merr.) B. Xue & R. M. K. Saunders | - | HQ286572 | - | - | HQ286578 | HQ286584 | - | - |
| 478 | Marsypopetalum_triste | *Marsypopetalum triste* (Pierre) B. Xue & R. M. K. Saunders | - | HQ286573 | - | - | HQ286579 | HQ286585 | - | - |
| 479 | Meiocarpidium_lepidotum | *Meiocarpidium lepidotum* (Oliv.) Engl. & Diels | - | - | - | EU169731 | EU169754 | EU169776 | EU169798 | - |

| No. | Voucher information | | GenBank accession numbers | | | | | | | |
|---|---|---|---|---|---|---|---|---|---|---|
| | Tip name in the tree | Taxa name | *atpB-rbcL* | *matK* | *ndhF* | *psbA-trnH* | *rbcL* | *trnL-F* | *trnS-G* | *ycf1* |
| 480 | Meiogyne_amicorum | *Meiogyne amicorum* (A. C. Sm.) B. Xue, D. M. Johnson & R. M. K. Saunders [= *Polyalthia amicorm A. C. Sm.*] | - | KF301021 | KF611912 | | - | KF573503 | | - |
| 481 | Meiogyne_amygdalina | *Meiogyne amygdalina* (A. Gray) B. Xue, D. M. Johnson & R. M. K. Saunders | - | KF301022 | KF611913 | | - | KF573497 | | - |
| 482 | Meiogyne_baillonii | *Meiogyne baillonii* (Guillaumin) Heusden | - | JQ723768 | JQ723793 | - | JQ723855 | JQ723908 | - | JQ723930 |
| 483 | Meiogyne_bidwillii | *Meiogyne bidwillii* (Benth.) D. C. Thomas, Chaowasku & R. M. K. Saunders | - | AY518795 | JQ723789 | - | JQ723851 | AY319089 | - | JQ723925 |
| 484 | Meiogyne_cylindrocarpa_subsp_cylindrocarpa_1 | *Meiogyne cylindrocarpa* (Burck) Heusden subsp. *cylindrocarpa* 1 | - | AY518796 | JQ723794 | | AY318981 | AY319093 | | JQ723931 |
| 485 | Meiogyne_cylindrocarpa_subsp_cylindrocarpa_2 | *Meiogyne cylindrocarpa* (Burck) Heusden subsp. *cylindrocarpa* 2 | - | JQ723769 | JQ723795 | | JQ723856 | JQ723909 | | JQ723932 |
| 486 | Meiogyne_dumetosa | *Meiogyne dumetosa*(Vieill. ex Guillaumin) Heusden | - | KF301024 | KF611915 | | KF301030 | KF573501 | | - |
| 487 | Meiogyne_glabra | *Meiogyne glabra* Heusden | - | JQ723772 | - | - | JQ723859 | JQ723912 | - | JQ723935 |
| 488 | Meiogyne_habrotricha | *Meiogyne habrotricha* (A. C. Sm.) B. Xue, D. M. Johnson & R. M. K. Saunders | - | KF301025 | KF611916 | | - | KF573498 | | - |
| 489 | Meiogyne_hainanensis | *Meiogyne hainanensis* (Merr.) Bân | - | JQ723773 | - | - | JQ723860 | - | - | JQ723936 |

| No. | Voucher information | | GenBank accession numbers | | | | | | | |
|---|---|---|---|---|---|---|---|---|---|---|
| | Tip name in the tree | Taxa name | *atpB-rbcL* | *matK* | *ndhF* | *psbA-trnH* | *rbcL* | *trnL-F* | *trnS-G* | *ycf1* |
| 490 | Meiogyne_heteropetala | *Meiogyne heteropetala* (F. Muell.) D. C. Thomas, Chaowasku & R. M. K. Saunders | - | JQ723766 | JQ723790 | - | JQ723853 | JQ723906 | - | JQ723927 |
| 491 | Meiogyne_hirsuta | *Meiogyne hirsuta* (Jessup) Jessup | - | JQ723774 | JQ723798 | - | JQ723861 | JQ723914 | - | JQ723937 |
| 492 | Meiogyne_insularis | *Meiogyne insularis* (A. C. Sm.) D. C. Thomas, B. Xue & R. M. K. Saunders [= *Meiogyne stenopetala* subsp. *insularis* (A. C. Sm.) Heusden] | - | KF301028 | KF611919 | | - | KF573502 | | - |
| 493 | Meiogyne_laddiana | *Meiogyne laddiana* (A. C. Sm.) B. Xue, D. M. Johnson & R. M. K. Saunders | - | KF301026 | KF611917 | | - | KF573499 | | - |
| 494 | Meiogyne_lecardii | *Meiogyne lecardii* (Guillaumin) Heusden | - | JQ723775 | JQ723799 | - | JQ723862 | JQ723915 | - | JQ723938 |
| 495 | Meiogyne_mindorensis | *Meiogyne mindorensis* (Merr.) Heusden | - | JQ723776 | JQ723800 | - | JQ723863 | JQ723916 | - | JQ723939 |
| 496 | Meiogyne_monosperma | *Meiogyne monosperma* (Hook. f. & Thomson) Heusden | - | JQ723777 | - | - | JQ723864 | - | - | JQ723940 |
| 497 | Meiogyne_pannosa | *Meiogyne pannosa* (Dalzell) J. Sinclair | - | JQ723778 | JQ723801 | - | JQ723865 | JQ723918 | - | JQ723941 |
| 498 | Meiogyne_sp2_Indu2457 | *Meiogyne* sp. | - | AY518797 | JQ723802 | | AY319002 | AY319115 | | JQ723942 |
| 499 | Meiogyne_stenopetala | *Meiogyne stenopetala* (F. Muell.) Heusden | - | JQ723779 | JQ723803 | - | JQ723866 | AY319083 | - | JQ723943 |
| 500 | Meiogyne_trichocarpa_1 | *Meiogyne trichocarpa* (Jessup) D. C. Thomas & R. M. K. Saunders 1 | - | JQ723770 | JQ723796 | | JQ723857 | JQ723910 | | JQ723933 |

| No. | Voucher information | | GenBank accession numbers | | | | | | | |
|---|---|---|---|---|---|---|---|---|---|---|
| | Tip name in the tree | Taxa name | *atpB-rbcL* | *matK* | *ndhF* | *psbA-trnH* | *rbcL* | *trnL-F* | *trnS-G* | *ycf1* |
| 501 | Meiogyne_trichocarpa_2 | *Meiogyne trichocarpa* (Jessup) D. C. Thomas & R. M. K. Saunders 2 [= *Meiogyne cylindrocarpa* (Burck) Heusden subsp. *trichocarpa* Jessup 2 | - | JQ723771 | JQ723797 | | JQ723858 | JQ723911 | | JQ723934 |
| 502 | Meiogyne_verrucosa | *Meiogyne verrucosa* Jessup | - | JQ723780 | JQ723804 | - | JQ723867 | JQ723920 | - | JQ723944 |
| 503 | Meiogyne_virgata | *Meiogyne virgata* (Blume) Miq. | - | AY518798 | JX544769 | JX544784 | - | AY319094 | - | JQ723945 |
| 504 | Meiogyne_cylindrocarpa_Guam | *Meiogyne cylindrocarpa*(Burck) Heusden | - | KF301023 | KF611914 | | KF301029 | KF573500 | | - |
| 505 | Meiogyne_sp_TNW_146 | *Meiogyne*sp, TNW 146 | - | KF301027 | KF611918 | | KF301031 | KF573504 | | KF301032 |
| 506 | Mezzettia_parviflora | *Mezzettia parviflora* Becc. | - | AY518881 | - | - | - | AY319095 | - | - |
| 507 | Miliusa_amplexicaulis | *Miliusa amplexicaulis* Ridl. | - | - | JQ690479 | JQ690480 | - | JQ690478 | - | JQ690481 |
| 508 | Miliusa_brahei | *Miliusa brahei* (F. Muell.) Jessup | - | - | JQ690431 | JQ690432 | - | JQ690430 | - | JQ690433 |
| 509 | Miliusa_campanulata | *Miliusa campanulata* Pierre | - | AY518842 | JQ690487 | JQ690488 | - | AY319096 | - | JQ690489 |
| 510 | Miliusa_cuneata | *Miliusa cuneata* Craib | - | AY518844 | JQ690491 | JQ690492 | - | AY319097 | - | JQ690493 |
| 511 | Miliusa_fusca | *Miliusa fusca* Pierre | - | - | JQ690443 | JQ690444 | - | JQ690442 | - | JQ690445 |
| 512 | Miliusa_horsfieldii | *Miliusa horsfieldii* (Benn.) Pierre | - | AY518849 | JQ690447 | JQ690448 | - | AY319098 | - | JQ690449 |
| 513 | Miliusa_indica | *Miliusa indica* Lesch. ex A. DC. | - | JQ723781 | JQ723806 | - | JQ723868 | JQ723921 | - | JQ723946 |
| 514 | Miliusa_koolsii | *Miliusa koolsii* (Kosterm.) J. Sinclair | - | - | JQ690455 | JQ690456 | - | JQ690454 | - | JQ690457 |
| 515 | Miliusa_lanceolata | *Miliusa lanceolata* Chaowasku & Ke ßler | - | - | JQ690459 | JQ690460 | - | JQ690458 | - | JQ690461 |
| 516 | Miliusa_macrocarpa | *Miliusa macrocarpa* Hook. f. & Thomson | - | JQ690499 | JQ690500 | JQ690501 | - | JQ690498 | - | JQ690502 |
| 517 | Miliusa_macropoda | *Miliusa macropoda* Miq. | - | - | JQ690463 | JQ690464 | - | JQ690462 | - | JQ690465 |

| No. | Voucher information | | GenBank accession numbers | | | | | | | |
|---|---|---|---|---|---|---|---|---|---|---|
| | Tip name in the tree | Taxa name | *atpB-rbcL* | *matK* | *ndhF* | *psbA-trnH* | *rbcL* | *trnL-F* | *trnS-G* | *ycf1* |
| 518 | Miliusa_montana | *Miliusa montana* Gardner ex Hook. f. & Thomson | - | JQ690507 | JQ690508 | JQ690509 | - | JQ690506 | - | JQ690510 |
| 519 | Miliusa_novoguineensis | *Miliusa novoguineensis* Mols & Keßler | - | - | JQ690467 | JQ690468 | - | JQ690466 | - | JQ690469 |
| 520 | Miliusa_parviflora | *Miliusa parviflora* Ridl. | - | - | JQ690471 | JQ690472 | - | JQ690470 | - | JQ690473 |
| 521 | Miliusa_sclerocarpa | *Miliusa sclerocarpa* (A. DC.) Kurz | - | - | JQ690475 | JQ690476 | - | JQ690474 | - | JQ690477 |
| 522 | Miliusa_thorelii | *Miliusa thorelii* Finet & Gagnep. | - | AY518846 | JQ690519 | JQ690520 | - | AY319104 | - | JQ690521 |
| 523 | Miliusa_traceyi | *Miliusa traceyi* Jessup | - | JQ690532 | JQ690533 | JQ690534 | - | JQ690531 | - | JQ690535 |
| 524 | Miliusa_velutina | *Miliusa velutina* (Dunal) Hook. f. & Thomson | - | AY518847 | JQ690536 | JQ690537 | - | AY319105 | - | JQ690538 |
| 525 | Mischogyne_michelioides | *Mischogyne michelioides* Exell | - | - | EU169718 | EU169741 | EU169764 | EU169786 | EU169809 | - |
| 526 | Mitrella_kentii | *Mitrella kentii* (Blume) Miq. | - | FJ743751 | JQ768616 | FJ743789 | AY841633 | AY841711 | - | - |
| 527 | Mitrephora_alba | *Mitrephora alba* Ridl. | - | AY518855 | JQ889983 | JQ889978 | - | AY319106 | - | JQ723947 |
| 528 | Mitrephora_celebica | *Mitrephora celebica* Scheff. | - | AY518859 | - | - | - | AY319107 | - | - |
| 529 | Mitrephora_keithii | *Mitrephora keithii* Ridl. | EU522343 | AY518857 | - | EU522122 | - | AY319108 | - | - |
| 530 | Mitrephora_maingayi | *Mitrephora maingayi* Hook. f. & Thomson | - | AY518856 | - | - | - | AY319109 | - | - |
| 531 | Mitrephora_polypyrena | *Mitrephora polypyrena* (Blume) Miq. | - | AY518858 | - | - | - | AY319110 | - | - |
| 532 | Mitrephora_vittata | *Mitrephora vittata* Weeras. & R. M. K. Saunders | - | FJ463223 | - | - | - | FJ463224 | - | - |
| 533 | Mkilua_fragrans | *Mkilua fragrans* Verdc. | EF179268 | - | EF179303 | DQ861696 | AY841634 | AY841712 | EF179345 | - |
| 534 | Monanthotaxis_ambrensis | *Monanthotaxis ambrensis* (Cavaco & Keraudren) Verdc. | - | KX761295 | KX787015 | KX786954 | KX761326 | KX786984 | | |

| No. | Voucher information | | GenBank accession numbers | | | | | | | |
|---|---|---|---|---|---|---|---|---|---|---|
| | Tip name in the tree | Taxa name | *atpB-rbcL* | *matK* | *ndhF* | *psbA-trnH* | *rbcL* | *trnL-F* | *trnS-G* | *ycf1* |
| 535 | Monanthotaxis_angustifolia | *Monanthotaxis angustifolia* (Exell) Verdc. | - | KX761303 | KX787023 | KX786962 | KX761334 | KX786992 | | |
| 536 | Monanthotaxis_aquila | *Monanthotaxis aquila* P.H.Hoekstra | - | KX761309 | KX787029 | KX786968 | KX761340 | KX786998 | | |
| 537 | Monanthotaxis_atewaesis | *Monanthotaxis atewensis* P.H.Hoekstra | - | KX761307 | KX787027 | KX786966 | KX761338 | KX786996 | | |
| 538 | Monanthotaxis_atopostema | *Monanthotaxis atopostema* P.H.Hoekstra | - | KX761310 | KX787030 | KX786969 | KX761341 | KX786999 | | |
| 539 | Monanthotaxis_boivinii | *Monanthotaxis boivinii* (Baill.) Verdc. | - | KX761299 | KX787019 | KX786958 | KX761330 | KX786988 | | |
| 540 | Monanthotaxis_buchananii | *Monanthotaxis buchananii* (Engl.) Verdc. | - | JQ768581 | JQ768617 | JQ768660 | JQ768700 | JQ768742 | | |
| 541 | Monanthotaxis_congoensis | *Monanthotaxis congoensis* Baill. | - | KX761287 | KX787007 | KX786946 | KX761318 | KX786976 | | |
| 542 | Monanthotaxis_couvreurii | *Monanthotaxis couvreurii* P.H.Hoekstra | - | KX761311 | KX787031 | KX786970 | KX761342 | KX787000 | | |
| 543 | Monanthotaxis_enghiana | *Monanthotaxis enghiana* (Diels) P.H.Hoekstra | - | JQ768578 | JQ768613 | JQ768657 | JQ768697 | JQ768739 | | - |
| 544 | Monanthotaxis_ferruginea | *Monanthotaxis ferruginea* (Oliv.) Verdc. | - | KX761306 | KX787026 | KX786965 | KX761337 | KX786995 | | - |
| 545 | Monanthotaxis_filamentosa | *Monanthotaxis filamentosa* (Diels) Verdc. | - | KX761291 | KX787011 | KX786950 | KX761322 | KX786980 | | |
| 546 | Monanthotaxis_filipes | *Monanthotaxis filipes* P.H.Hoekstra | - | KX761315 | KX787035 | KX786973 | KX761346 | KX787004 | | |
| 547 | Monanthotaxis_foliosa_var_ferruginea | *Monanthotaxis foliosa* var. *ferruginea* (Robyns & Ghesq.) Verdc. | - | KX761290 | KX787010 | KX786949 | KX761321 | KX786979 | | |

| No. | Voucher information | | GenBank accession numbers | | | | | | | |
|---|---|---|---|---|---|---|---|---|---|---|
| | Tip name in the tree | Taxa name | *atpB-rbcL* | *matK* | *ndhF* | *psbA-trnH* | *rbcL* | *trnL-F* | *trnS-G* | *ycf1* |
| 548 | Monanthotaxis_fornicata | *Monanthotaxis fornicata* (Baill.) Verdc. | - | JQ768583 | JQ768619 | JQ768662 | JQ768702 | JQ768744 | | |
| 549 | Monanthotaxis_glaucifoli a | *Monanthotaxis glaucifolia* (Hutch. & Dalziel) P.H. Koekstra | - | KX761298 | KX787018 | KX786957 | KX761329 | KX786987 | | |
| 550 | Monanthotaxis_glomerula ta | *Monanthotaxis glomerulata* (Le Thomas) Verdc. | - | JQ768584 | JQ768620 | JQ768663 | JQ768703 | JQ768745 | - | - |
| 551 | Monanthotaxis_gracilis | *Monanthotaxis gracilis* (Hook.f.) P.H.Hoekstra | - | KX786600 | KX786610 | KX786621 | - | KX786640 | | - |
| 552 | Monanthotaxis_heteranth a | *Monanthotaxis heterantha* (Baill.) Verdc. | - | KX761297 | KX787017 | KX786956 | KX761328 | KX786986 | | |
| 553 | Monanthotaxis_hirsuta | *Monanthotaxis hirsuta* (Benth.) P.H.Hoekstra | - | KX761293 | KX787013 | KX786952 | KX761324 | KX786982 | | |
| 554 | Monanthotaxis_klainei_v ar_klainei | *Monanthotaxis klainei* (Engl.) Verdc. var. *klainei* | - | KX761304 | KX787024 | KX786963 | KX761335 | KX786993 | | |
| 555 | Monanthotaxis_klainei_v ar_lastoursvillensis | *Monanthotaxis klainei* var. *lastoursvillensis* (Pellegr.) Verdc. | - | KX761289 | KX787009 | KX786948 | KX761320 | KX786978 | | |
| 556 | Monanthotaxis_komorens is | *Monanthotaxis komorensis* P.H.Hoekstra | - | KX761296 | KX787016 | KX786955 | KX761327 | KX786985 | | |
| 557 | Monanthotaxis_latistamin a | *Monanthotaxis latistamina* P.H.Hoekstra | - | KX761288 | KX787008 | KX786947 | KX761319 | KX786977 | | |
| 558 | Monanthotaxis_maputensi s | *Monanthotaxis maputensis* P.H.Hoekstra | - | KX761308 | KX787028 | KX786967 | KX761339 | KX786997 | | |
| 559 | Monanthotaxis_micrantha | *Monanthotaxis micrantha* (Baker) Verdc. | - | KX761300 | KX787020 | KX786959 | KX761331 | KX786989 | | |

| No. | Voucher information | | GenBank accession numbers | | | | | | | |
|---|---|---|---|---|---|---|---|---|---|---|
| | Tip name in the tree | Taxa name | *atpB-rbcL* | *matK* | *ndhF* | *psbA-trnH* | *rbcL* | *trnL-F* | *trnS-G* | *ycf1* |
| 560 | Monanthotaxis_montana | *Monanthotaxis montana* (Engl. & Diels) P.H.Hoekstra | - | KX786604 | KX786611 | KX786624 | - | KX786643 | | |
| 561 | Monanthotaxis_obovata | *Monanthotaxis obovata* (Benth.) P. H. Hoekstra | - | JQ768579 | JQ768614 | JQ768658 | JQ768698 | JQ768740 | - | |
| 562 | Monanthotaxis_parvifolia | *Monanthotaxis parvifolia* subsp. *kenyensis* Verdc. | - | KX761312 | KX787032 | KX786971 | KX761343 | KX787001 | | - |
| 563 | Monanthotaxis_pellegrinii | *Monanthotaxis pellegrinii* Verdc. | - | KX761302 | KX787022 | KX786961 | KX761333 | KX786991 | | |
| 564 | Monanthotaxis_scamnopetala | *Monanthotaxis scamnopetala* (Exell) P.H.Koekstra | - | KX761286 | KX787006 | KX786945 | KX761317 | KX786975 | | |
| 565 | Monanthotaxis_schweinfurthii_var_schweinfurthii | *Monanthotaxis schweinfurthii* (Engl. & Diels) Verdc. var. *schweinfurthii* | - | KX761313 | KX787033 | - | KX761344 | KX787002 | | |
| 566 | Monanthotaxis_schweinfurthii_var_seretii | *Monanthotaxis schweinfurthii* var. *seretii* (De Wild.) Verdc. | - | KX761294 | KX787014 | KX786953 | KX761325 | KX786983 | | - |
| 567 | Monanthotaxis_schweinfurthii_var_tisserantii | *Monanthotaxis schweinfurthii* var. *tisserantii* (Le Thomas) Verdc. | - | JQ768585 | JQ768621 | JQ768664 | JQ768704 | JQ768746 | | |
| 568 | Monanthotaxis_sororia | *Monanthotaxis sororia* (Diels) Verdc. | - | KX761301 | KX787021 | KX786960 | KX761332 | KX786990 | | |
| 569 | Monanthotaxis_sp_869 | *Monanthotaxis* sp. | - | KX761314 | KX787034 | KX786972 | KX761345 | KX787003 | | |
| 570 | Monanthotaxis_sp_10971 | *Monanthotaxis* sp. 10971 | - | KX786605 | KX786612 | KX786625 | - | KX786644 | | |
| 571 | Monanthotaxis_trichocarpa | *Monanthotaxis trichocarpa* (Engl. & Diels) Verdc. | - | JQ768586 | JQ768622 | JQ768665 | JQ768705 | JQ768747 | | |
| 572 | Monanthotaxis_tripetala | *Monanthotaxis tripetala* P.H.Hoekstra | - | KX761316 | KX787036 | KX786974 | KX761347 | KX787005 | | |
| 573 | Monanthotaxis_velutina | *Monanthotaxis velutina* (Sprague & Hutch.) P.H.Hoekstra | - | KX761292 | KX787012 | KX786951 | KX761323 | KX786981 | | - |
| 574 | Monanthotaxis_whytei | *Monanthotaxis whytei* (Stapf) Verdc. | EF179269 | EF179278 | EF179304 | EF179315 | AY841635 | AY841713 | EF179346 | |

| No. | Voucher information | | GenBank accession numbers | | | | | | | |
|---|---|---|---|---|---|---|---|---|---|---|
| | Tip name in the tree | Taxa name | *atpB-rbcL* | *matK* | *ndhF* | *psbA-trnH* | *rbcL* | *trnL-F* | *trnS-G* | *ycf1* |
| 575 | Monocarpia_euneura | *Monocarpia euneura* Miq. | AY841381 | AY518865 | AY841412 | AY841477 | - | AY319111 | AY841559 | - |
| 576 | Monocarpia_marginalis | *Monocarpia marginalis* (Scheff.) J. Sinclair | - | JQ690397 | JQ690398 | JQ690399 | JQ690395 | JQ690396 | - | JQ690400 |
| 577 | Monocyclanthus_vignei | *Monocyclanthus vignei* Keay | - | - | EU169719 | EU169742 | EU169765 | EU169787 | EU169810 | - |
| 578 | Monodora_angolensis | *Monodora angolensis* Welw. | - | - | EU216648 | EU216672 | - | EU216718 | EU216627 | - |
| 579 | Monodora_brevipes | *Monodora carolinae* Couvreur | - | - | EU216653 | EU216677 | - | EU216723 | EU216632 | - |
| 580 | Monodora_carolinae | *Monodora crispata* Engl. | - | - | EU169720 | EU169743 | AY841637 | AY841715 | EU169811 | - |
| 581 | Monodora_crispata | *Monodora globiflora* Couvreur | - | - | EU216654 | EU216678 | - | EU216724 | EU216633 | - |
| 582 | Monodora_globiflora | *Monodora grandidieri* Baill. | - | - | EU216649 | EU216673 | - | EU216719 | EU216628 | - |
| 583 | Monodora_grandidieri | *Monodora hastipetala* Couvreur | - | - | EU216655 | EU216679 | - | EU216725 | EU216634 | - |
| 584 | Monodora_hastipetala | *Monodora junodii* Engl. & Diels | - | - | EU216651 | EU216675 | - | EU216721 | EU216630 | - |
| 585 | Monodora_junodii | *Monodora laurentii* De Wild. | - | - | EU216650 | EU216674 | - | EU216720 | EU216629 | - |
| 586 | Monodora_laurentii | *Monodora minor* Engl. & Diels | - | - | EU216656 | EU216680 | - | EU216726 | EU216635 | - |
| 587 | Monodora_minor | *Monodora myristica* (Gaertn.) Dunal | EF179270 | AY743485 | EF179305 | DQ125129 | AY743447 | EU216716 | EF179347 | - |
| 588 | Monodora_myristica | *Monodora stenopetala* Oliv. | - | - | EU216652 | EU216676 | - | EU216722 | EU216631 | - |
| 589 | Monodora_stenopetala | *Monodora tenuifolia* Benth. | - | - | EU216647 | EU216671 | AY841638 | AY841716 | EU216626 | - |
| 590 | Monodora_tenuifolia | *Monodoraundulate* (P. Beauv.) Couvreur | - | - | EU169722 | EU169744 | EU169766 | EU169788 | EU169813 | - |
| 591 | Monoon_australe | *Monoon australe* (Benth.) B. Xue & R. M. K. Saunders | - | JX227872 | - | - | JX227897 | JX227849 | - | - |
| 592 | Monoon_borneense | *Monoon borneense* (H. Okada) B. Xue & R.M.K. Saunders | - | JX227873 | - | - | JX227898 | JX227850 | - | - |

| No. | Voucher information | | GenBank accession numbers | | | | | | | |
|---|---|---|---|---|---|---|---|---|---|---|
| | Tip name in the tree | Taxa name | *atpB-rbcL* | *matK* | *ndhF* | *psbA-trnH* | *rbcL* | *trnL-F* | *trnS-G* | *ycf1* |
| | | [= *Polyalthia sinclairiana* I.M. Turner] | | | | | | | | |
| 593 | Monoon_cf_glabrum | *Monoon* cf. *glabrum* (Hook. f. & Thomson) B. Xue & R.M.K. Saunders [= *Polyalthia* cf. *glabra* J. Sinclair] | - | AY518782 | - | - | AY319032 | AY319146 | - | - |
| 594 | Monoon_coffeoides | *Moonon coffeoides* (Thwaites ex Hook. f. & Thomson) B. Xue & R. M. K. Saunders | EU522344 | EU522233 | - | EU522123 | EU522288 | EU522178 | - | - |
| 595 | Monoon_congestum | *Monoon congestum* (Ridl.) B. Xue & R.M.K. Saunders [= *Polyalthia congesta* (Ridl.) J. Sinclair] | - | AY518790 | - | - | AY319019 | AY319133 | - | - |
| 596 | Monoon_cupulare | *Monoon cupulare* (King) B. Xue & R.M.K. Saunders [=*Enicosanthum cupulare* (King) Airy-Shaw] | - | JX227874 | - | - | JX227899 | — | - | - |
| 597 | Monoon_erianthoides | *Monoon erianthoides* (Airy Shaw) B. Xue & R. M. K. Saunders | - | JX227875 | - | - | JX227900 | JX227851 | - | - |
| 598 | Monoon_fuscum | *Monoon fuscum* (King) B. Xue & R. M. K. Saunders | - | AY518787 | JX544779 | JX544792 | - | AY319085 | - | JX544767 |
| 599 | Monoon_hypogaeum | *Monoon hypogaeum* (King) B. Xue & R. M. K. Saunders | - | JX227876 | - | - | JX227901 | JX227852 | - | - |
| 600 | Monoon_kingii | *Monoon kingii* (Baker f.) B. Xue & R. M. K. Saunders | - | JX227877 | - | - | JX227902 | JX227853 | - | - |

| No. | Voucher information | | GenBank accession numbers | | | | | | | |
|---|---|---|---|---|---|---|---|---|---|---|
| | Tip name in the tree | Taxa name | *atpB-rbcL* | *matK* | *ndhF* | *psbA-trnH* | *rbcL* | *trnL-F* | *trnS-G* | *ycf1* |
| 601 | Monoon_klemmei | *Monoon klemmei* (Elmer) B. Xue & R. M. K. Saunders | - | JX227878 | - | - | JX227903 | JX227854 | - | - |
| 602 | Monoon_lateriflorum | *Moonon lateriflorum* (Blume) B. Xue & R. M. K. Saunders | - | AY518781 | JQ723811 | - | JQ723870 | AY319138 | - | JQ723951 |
| 603 | Monoon_laui | *Monoon laui* (Merr.) B. Xue & R. M. K. Saunders | - | JX227879 | - | - | JX227904 | JX227855 | - | - |
| 604 | Monoon_membranifolium | *Monoon membranifolium* (J. Sinclair) B. Xue & R. M. K. Saunders | - | AY518788 | - | - | - | AY319086 | - | - |
| 605 | Monoon_michaeli | *Monoon michaelii* (C.T. White) B. Xue & R. M. K. Saunders | - | JX227880 | - | - | JX227905 | JX227856 | - | - |
| 606 | Monoon_paradoxum | *Monoon paradoxum* (Becc.) B. Xue & R. M. K. Saunders | - | AY518789 | - | - | - | AY319087 | - | - |
| 607 | Monoon_sclerophyllum | *Monoon sclerophyllum* (Hook. f. & Thomson) B. Xue & R.M.K. Saunders [= *Polyalthia sclerophylla* Hook. f. & Thomson] | - | AY518783 | - | - | AY319033 | AY319147 | - | - |
| 608 | Monoon_sympetalum | *Monoon sympetalum* (Merr.) B. Xue & R. M. K. Saunders | - | JX227881 | - | - | JX227906 | JX227857 | - | - |
| 609 | Monoon_xanthopetalum | *Monoon xanthopetalum* (Merr.) B. Xue & R.M.K. Saunders [= *Polyalthia xanthopetala* Merr.] | - | AY518792 | - | - | AY319041 | AY319155 | - | JQ723951 |
| 610 | Monoon_longifolium | *Moonon longifolium* (Sonn.) B. Xue & R. M. K. Saunders | EU522346 | AY518786 | - | EU522125 | EU522290 | EU522180 | - | - |

| No. | Voucher information | | GenBank accession numbers | | | | | | | |
|---|---|---|---|---|---|---|---|---|---|---|
| | Tip name in the tree | Taxa name | *atpB-rbcL* | *matK* | *ndhF* | *psbA-trnH* | *rbcL* | *trnL-F* | *trnS-G* | *ycf1* |
| 611 | Monoon_obtusa | *Moonon obtusum* (Craib) B. Xue & R. M. K. Saunders | - | AY518784 | JX544780 | JX544793 | - | AY319154 | - | JX544768 |
| 612 | Mosannona_costaricensis | *Mosannona costaricensis* (R. E. Fr.) Chatrou | AY841382 | AY743503 | AY841413 | AY841479 | AY743510 | AY743496 | AY841560 | - |
| 613 | Mosannona_discolor | *Mosannona discolor* (R. E. Fr.) Chatrou | - | AY743504 | - | AY841480 | AY743511 | AY743497 | - | - |
| 614 | Mosannona_garwoodii | *Mosannona garwoodii* Chatrou & Welzenis | - | AY743505 | - | AY841481 | AY743512 | AY743498 | - | - |
| 615 | Mosannona_pacifica | *Mosannona pacifica* Chatrou | - | AY743506 | - | AY841482 | AY743513 | AY743499 | - | - |
| 616 | Mosannona_papillosa | *Mosannona papillosa* Chatrou | - | AY743507 | - | AY841483 | AY743514 | AY743500 | - | - |
| 617 | Mosannona_vasquezii | *Mosannona vasquezii* Chatrou | - | AY743508 | - | AY841484 | AY743515 | AY319178 | - | - |
| 618 | Mwasumbia_alba | *Mwasumbia alba* Couvreur & D. M. Johnson | - | - | - | - | EU747680 | EU747674 | - | - |
| 619 | Myristica_fragrans | *Myristica fragrans* Houtt. | - | AJ966803 | AY218188 | - | AF206798 | - | - | - |
| 620 | Neouvaria_acuminatissima | *Neo-uvaria acuminatissima* (Miq.) Airy Shaw | - | AY518793 | - | - | - | AY319112 | - | - |
| 621 | Neouvaria_parallelivenia | *Neo-uvaria parallelivenia* (Boerl.) H. Okada & K. Ueda | - | AY518794 | - | - | - | AY319113 | - | - |
| 622 | Neouvaria_telopea | *Neo-uvaria telopea* Chaowasku | - | JX544751 | JX544778 | JX544791 | JX544755 | JX544783 | - | JX544766 |
| 623 | Neostenanthera_myristicifolia | *Neostenanthera myristicifolia* (Oliv.) Exell | EF179271 | AY743486 | EF179306 | DQ125130 | AY743448 | AY743467 | EF179348 | - |
| 624 | Neostenanthera_platypetala | *Neostenanthera platypetala* (Engl. & Diels) Pellegr | MF322671 | MF322633 | MF322643 | - | MF322648 | MF322661 | - | - |

| No. | Voucher information | | GenBank accession numbers | | | | | | | |
|---|---|---|---|---|---|---|---|---|---|---|
| | Tip name in the tree | Taxa name | *atpB-rbcL* | *matK* | *ndhF* | *psbA-trnH* | *rbcL* | *trnL-F* | *trnS-G* | *ycf1* |
| 625 | Onychopetalum_periquino | *Onychopetalum periquino* (Rusby) D. M. Johnson & N. A. Murray | AY841383 | AY518876 | AY841414 | AY841485 | - | AY319179 | AY841561 | - |
| 626 | Ophrypetalum_odoratum | *Ophrypetalum odoratum* Diels | - | - | EU169723 | EU169745 | EU169767 | EU169789 | EU169814 | - |
| 627 | Orophea_brandisii | *Orophea brandisii* Hook. f. & Thomson | - | AY518813 | - | - | - | AY319116 | - | - |
| 628 | Orophea_celebica | *Orophea celebica* (Blume) Miq. | - | AY518814 | - | - | - | AY319117 | - | - |
| 629 | Orophea_creaghii | *Orophea creaghii* (Ridl.) Leonard á & Keßler | - | AY518817 | - | - | - | AY319118 | - | - |
| 630 | Orophea_enneandra | *Orophea enneandra* Blume | - | AY518816 | - | - | - | AY319120 | - | - |
| 631 | Orophea_enterocarpa | *Orophea enterocarpa* Maingay ex Hook. f. & Thomson | - | AY518815 | JQ690416 | JQ690417 | - | AY319119 | - | JQ690418 |
| 632 | Orophea_kerrii | *Orophea kerrii* Keßler | - | AY518818 | JQ690419 | JQ690420 | - | AY319121 | - | JQ690421 |
| 633 | Orophea_polycarpa | *Orophea polycarpa* A. DC. | - | AY518819 | - | - | - | AY319123 | - | - |
| 634 | Oxandra_asbeckii | *Oxandra asbeckii* (Pulle) R. E. Fr. | - | - | - | AY841486 | AY841639 | AY841717 | - | - |
| 635 | Oxandra_espintana | *Oxandra espintana* (Spruce ex Benth.) Baill. | - | DQ018260 | - | AY841487 | - | AY319180 | - | - |
| 636 | Oxandra_euneura | *Oxandra euneura* Diels | - | - | - | AY841488 | AY841640 | AY841718 | - | - |
| 637 | Oxandra_lanceolata | *Oxandra lanceolata* (Sw.) Baill. | - | KJ012702 | - | KJ426857 | KJ082471 | - | - | - |
| 638 | Oxandra_laurifolia | *Oxandra laurifolia* (Sw.) A. Rich. | - | HM446726 | - | AY841489 | AY841528 | AY841542 | - | - |
| 639 | Oxandra_longipetala | *Oxandra longipetala* R. E. Fr. | - | - | - | AY841490 | AY841641 | AY841719 | - | - |
| 640 | Oxandra_macrophylla | *Oxandra macrophylla* R. E. Fr. | - | - | - | AY841491 | AY841642 | AY841720 | - | - |
| 641 | Oxandra_nitida | *Oxandra nitida* R. E. Fr. | - | - | - | AY841492 | AY841529 | AY841543 | - | - |
| 642 | Oxandra_polyantha | *Oxandra polyantha* R. E. Fr. | - | - | - | AY841493 | AY841643 | AY841721 | - | - |

| No. | Voucher information | | GenBank accession numbers | | | | | | | |
|---|---|---|---|---|---|---|---|---|---|---|
| | Tip name in the tree | Taxa name | *atpB-rbcL* | *matK* | *ndhF* | *psbA-trnH* | *rbcL* | *trnL-F* | *trnS-G* | *ycf1* |
| 643 | Oxandra_riedeliana | *Oxandra riedeliana* R. E. Fr. | - | KP859350 | - | - | - | KP859336 | - | - |
| 644 | Oxandra_sphaerocarpa | *Oxandra sphaerocarpa* R. E. Fr. | - | - | - | AY841494 | AY841644 | AY841722 | - | - |
| 645 | Oxandra_venezuelana | *Oxandra venezuelana* R. E. Fr. | - | JQ690413 | JQ690414 | AY841495 | AY841645 | AY841723 | - | JQ690415 |
| 646 | Oxandra_xylopioides | *Oxandra xylopioides* Diels | - | - | - | AY841496 | AY841646 | AY841724 | - | - |
| 647 | Phaeanthus_ebracteolatus | *Phaeanthus ebracteolatus* (C. Presl) Merr. | - | AY518863 | - | - | JF738713 | AY319125 | - | - |
| 648 | Phaeanthus_sp_WP2A0446 | *Phaeanthus* sp. WP2A0446 | - | GQ248183 | - | GQ248373 | GQ248677 | - | - | - |
| 649 | Phaeanthus_splendens | *Phaeanthus splendens* Miq. | - | AY518864 | JX544777 | JX544790 | JX544754 | AY319126 | - | JX544765 |
| 650 | Phoenicanthus_obliquus | *Phoenicanthus obliquus* (Hook. f. & Thomson) Alston | MF322677 | MF322639 | - | MF322658 | MF322654 | MF322667 | - | - |
| 651 | Piptostigma_mortehani | *Piptostigma mortehani* De Wild. | AY841384 | AY743492 | AY841415 | AY841498 | AY743454 | AY743473 | AY841562 | - |
| 652 | Piptostigma_oyemense | *Piptostigma oyemense* Pellegr. | - | KC627444 | - | KC667604 | KC627994 | - | - | - |
| 653 | Piptostigma_pilosum | *Piptostigma pilosum* Oliv. | - | - | - | AY841499 | AY841648 | AY319181 | - | - |
| 654 | Platymitra_macrocarpa | *Platymitra macrocarpa* Boerl. | - | AY518812 | JQ723809 | JQ690423 | - | AY319127 | - | JQ690424 |
| 655 | Platymitra_sp_TC2012 | *Platymitra* sp. 1 TC-2012 | - | JQ690426 | JQ690427 | JQ690428 | - | JQ690425 | - | JQ690429 |
| 656 | Polyalthia_angustissima | *Polyalthia angustissima* Ridl. | - | JX227882 | JX544831 | JX544807 | JX227907 | JX544795 | - | - |
| 657 | Polyalthia_borneensis | *Polyalthia borneensis* Merr. | - | AY518821 | - | - | AY319014 | AY319128 | - | - |
| 658 | Polyalthia_bullata | *Polyalthia bullata* King | - | JX544825 | JX544839 | JX544809 | JX227908 | JX544800 | - | JX544818 |
| 659 | Polyalthia_cauliflora | *Polyalthia cauliflora* Hook. f. & Thomson | - | AY518823 | JX544837 | - | - | AY319129 | - | - |
| 660 | Polyalthia_celebica | *Polyalthia celebica* Miq. | - | AY518827 | JX544838 | JX544808 | - | AY319130 | - | - |

| No. | Voucher information | | GenBank accession numbers | | | | | | | |
|---|---|---|---|---|---|---|---|---|---|---|
| | Tip name in the tree | Taxa name | *atpB-rbcL* | *matK* | *ndhF* | *psbA-trnH* | *rbcL* | *trnL-F* | *trnS-G* | *ycf1* |
| 661 | Polyalthia_cf_miliusoides | *Polyalthia* cf. *miliusoides* I.M. Turner [as *Polyalthia longipes* (Miq.) Koord. & Valeton in Genbank] | - | AY518829 | - | - | AY319028 | AY319142 | - | - |
| 662 | Polyalthia_cf_subsessilifolia | *Polyalthia* cf. *subsessilifolia* Baker f. | - | JX227884 | - | - | JX227909 | JX227860 | - | - |
| 663 | Polyalthia_cinnamomea | *Polyalthia cinnamomea* (Hook. f. & Thomson) Hook. f. & Thomson | - | AY518828 | - | - | AY319018 | AY319132 | - | - |
| 664 | Polyalthia_debilis | *Polyalthia debilis* (Pierre) Finet & Gagnep. | - | AY518832 | - | - | AY319020 | AY319134 | - | - |
| 665 | Polyalthia_evecta | *Polyalthia evecta* (Pierre) Finet & Gagnep. | - | JX227885 | - | - | JX227910 | JX227861 | - | - |
| 666 | Polyalthia_flagellaris | *Polyalthia flagellaris* (Becc.) Airy-Shaw | - | AY518824 | - | - | AY319022 | AY319136 | - | - |
| 667 | Polyalthia_hispida | *Polyalthia hispida* B. Xue & R. M. K. Saunders | - | JX227886 | - | - | JX227911 | JX227862 | - | - |
| 668 | Polyalthia_insignis | *Polyalthia insignis* (Hook. f.) Airy Shaw | - | JX227887 | - | - | JX227912 | JX227863 | - | - |
| 669 | Polyalthia_johnsonii | *Polyalthia johnsonii* (F. Muell.) B. Xue & R. M. K. Saunders | - | JQ723767 | JQ723791 | JX544810 | JQ723854 | JX544801 | - | JQ723928 |
| 670 | Polyalthia_kanchanaburiana | *Polyalthia kanchanaburiana* Khumch. & Thongp. | - | JX227888 | - | - | JX227913 | JX227864 | - | - |
| 671 | Polyalthia_lanceolata | *Polyalthia lanceolata* S. Vidal | - | JX227889 | - | - | JX227914 | JX227865 | - | - |
| 672 | Polyalthia_lateritia | *Polyalthia lateritia* J. Sinclair | - | JX227890 | - | - | JX227915 | JX227866 | - | - |

| No. | Voucher information | | GenBank accession numbers | | | | | | | |
|---|---|---|---|---|---|---|---|---|---|---|
| | Tip name in the tree | Taxa name | *atpB-rbcL* | *matK* | *ndhF* | *psbA-trnH* | *rbcL* | *trnL-F* | *trnS-G* | *ycf1* |
| 673 | Polyalthia_longirostris | *Polyalthia longirostris* (Scheff.) B. Xue & R. M. K. Saunders | - | AY518826 | - | - | - | AY319091 | - | - |
| 674 | Polyalthia_motleyana | *Polyalthia motleyana* (Hook. f.) Airy Shaw | - | JX227891 | - | - | JX227916 | JX227867 | - | - |
| 675 | Polyalthia_obliqua | *Polyalthia obliqua* Hook. f. & Thomson | - | AY518822 | - | - | AY319029 | AY319143 | - | - |
| 676 | Polyalthia_parviflora | *Polyalthia parviflora* Ridl. | - | JX227892 | JX544836 | - | JX227917 | JX544799 | - | - |
| 677 | Polyalthia_stenopetala | *Polyalthia stenopetala* (Hook. f. & Thomson) Finet & Gagnep. | - | JX227896 | JX544832 | - | JX544823 | AY319148 | - | - |
| 678 | Polyalthia_subcordata | *Polyalthia subcordata* (Blume) Blume | - | AY518830 | - | - | AY319036 | AY319150 | - | - |
| 679 | Polyalthia_suberosa | *Polyalthia suberosa* (Roxb.) Thwaites | AY841386 | AY220439 | AY841417 | AY841502 | AY238956 | AY319152 | AY841564 | JQ723952 |
| 680 | Polyalthia_submontana | *Polyalthia submontana* (Jessup) B. Xue & R. M. K. Saunders | - | JX227893 | - | - | JX227918 | JX227869 | - | - |
| 681 | Polyalthia_trochilia | *Polyalthia trochilia* I. M. Turner | - | JX227894 | - | - | JX227919 | JX227870 | - | MG264593 |
| 682 | Polyalthia_xanthocarpa | *Polyalthia xanthocarpa* B. Xue & R. M. K. Saunders | - | JX227895 | - | - | JX227920 | JX227871 | - | MG264595 |
| 683 | Polyalthiopsis_floribunda_CMU1 | *Polyalthiopsis floribunda* (Jovet-Ast) Chaowasku, Chaowasku 168 | - | MG264583 | MG264588 | MG264570 | MG264580 | MG264575 | - | - |
| 684 | Polyalthiopsis_floribunda_CMU8 | *Polyalthiopsis floribunda* (Jovet-Ast) Chaowasku, Chaowasku 128 | - | MG264585 | MG264590 | MG264572 | - | MG264577 | - | - |
| 685 | Polyceratocarpus_microtrichus | *Polyceratocarpus microtrichus* (Engl. & Diels) Ghesq. ex Pellegr. | - | - | - | - | EU747683 | EU747677 | - | - |

| No. | Voucher information | | GenBank accession numbers | | | | | | | |
|---|---|---|---|---|---|---|---|---|---|---|
| | Tip name in the tree | Taxa name | *atpB-rbcL* | *matK* | *ndhF* | *psbA-trnH* | *rbcL* | *trnL-F* | *trnS-G* | *ycf1* |
| 686 | Polyceratocarpus_parviflorus | *Polyceratocarpus parviflorus* (Baker f.) Ghesq. | - | KC627837 | - | KC688815 | EU747682 | EU747676 | - | - |
| 687 | Polyceratocarpus_pellegrinii | *Polyceratocarpus pellegrinii* Le Thomas | - | - | - | - | EU747684 | EU747678 | - | - |
| 688 | Polyceratocarpus_askhambryan_iringae | *Polyceratocarpus* sp. *ined. (Polyceratocarpus askhambryan-iringae* A. R. Marshall & D. M. Johnson) | - | - | - | - | EU747681 | EU747675 | - | - |
| 689 | Popowia_hirta | *Popowia hirta* Miq. | - | AY518860 | JX544830 | JX544806 | - | AY319156 | - | JX544816 |
| 690 | Popowia_odoardi | *Popowia odoardi* Diels | - | AY518861 | - | - | - | AY319157 | - | - |
| 691 | Popowia_pisocarpa | *Popowia pisocarpa* (Blume) Endl. | - | AY518862 | JQ723812 | - | - | AY319158 | - | JQ723953 |
| 692 | Porcelia_steinbachii | *Porcelia steinbachii* (Diels) R. E. Fr. | - | - | - | - | AY841649 | AY841727 | - | - |
| 693 | Pseudartabotrys_letestui | *Pseudartabotrys letestui* Pellegr. | EF179272 | DQ125061 | EF179307 | DQ125131 | AY841650 | AY841728 | EF179349 | - |
| 694 | Pseudephedranthus_fragrans | *Pseudephedranthus fragrans* (R. E. Fr.) Aristeg. | - | - | - | AY841504 | AY841651 | AY841729 | - | - |
| 695 | Pseudephedranthus_fragrans_Pfa683 | *Pseudephedranthus fragrans* (R. E. Fr.) Aristeg. *Pfa683* | MF322675 | MF322637 | - | MF322656 | MF322652 | MF322665 | - | - |
| 696 | Pseudephedranthus_fragrans_Pfa878 | *Pseudephedranthus fragrans* (R. E. Fr.) Aristeg. *Pfa878* | MF322676 | MF322638 | - | MF322657 | MF322653 | MF322666 | - | - |
| 697 | Pseudomalmea_diclina | *Pseudomalmea diclina* (R. E. Fr.) Chatrou | AY841388 | AY841398 | AY841419 | AY841506 | AY841530 | AY319182 | AY841566 | - |
| 698 | Pseudomalmea_sp_Idarraga_13 | *Pseudomalmea* sp. *Idarraga 13* | - | - | - | AY841507 | AY841652 | AY841730 | - | - |
| 699 | Pseudoxandra_longipes | *Pseudoxandra longipes* Maas | - | - | - | AY841509 | AY841532 | AY841546 | - | - |

| No. | Voucher information | | GenBank accession numbers | | | | | | | |
|---|---|---|---|---|---|---|---|---|---|---|
| | Tip name in the tree | Taxa name | *atpB-rbcL* | *matK* | *ndhF* | *psbA-trnH* | *rbcL* | *trnL-F* | *trnS-G* | *ycf1* |
| 700 | Pseudoxandra_lucida | *Pseudoxandra lucida* R. E. Fr. | AY841389 | AY518870 | AY841420 | AY841510 | - | AY319190 | AY841567 | - |
| 701 | Pseudoxandra_polyphleba | *Pseudoxandra polyphleba* (Diels) R. E. Fr. | - | JQ769091 | JQ769092 | AY841512 | AY841654 | AY841732 | - | - |
| 702 | Pseudoxandra_spiritussancti | *Pseudoxandra spiritus-sancti* Maas | AY841390 | AY841399 | AY841421 | AY841513 | AY841533 | AY841547 | AY841568 | - |
| 703 | Pseuduvaria_acerosa | *Pseuduvaria acerosa* Y. C. F. Su & R. M. K. Saunders | EU522347 | EU522236 | - | EU522126 | EU522291 | EU522181 | - | - |
| 704 | Pseuduvaria_aurantiaca | *Pseuduvaria aurantiaca* (Miq.) Merr. | EU522348 | EU522237 | - | EU522127 | EU522292 | EU522182 | - | - |
| 705 | Pseuduvaria_beccarii | *Pseuduvaria beccarii* (Scheff.) J. Sinclair | EU522349 | EU522238 | - | EU522128 | EU522293 | EU522183 | - | - |
| 706 | Pseuduvaria_borneensis | *Pseuduvaria borneensis* Y. C. F. Su & R. M. K. Saunders | EU522350 | EU522239 | - | EU522129 | EU522294 | EU522184 | - | - |
| 707 | Pseuduvaria_brachyantha | *Pseuduvaria brachyantha* Y. C. F. Su & R. M. K. Saunders | EU522351 | AY518837 | - | EU522130 | EU522295 | AY319160 | - | - |
| 708 | Pseuduvaria_bruneiensis | *Pseuduvaria bruneiensis Y.C.F. Su & R.M.K. Saunders* | EU522352 | EU522241 | - | EU522131 | EU522296 | EU522186 | - | - |
| 709 | Pseuduvaria_calliura | *Pseuduvaria calliura* Airy Shaw | EU522353 | EU522242 | - | EU522132 | EU522297 | EU522187 | - | - |
| 710 | Pseuduvaria_cerina | *Pseuduvaria cerina* J. Sinclair | EU522354 | EU522243 | - | EU522133 | EU522298 | EU522188 | - | - |
| 711 | Pseuduvaria_clemensiae | *Pseuduvaria clemensiae* Y. C. F. Su & R. M. K. Saunders | EU522355 | EU522244 | - | EU522134 | EU522299 | EU522189 | - | - |
| 712 | Pseuduvaria_coriacea | *Pseuduvaria coriacea* Y. C. F. Su & R. M. K. Saunders | EU522356 | AY518838 | - | EU522135 | EU522300 | AY319161 | - | - |
| 713 | Pseuduvaria_costata | *Pseuduvaria costata* (Scheff.) J. Sinclair | EU522357 | EU522246 | - | EU522136 | EU522301 | EU522191 | - | - |

| No. | Voucher information | | GenBank accession numbers | | | | | | | |
|---|---|---|---|---|---|---|---|---|---|---|
| | Tip name in the tree | Taxa name | *atpB-rbcL* | *matK* | *ndhF* | *psbA-trnH* | *rbcL* | *trnL-F* | *trnS-G* | *ycf1* |
| 714 | Pseuduvaria_cymosa | *Pseuduvaria cymosa* (J. Sinclair) Y. C. F. Su & R. M. K. Saunders | EU522358 | EU522247 | - | EU522137 | EU522302 | EU522192 | - | - |
| 715 | Pseuduvaria_dielsiana | *Pseuduvaria dielsiana* (Lauterb.) J. Sinclair | EU522359 | EU522248 | - | EU522138 | EU522303 | EU522193 | - | - |
| 716 | Pseuduvaria_dolichonema | *Pseuduvaria dolichonema* (Diels) J. Sinclair | EU522360 | EU522249 | - | EU522139 | EU522304 | EU522194 | - | - |
| 717 | Pseuduvaria_filipes | *Pseuduvaria filipes* (Lauterb. & K. Schum.) J. Sinclair | EU522361 | EU522250 | - | EU522140 | EU522305 | EU522195 | - | - |
| 718 | Pseuduvaria_fragrans | *Pseuduvaria fragrans* Y. C. F. Su, Chaowasku & R. M. K. Saunders | - | JQ723784 | JX544829 | - | JQ723871 | JQ723924 | - | JQ723954 |
| 719 | Pseuduvaria_froggattii | *Pseuduvaria froggattii* (F. Muell.) Jessup | EU522362 | EU522251 | - | EU522141 | EU522306 | EU522196 | - | - |
| 720 | Pseuduvaria_galeata | *Pseuduvaria galeata* J. Sinclair | EU522363 | EU522252 | - | EU522142 | EU522307 | EU522197 | - | - |
| 721 | Pseuduvaria_glabrescens | *Pseuduvaria glabrescens* (Jessup) Y. C. F. Su & R. M. K. Saunders | EU522364 | EU522253 | - | EU522143 | EU522308 | EU522198 | - | - |
| 722 | Pseuduvaria_grandifolia | *Pseuduvaria grandifolia* (Warb.) J. Sinclair | EU522365 | EU522254 | - | EU522144 | EU522309 | EU522199 | - | - |
| 723 | Pseuduvaria_hylandii | *Pseuduvaria hylandii* Jessup | EU522366 | EU522255 | - | EU522145 | EU522310 | EU522200 | - | - |
| 724 | Pseuduvaria_kingiana | *Pseuduvaria kingiana* Y. C. F. Su & R. M. K. Saunders | EU522367 | EU522256 | - | EU522146 | EU522311 | EU522201 | - | - |
| 725 | Pseuduvaria_latifolia | *Pseuduvaria latifolia* (Blume) Bakh. f. | EU522368 | EU522257 | - | EU522147 | EU522312 | EU522202 | - | - |
| 726 | Pseuduvaria_lignocarpa | *Pseuduvaria lignocarpa* J. Sinclair | EU522369 | EU522258 | - | EU522148 | EU522313 | EU522203 | - | - |

| No. | Voucher information | | GenBank accession numbers | | | | | | | |
|---|---|---|---|---|---|---|---|---|---|---|
| | Tip name in the tree | Taxa name | *atpB-rbcL* | *matK* | *ndhF* | *psbA-trnH* | *rbcL* | *trnL-F* | *trnS-G* | *ycf1* |
| 727 | Pseuduvaria_luzonensis | *Pseuduvaria luzonensis* (Merr.) Y. C. F. Su & R. M. K. Saunders | EU522370 | EU522259 | - | EU522149 | EU522314 | EU522204 | - | - |
| 728 | Pseuduvaria_macgregorii | *Pseuduvaria macgregorii* Merr. | EU522371 | EU522260 | - | EU522150 | EU522315 | EU522205 | - | - |
| 729 | Pseuduvaria_macrocarpa | *Pseuduvaria macrocarpa* (Burck) Y. C. F. Su & R. M. K. Saunders | EU522372 | EU522261 | - | EU522151 | EU522316 | EU522206 | - | - |
| 730 | Pseuduvaria_macrophylla | *Pseuduvaria macrophylla* (Oliv.) Merr. | EU522373 | EU522262 | - | EU522152 | EU522317 | EU522207 | - | - |
| 731 | Pseuduvaria_megalopus | *Pseuduvaria megalopus* (K. Schum.) Y. C. F. Su & Mols 15599 | - | AY518836 | - | - | - | AY319124 | - | - |
| 732 | Pseuduvaria_megalopus_2 | *Pseuduvaria megalopus (*K. Schum.) Y. C. F. Su & Mols 16235 | EU522374 | EU522263 | - | EU522153 | EU522318 | EU522208 | - | - |
| 733 | Pseuduvaria_mindorensis | *Pseuduvaria mindorensis* Y. C. F. Su & R. M. K. Saunders | EU522375 | EU522264 | - | EU522154 | EU522319 | EU522209 | - | - |
| 734 | Pseuduvaria_mollis | *Pseuduvaria mollis* (Warb.) J. Sinclair | EU522376 | EU522265 | - | EU522155 | EU522320 | EU522210 | - | - |
| 735 | Pseuduvaria_monticola | *Pseuduvaria monticola* J. Sinclair | EU522377 | EU522266 | - | EU522156 | EU522321 | EU522211 | - | - |
| 736 | Pseuduvaria_mulgraveana | *Pseuduvaria mulgraveana* Jessup | EU522378 | EU522267 | - | EU522157 | EU522322 | EU522212 | - | - |
| 737 | Pseuduvaria_multiovulata | *Pseuduvaria multiovulata* (C. E. C. Fisch.) J. Sinclair | EU522379 | EU522268 | - | EU522158 | EU522323 | EU522213 | - | - |
| 738 | Pseuduvaria_novaguineensis | *Pseuduvaria nova-guineensis* J. Sinclair | EU522380 | EU522269 | - | EU522159 | EU522324 | EU522214 | - | - |
| 739 | Pseuduvaria_obliqua | *Pseuduvaria obliqua* Y. C. F. Su & R. M. K. Saunders | EU522381 | EU522270 | - | EU522160 | EU522325 | EU522215 | - | - |

| No. | Voucher information | | GenBank accession numbers | | | | | | | |
|---|---|---|---|---|---|---|---|---|---|---|
| | Tip name in the tree | Taxa name | *atpB-rbcL* | *matK* | *ndhF* | *psbA-trnH* | *rbcL* | *trnL-F* | *trnS-G* | *ycf1* |
| 740 | Pseuduvaria_oxycarpa | *Pseuduvaria oxycarpa* (Boerl. ex Koord.) Y. C. F. Su & R. M. K. Saunders | EU522382 | EU522271 | - | EU522161 | EU522326 | EU522216 | - | - |
| 741 | Pseuduvaria_pamattonis | *Pseuduvaria pamattonis* (Miq.) Y. C. F. Su & R. M. K. Saunders | EU522383 | AY518840 | - | EU522162 | EU522327 | AY319163 | - | - |
| 742 | Pseuduvaria_parvipetala | *Pseuduvaria parvipetala* Y. C. F. Su & R. M. K. Saunders | EU522384 | EU522273 | - | EU522163 | EU522328 | EU522218 | - | - |
| 743 | Pseuduvaria_philippinensis | *Pseuduvaria philippinensis* Merr. | EU522385 | EU522274 | - | EU522164 | EU522329 | EU522219 | - | - |
| 744 | Pseuduvaria_phuyensis | *Pseuduvaria phuyensis* (R. M. K. Saunders, Y. C. F. Su & Chalermglin) Y. C. F. Su & R. M. K. Saunders | EU522342 | AY518841 | - | EU522121 | EU522287 | AY319114 | - | - |
| 745 | Pseuduvaria_reticulata | *Pseuduvaria reticulata* (Blume) Miq. | EU522386 | EU522275 | - | EU522165 | EU522330 | EU522220 | - | - |
| 746 | Pseuduvaria_rugosa | *Pseuduvaria rugosa* (Blume) Merr. | EU522387 | AY518839 | - | EU522166 | EU522331 | AY319162 | - | - |
| 747 | Pseuduvaria_sessilicarpa | *Pseuduvaria sessilicarpa* (J. Sinclair) Y. C. F. Su & R. M. K. Saunders | EU522388 | EU522277 | - | EU522167 | EU522332 | EU522222 | - | - |
| 748 | Pseuduvaria_sessilifolia | *Pseuduvaria sessilifolia* J. Sinclair | EU522389 | EU522278 | - | EU522168 | EU522333 | EU522223 | - | - |
| 749 | Pseuduvaria_setosa | *Pseuduvaria setosa* (King) J. Sinclair | EU522390 | EU522279 | - | EU522169 | EU522334 | EU522224 | - | - |
| 750 | Pseuduvaria_silvestris | *Pseuduvaria silvestris* (Diels) J. Sinclair | EU522391 | EU522280 | - | EU522170 | EU522335 | EU522225 | - | - |
| 751 | Pseuduvaria_subcordata | *Pseuduvaria subcordata* Y. C. F. Su & R. M. K. Saunders | EU522392 | EU522281 | - | EU522171 | EU522336 | EU522226 | - | - |
| 752 | Pseuduvaria_taipingensis | *Pseuduvaria taipingensis* J. Sinclair | EU522393 | EU522282 | - | EU522172 | EU522337 | EU522227 | - | - |

| No. | Voucher information | | GenBank accession numbers | | | | | | | |
|---|---|---|---|---|---|---|---|---|---|---|
| | Tip name in the tree | Taxa name | *atpB-rbcL* | *matK* | *ndhF* | *psbA-trnH* | *rbcL* | *trnL-F* | *trnS-G* | *ycf1* |
| 753 | Pseuduvaria_trimera | *Pseuduvaria trimera* (Craib) Y. C. F. Su & R. M. K. Saunders | EU522394 | EU522283 | - | EU522173 | EU522338 | EU522228 | - | - |
| 754 | Pseuduvaria_unguiculata | *Pseuduvaria unguiculata* (Elmer) Y. C. F. Su & R. M. K. Saunders | EU522395 | EU522284 | - | EU522174 | EU522339 | EU522229 | - | - |
| 755 | Pseuduvaria_villosa | *Pseuduvaria villosa* Jessup | EU522396 | EU522285 | - | EU522175 | EU522340 | EU522230 | - | - |
| 756 | Pyramidanthe_prismatica | *Pyramidanthe prismatica* (Hook. f. & Thomson) J. Sinclair | - | JN175163 | - | JN175178 | JN175193 | JN175208 | - | - |
| 757 | Ruizodendron_ovale | *Ruizodendron ovale* (Ruiz & Pav.) R. E. Fr. | - | - | - | AY841514 | AY841657 | AY841735 | - | - |
| 758 | Sageraea_lanceolata | *Sageraea lanceolata* Miq. | - | AY518799 | JX544774 | JX544787 | - | AY319164 | - | JX544762 |
| 759 | Sanrafaelia_ruffonammari | *Sanrafaelia ruffonammari* Verdc. | - | - | EU169724 | EU169746 | EU169768 | EU169790 | EU169815 | - |
| 760 | Sapranthus_microcarpus | *Sapranthus microcarpus* (Donn. Sm.) R. E. Fr. | - | AY518806 | - | - | - | AY319166 | - | - |
| 761 | Sapranthus_palanga | *Sapranthus palanga* R. E. Fr. | - | JQ586518 | - | - | JQ590191 | - | - | - |
| 762 | Sapranthus_viridiflorus | *Sapranthus viridiflorus* G. E. Schatz | AY841391 | AY743493 | AY841422 | AY841515 | JQ590194 | AY319165 | AY841569 | JQ723955 |
| 763 | Schefferomitra_subaequalis | *Schefferomitra subaequalis* (Scheff.) Diels | - | KX786606 | - | KX786626 | KX786628 | KX786645 | - | - |
| 764 | Sirdavidia_solannona | *Sirdavidia solannona* Couvreur & Sauquet | - | - | - | - | KP144082 | KP144080 | - | - |
| 765 | Sphaerocoryne_affinis | *Sphaerocoryne affinis* (Teijsm. & Binn.) Ridl. | - | KM924852 | KM924886 | KM924990 | - | KM924953 | | |
| 766 | Sphaerocoryne_blanfordiana | *Sphaerocoryne blanfordiana* C.E.C.Fisch. | - | KM924853 | KM924887 | KM924991 | - | KM924954 | | |

| No. | Voucher information | | GenBank accession numbers | | | | | | | |
|---|---|---|---|---|---|---|---|---|---|---|
| | Tip name in the tree | Taxa name | *atpB-rbcL* | *matK* | *ndhF* | *psbA-trnH* | *rbcL* | *trnL-F* | *trnS-G* | *ycf1* |
| 767 | Sphaerocoryne_gracilipes | *Sphaerocoryne gracilipes* (Benth.) X.Guo & R.M.K.Saunders= Friesodielsia gracilipes (Benth.) Steenis | - | KX786599 | KX786609 | KX786620 | - | KX786639 | | |
| 768 | Sphaerocoryne_gracilis | *Sphaerocoryne gracilis* (Oliv. ex Engl. & Diels) Verdc. | - | - | JQ768623 | EU169732 | EU169755 | EU169777 | EU169799 | - |
| 769 | Sphaerocoryne_sp_DCT2012 | *Sphaerocoryne* sp. 07/4 | - | AY518878 | JQ768624 | FJ743788 | AY319071 | AY319185 | - | - |
| 770 | Stelechocarpus_burahol | *Stelechocarpus burahol* (Blume) Hook. f. & Thomson | - | AY518803 | JX544775 | JX544788 | - | AY319167 | - | JQ723956 |
| 771 | Stenanona_cauliflora | *Stenanona cauliflora* (J. W. Walker) G. E. Schatz | | KU727305 | - | KU727347 | KU727386 | KU727424 | - | - |
| 772 | Stenanona_costaricensis | *Stenanona costaricensis* R. E. Fr. | - | AY518801 | JX544772 | AY841516 | JQ590198 | AY319183 | - | JX544759 |
| 773 | Stenanona_flagelliflora | *Stenanona flagelliflora* T. Wendt & G.E. Schatz | - | KU727307 | - | KU727349 | KU727388 | KU727426 | - | - |
| 774 | Stenanona_hondurensis | *Stenanona hondurensis* G. E. Schatz, F. G. Coe & Maas | - | KU727311 | - | KU727353 | KU727392 | KU727430 | - | - |
| 775 | Stenanona_humilis | *Stenanona humilis* (Miranda) G. E. Schatz | - | KU727313 | - | KU727355 | KU727394 | KU727432 | - | - |
| 776 | Stenanona_migueliana | *Stenanona migueliana* Ortiz-Rodr. & G.E. Schatz | - | KU727316 | - | KU727358 | KU727397 | KU727435 | - | - |
| 777 | Stenanona_monticola | *Stenanona monticola* Maas & G. E. Schatz | - | KU727321 | - | KU727363 | KU727402 | KU727440 | - | - |
| 778 | Stenanona_panamensis | *Stenanona panamensis* Standl. | - | AY518802 | - | - | - | AY319184 | - | - |

| No. | Voucher information | | GenBank accession numbers | | | | | | | |
|---|---|---|---|---|---|---|---|---|---|---|
| | Tip name in the tree | Taxa name | *atpB-rbcL* | *matK* | *ndhF* | *psbA-trnH* | *rbcL* | *trnL-F* | *trnS-G* | *ycf1* |
| 779 | Stenanona_sp_A | *Stenanona* sp. A | - | KU727322 | - | KU727364 | — | KU727441 | - | - |
| 780 | Stenanona_stenopetala | *Stenanona stenopetala* (Donn. Sm.) G.E. Schatz | - | KU727324 | - | KU727366 | KU727404 | KU727443 | - | - |
| 781 | Stenanona_tuberculata | *Stenanona tuberculata* G.E. Schatz & Maas | - | KU727327 | - | KU727369 | - | KU727446 | - | - |
| 782 | Tetrameranthus_duckei | *Tetrameranthus duckei* R. E. Fr. | - | - | - | AY841439 | AY841658 | AY841736 | - | - |
| 783 | Tetrameranthus_laomae | *Tetrameranthus laomae* D. R. Simpson | - | - | - | - | AY841659 | AY841737 | - | - |
| 784 | Toussaintia_orientalis | *Toussaintia orientalis* Verdc. | - | - | EU169710 | EU169733 | EU169756 | EU169778 | EU169800 | - |
| 785 | Tridimeris_hahniana | *Tridimeris hahniana* (Baill.) Baill. | - | - | - | - | - | AY319169 | - | - |
| 786 | Tridimeris_sp_1_TC2012 | *Tridimeris* sp. TC-2012 | - | JX544750 | JX544773 | JX544786 | JX544753 | JX544782 | - | JX544761 |
| 787 | Tridimeris_tuxtlensis | *Tridimeris tuxtlensis* G.E. Schatz ind. | | KU727329 | - | - | - | - | - | - |
| 788 | Trigynaea_duckei | *Trigynaea duckei* (R. E. Fr.) R. E. Fr. | - | - | - | - | AY841660 | AY841738 | - | - |
| 789 | Trigynaea_lanceipetala | *Trigynaea lanceipetala* D. M. Johnson & N. A. Murray | EF179274 | AY743487 | EF179309 | - | AY743449 | AY743468 | EF179351 | - |
| 790 | Trivalvaria_costata | *Trivalvaria costata* (Hook. f. & Thomson) I. M. Turner | - | - | JQ723815 | - | - | - | - | JQ723957 |
| 791 | Trivalvaria_costata_2 | *Trivalvaria costata (Hook. f. & Thomson) I. M. Turner* | - | HQ286574 | - | - | HQ286580 | HQ286587 | - | - |
| 792 | Trivalvaria_macrophylla | *Trivalvaria macrophylla* (Blume) Miq. | - | HQ286576 | - | - | HQ286582 | HQ286588 | - | - |
| 793 | Trivalvaria_sp_TC2012 | *Trivalvaria* sp. 1 TC-2012 | - | JX544824 | JX544828 | JX544805 | JX544822 | JX544794 | - | JX544814 |
| 794 | Unonopsis_pittieri | *Unonopsis pittieri* Saff. | - | GQ982122 | - | AY841517 | AY841661 | DQ018201 | - | - |

| No. | Voucher information | | GenBank accession numbers | | | | | | | |
|---|---|---|---|---|---|---|---|---|---|---|
| | Tip name in the tree | Taxa name | *atpB-rbcL* | *matK* | *ndhF* | *psbA-trnH* | *rbcL* | *trnL-F* | *trnS-G* | *ycf1* |
| 795 | Unonopsis_stipitata | *Unonopsis stipitata* Diels | AY841392 | AY841400 | AY841423 | AY841519 | AY841662 | DQ018202 | AY841570 | - |
| 796 | Uvaria_acuminata | *Uvaria acuminata* Oliv. | - | FJ743753 | - | FJ743791 | FJ743825 | FJ743860 | - | - |
| 797 | Uvaria_afzelii | *Uvaria afzelii* G. Elliot | - | JN175164 | - | JN175179 | JN175194 | JN175209 | - | - |
| 798 | Uvaria_angolensis | *Uvaria angolensis* Welw. ex Oliv. | - | FJ743754 | - | FJ743792 | FJ743826 | FJ743861 | - | - |
| 799 | Uvaria_anonoides | *Uvaria anonoides* Baker f. | - | FJ743755 | - | FJ743793 | FJ743827 | FJ743862 | - | - |
| 800 | Uvaria_antsiranensis | *Uvaria antsiranensis* Le Thomas | - | JN175165 | - | JN175180 | JN175195 | JN175210 | - | - |
| 801 | Uvaria_argenteum | *Uvaria argenteum* Blume | - | FJ743741 | - | FJ743778 | FJ743816 | FJ743850 | - | - |
| 802 | Uvaria_baumannii | *Uvaria baumannii* Engl. & Diels | - | FJ743756 | - | FJ743794 | FJ743828 | FJ743863 | - | - |
| 803 | Uvaria_boniana | *Uvaria boniana* Finet & Gagnep. | - | FJ743757 | - | FJ743795 | FJ743829 | FJ743864 | - | - |
| 804 | Uvaria_borneensis | *Uvaria borneensis* (Merr.) Utteridge | - | FJ743758 | - | FJ743796 | FJ743830 | FJ743865 | - | - |
| 805 | Uvaria_buchholzii | *Uvaria buchholzii* Engl. & Diels | - | GU951715 | - | GU951724 | GU951733 | GU951742 | - | - |
| 806 | Uvaria_calamistrata | *Uvaria calamistrata* Hance | - | FJ743759 | - | FJ743797 | FJ743831 | FJ743866 | - | - |
| 807 | Uvaria_chamae | *Uvaria chamae* P. Beauv. | - | FJ743760 | - | FJ743798 | AY841663 | AY841741 | - | - |
| 808 | Uvaria_cherrevensis | *Uvaria cherrevensis* (Pierre ex Finet & Gagnep.) L. L. Zhou, Y. C. F. Su & R. M. K. Saunders | - | FJ743750 | - | FJ743787 | FJ743823 | FJ743858 | - | - |
| 809 | Uvaria_clavata | *Uvaria clavata* Pierre ex Engl. & Diels | - | FJ743761 | - | FJ743799 | FJ743833 | FJ743867 | - | - |
| 810 | Uvaria_commersoniana | *Uvaria commersoniana* Baill. | - | JN175166 | - | JN175181 | JN175196 | JN175211 | - | - |
| 811 | Uvaria_concava | *Uvaria concava* Teijsm. & Binn. | - | JN175167 | - | JN175182 | JN175197 | JN175212 | - | - |
| 812 | Uvaria_cordata | *Uvaria cordata* (Dunal) Alston | - | JN175168 | - | JN175183 | JN175198 | JN175213 | - | - |

| No. | Voucher information | | GenBank accession numbers | | | | | | | |
|---|---|---|---|---|---|---|---|---|---|---|
| | Tip name in the tree | Taxa name | *atpB-rbcL* | *matK* | *ndhF* | *psbA-trnH* | *rbcL* | *trnL-F* | *trnS-G* | *ycf1* |
| 813 | Uvaria_cuneifolia | *Uvaria cuneifolia* (Hook.f. & Thomson) L. L. Zhou, Y. C. F. Su & R. M. K. Saunders | - | FJ743749 | - | FJ743786 | FJ743822 | FJ743857 | - | - |
| 814 | Uvaria_dasoclema | *Uvaria dasoclema* L. L. Zhou, Y. C. F. Su & R. M. K. Saunders | - | GU951716 | - | GU951725 | GU951734 | GU951743 | - | - |
| 815 | Uvaria_dependens | *Uvaria dependens* Engl. & Diels | - | JN175169 | - | JN175184 | JN175199 | JN175214 | - | - |
| 816 | Uvaria_dulcis | *Uvaria dulcis* Dunal | - | FJ743740 | - | FJ743777 | FJ743815 | FJ743849 | - | - |
| 817 | Uvaria_elmeri | *Uvaria elmeri* Merr. | - | JN175170 | - | JN175185 | JN175200 | JN175215 | - | - |
| 818 | Uvaria_excelsum | *Uvaria excelsum* (Hook.f. & Thomson) King | - | FJ743742 | - | FJ743779 | FJ743817 | FJ743851 | - | - |
| 819 | Uvaria_faulknerae | *Uvaria faulknerae* Verdc. | - | FJ743762 | - | FJ743800 | FJ743834 | FJ743868 | - | - |
| 820 | Uvaria_flava | *Uvaria flava* Teijsm. & Binn. | - | FJ743763 | - | FJ743801 | FJ743835 | FJ743869 | - | - |
| 821 | Uvaria_glabrum | *Uvaria glabra* Span | - | FJ743743 | - | FJ743780 | FJ743818 | FJ743852 | - | - |
| 822 | Uvaria_grandiflora | *Uvaria grandiflora* Roxb. ex Hornem. | - | FJ743764 | - | FJ743802 | FJ743836 | FJ743870 | - | - |
| 823 | Uvaria_hahnii | *Uvaria griffithii L. L. Zhou, Y. C. F. Su & R. M. K. Saunders* | - | FJ743746 | - | FJ743783 | AY841607 | AY841685 | - | - |
| 824 | Uvaria_hamiltonii | *Uvaria hahnii* (Finet & Gagnep.) J. Sinclair | - | JN175171 | - | JN175186 | JN175201 | JN175216 | - | - |
| 825 | Uvaria_hirsuta | *Uvaria hamiltonii* Hook. f. & Thomson | - | FJ743765 | - | FJ743803 | FJ743837 | FJ743871 | - | - |
| 826 | Uvaria_hookeri | *Uvaria hirsuta* Jack | - | FJ743766 | - | FJ743804 | FJ743838 | FJ743872 | - | - |
| 827 | Uvaria_laha | *Uvaria hookeri* King | - | FJ743744 | - | FJ743781 | AY841606 | AY841684 | - | - |
| 828 | Uvaria_leichhardtii | *Uvaria laha* Miq. | - | JN175172 | - | JN175187 | JN175202 | JN175217 | - | - |

| No. | Voucher information | | GenBank accession numbers | | | | | | | |
|---|---|---|---|---|---|---|---|---|---|---|
| | Tip name in the tree | Taxa name | *atpB-rbcL* | *matK* | *ndhF* | *psbA-trnH* | *rbcL* | *trnL-F* | *trnS-G* | *ycf1* |
| 829 | Uvaria_leptocladon | *Uvaria leichhardtii* (F. Muell.) L. L. Zhou, Y. C. F. Su & R. M. K. Saunders | - | GU951717 | - | GU951726 | GU951735 | GU951744 | - | - |
| 830 | Uvaria_lobbiana | *Uvaria leptocladon* Oliv. | - | FJ743767 | - | FJ743805 | FJ743839 | FJ743873 | - | - |
| 831 | Uvaria_lucida | *Uvaria lobbiana* Hook. f. & Thomson | - | FJ743768 | - | FJ743806 | FJ743840 | FJ743874 | - | - |
| 832 | Uvaria_lurida | *Uvaria lucida* Benth. | EF179275 | AY238966 | EF179310 | AY841440 | AY238957 | EF179319 | EF179352 | - |
| 833 | Uvaria_macropoda | *Uvaria lurida* Hook. f. & Thomson | - | FJ743769 | - | FJ743807 | FJ743841 | FJ743875 | - | - |
| 834 | Uvaria_manjensis | *Uvaria macropoda* Hook. f. & Thomson | - | FJ743770 | - | FJ743808 | FJ743842 | FJ743876 | - | - |
| 835 | Uvaria_marenteria | *Uvaria manjensis* Cavaco & Keraudren | - | JN175173 | - | JN175188 | JN175203 | JN175218 | - | - |
| 836 | Uvaria_micranthum | *Uvaria marenteria* (DC.) Baill. | - | JN175174 | - | JN175189 | JN175204 | JN175219 | - | - |
| 837 | Uvaria_pandensis | *Uvaria micrantha* (A.DC.) Hook. f. & Thomson | - | FJ743745 | - | FJ743782 | FJ743819 | FJ743854 | - | - |
| 838 | Uvaria_pauciovulata | *Uvaria pandensis* Verdc. | - | FJ743771 | - | FJ743809 | FJ743843 | FJ743877 | - | - |
| 839 | Uvaria_rufa | *Uvaria pauciovulata* Hook. f. & Thomson | - | JN175175 | - | JN175190 | JN175205 | JN175220 | - | - |
| 840 | Uvaria_rupestris | *Uvaria rufa* Blume | - | FJ743772 | - | FJ743810 | FJ743844 | FJ743878 | - | - |
| 841 | Uvaria_sambiranensis | *Uvaria rupestris* (Jessup) L. L. Zhou, Y. C. F. Su & R. M. K. Saunders | - | GU951718 | - | GU951727 | GU951736 | GU951745 | - | - |
| 842 | Uvaria_sankowskyi | *Uvaria sambiranensis* Deroin & L. Gaut. | - | JN175176 | - | JN175191 | JN175206 | JN175221 | - | - |
| 843 | Uvaria_scabridula | *Uvaria sankowskyi* L. L. Zhou, Y. C. F. Su & R. M. K. Saunders | - | GU951719 | - | GU951728 | GU951737 | GU951746 | - | - |

| No. | Voucher information | | GenBank accession numbers | | | | | | | |
|---|---|---|---|---|---|---|---|---|---|---|
| | Tip name in the tree | Taxa name | *atpB-rbcL* | *matK* | *ndhF* | *psbA-trnH* | *rbcL* | *trnL-F* | *trnS-G* | *ycf1* |
| 844 | Uvaria_semecarpifolia | *Uvaria scabridula* (Jessup) L. L. Zhou, Y. C. F. Su & R. M. K. Saunders | - | GU951720 | - | GU951729 | GU951738 | GU951747 | - | - |
| 845 | Uvaria_siamensis | *Uvaria semecarpifolia* Hook. f. & Thomson | - | FJ743773 | - | FJ743811 | FJ743845 | FJ743879 | - | - |
| 846 | Uvaria_sphenocarpa | *Uvaria siamensis* (Scheff.) L. L. Zhou, Y. C. F. Su & R. M. K. Saunders | - | FJ743752 | - | FJ743790 | FJ743824 | FJ743859 | - | - |
| 847 | Uvaria_topazensis | *Uvaria sphenocarpa* Hook. f. & Thomson | - | FJ743774 | - | FJ743812 | FJ743846 | FJ743880 | - | - |
| 848 | Uvaria_uhrii | *Uvaria topazensis* (Jessup) L. L. Zhou, Y. C. F. Su & R. M. K. Saunders | - | GU951721 | - | GU951730 | GU951739 | GU951748 | - | - |
| 849 | Uvaria_unguiculata | *Uvaria uhrii* (F. Muell.) L. L. Zhou, Y. C. F. Su & R. M. K. Saunders | - | GU951722 | - | GU951731 | GU951740 | GU951749 | - | - |
| 850 | Uvaria_versicolor | *Uvaria unguiculata* (Jessup) L. L. Zhou, Y. C. F. Su & R. M. K. Saunders | - | GU951723 | - | GU951732 | GU951741 | GU951750 | - | - |
| 851 | Uvaria_viridiflorum | *Uvaria versicolor* Pierre ex Engl. & Diels | - | FJ743775 | - | FJ743813 | FJ743847 | FJ743881 | - | - |
| 852 | Uvaria_welwitschii | *Uvaria welwitschii* (Hiern) Engl. & Diels | - | JN175177 | - | JN175192 | JN175207 | JN175222 | - | - |
| 853 | Uvaria_wrayi | *Uvaria wrayi* (King) L. L. Zhou, Y. C. F. Su & R. M. K. Saunders | - | FJ743747 | - | FJ743784 | FJ743821 | FJ743856 | - | - |

| No. | Voucher information | | GenBank accession numbers | | | | | | | |
|---|---|---|---|---|---|---|---|---|---|---|
| | Tip name in the tree | Taxa name | *atpB-rbcL* | *matK* | *ndhF* | *psbA-trnH* | *rbcL* | *trnL-F* | *trnS-G* | *ycf1* |
| 854 | Uvaria_zeylanica | *Uvaria zeylanica* Aubl. ex Willd. | - | FJ743776 | - | FJ743814 | FJ743848 | FJ743882 | - | - |
| 855 | Uvariastrum_insculptum | *Uvariastrum insculptum* (Engl. & Diels) Sprague & Hutch. | - | - | EU169725 | EU169747 | EU169769 | EU169791 | - | - |
| 856 | Uvariastrum_pierreanum | *Uvariastrum pierreanum* Engl. | - | - | - | EU169748 | EU169770 | EU169792 | EU169816 | - |
| 857 | Uvariastrum_pynaertii | *Uvariastrum pynaertii* De Wild. | - | KC627652 | - | KC667802 | KC628249 | - | - | - |
| 858 | Uvariodendron_connivens | *Uvariodendron connivens* (Benth.) R. E. Fr. | - | - | - | KC688780 | - | - | - | - |
| 859 | Uvariodendron_kirkii | *Uvariodendron kirkii* Verdc. | - | - | EU169726 | EU169749 | EU169771 | EU169793 | EU169817 | - |
| 860 | Uvariodendron_molundense | *Uvariodendron molundense* (Diels) R. E. Fr. | - | - | EU169727 | EU169750 | EU169772 | EU169794 | EU169818 | - |
| 861 | Uvariodendron_sp_PM4812 | *Uvariodendron* sp. PM4812 | - | KC627429 | - | KC667591 | KC627977 | - | - | - |
| 862 | Uvariopsis_bakeriana | *Uvariopsis bakeriana* (Hutch. & Dalziel) Robyns & Ghesq. | - | KC627573 | - | KC667743 | KC628178 | - | - | - |
| 863 | Uvariopsis_korupensis | *Uvariopsis korupensis* Gereau & Kenfack | - | KC627757 | EU169729 | EU169752 | EU169774 | EU169796 | EU169820 | - |
| 864 | Uvariopsis_tripetala | *Uvariopsis tripetala* (Baker f.) G. E. Schatz | - | - | EU169712 | EU169735 | EU169758 | EU169780 | EU169802 | - |
| 865 | Uvariopsis_vanderystii | *Uvariopsis vanderystii* Robyns & Ghesq. | - | - | EU169728 | EU169751 | EU169773 | EU169795 | EU169819 | - |
| 866 | Wangia_florulenta_73371 | *Wangia florulenta (C.Y.Wu ex P.T.Li) B.Xue* | - | KX495154 | KX495158 | | KX495156 | KX495162 | | KX495159 |
| 867 | Wangia_saccopetaloides_17336 | *Wangia saccopetaloides* (W. T. Wang) X. Guo & R. M. K. Saunders | - | KF680920 | KF680923 | KF680924 | KF680926 | KF680930 | - | KF680932 |

| No. | Voucher information | | GenBank accession numbers | | | | | | | |
|---|---|---|---|---|---|---|---|---|---|---|
| | Tip name in the tree | Taxa name | *atpB-rbcL* | *matK* | *ndhF* | *psbA-trnH* | *rbcL* | *trnL-F* | *trnS-G* | *ycf1* |
| 868 | Winitia_cauliflorus | *Winitia cauliflora* (Scheff.) Chaowasku | - | AY518800 | JX544776 | JX544789 | - | AY319168 | - | JX544764 |
| 869 | Winitia_expansa | *Winitia expansa* Chaowasku | - | KC857617 | KC857618 | KC857619 | - | KC857616 | - | KC857620 |
| 870 | Wuodendron_praecox_C108 | *Wuodendron praecox* (Hook. f. & Thomson) B.Xue, Y.H.Tan & X.L.Hou | - | MF687367 | MF687369 | MF687371 | MF687373 | MF687375 | | MF687377 |
| 871 | Wuodendron_praecox_XB190 | *Wuodendron praecox* (Hook. f. & Thomson) B.Xue, Y.H.Tan & X.L.Hou | - | JX544749 | JX544770 | JX544785 | JX544752 | JX544781 | | JX544757 |
| 872 | Xylopia_acutiflora | *Xylopia acutiflora* (Dunal) A. Rich. | - | KC627830 | - | KC668016 | KC627989 | - | - | - |
| 873 | Xylopia_aethiopica | *Xylopia aethiopica* (Dunal) A. Rich. | - | KC627474 | - | KC667638 | KC628042 | - | - | - |
| 874 | Xylopia_ambanjensis | *Xylopia ambanjensis* Cavaco & Keraudren | - | - | KX998992 | KX999026 | - | KX999065 | - | - |
| 875 | Xylopia_aromatica | *Xylopia aromatica* (Lam.) Mart. | - | KP052711 | KP052719 | KP052727 | - | KP052735 | - | - |
| 876 | Xylopia_aurantiiodora | *Xylopia aurantiiodora* De Wild. & T. Durand | - | - | - | KX999027 | - | KX999066 | - | - |
| 877 | Xylopia_bemarivensis | *Xylopia bemarivensis* Diels | - | - | - | KX999028 | - | KX999067 | - | - |
| 878 | Xylopia_benthamii | *Xylopia benthamii* R. E. Fr. | - | KX998972 | KX998993 | KX999029 | - | KX999068 | - | - |
| 879 | Xylopia_bocatorena | *Xylopia bocatorena* Schery | - | KP052712 | KP052720 | KP052728 | - | KP052736 | - | - |
| 880 | Xylopia_buxifolia | *Xylopia buxifolia* Baill. | - | KP052735 | KP052721 | KP052729 | - | KP052737 | - | - |
| 881 | Xylopia_calophylla | *Xylopia calophylla* R. E. Fr. | - | KX998973 | KX998994 | - | - | - | - | - |
| 882 | Xylopia_cupularis | *Xylopia cupularis* Mildbr. | - | KM924855 | KM924895 | KM924993 | KM924923 | KM924956 | - | - |
| 883 | Xylopia_erythrodactyla | *Xylopia erythrodactyla* D. M. Johnson & N. A. Murray | - | KX998974 | KX998995 | KX999030 | - | KX999069 | - | - |

| No. | Voucher information | | GenBank accession numbers | | | | | | | |
|---|---|---|---|---|---|---|---|---|---|---|
| | Tip name in the tree | Taxa name | *atpB-rbcL* | *matK* | *ndhF* | *psbA-trnH* | *rbcL* | *trnL-F* | *trnS-G* | *ycf1* |
| 884 | Xylopia_ferruginea | *Xylopia ferruginea* (Hook. f. & Thomson) Hook. f. & Thomson | - | DQ125063 | EF179311 | DQ125133 | AY841666 | AY841744 | - | - |
| 885 | Xylopia_flamignii | *Xylopia flamignii* Boutique | - | - | KX998997 | KX999031 | - | KX999070 | - | - |
| 886 | Xylopia_frutescens_var_ferruginea | *Xylopia frutescen* var. *ferruginea* R. E. Fr. | - | KX998976 | KX998998 | KX999032 | - | KX999071 | - | - |
| 887 | Xylopia_frutescens_var_frutescens | *Xylopia frutescens* Aubl. | - | KX998977 | KX998999 | KX999033 | - | KX999072 | - | - |
| 888 | Xylopia_gilbertii | *Xylopia gilbertii* Boutique | - | KM924856 | KM924896 | KM924994 | KM924924 | KM924957 | - | - |
| 889 | Xylopia_heterotricha | *Xylopia heterotricha* D. M. Johnson & N. A. Murray | - | KX998979 | KX999001 | - | - | - | - | - |
| 890 | Xylopia_holtzii | *Xylopia holtzii* Engl. | - | KM924857 | KM924897 | KM924995 | KM924925 | KM924958 | - | - |
| 891 | Xylopia_humblotiana | *Xylopia humblotiana* Baill. | - | - | KX999002 | KX999035 | - | - | - | - |
| 892 | Xylopia_hypolampa | *Xylopia hypolampra* Mildbr. & Diels | - | KM924858 | KM924898 | KM924996 | KM924926 | KM924959 | - | - |
| 893 | Xylopia_hypolampra | *Xylopia hypolampra* Mildbr. | - | KX998980 | KX999003 | KX999036 | - | KX999074 | - | - |
| 894 | Xylopia_kalabenonensis | *Xylopia kalabenonensis* D. M. Johnson, Deroin, & Callmander | - | KX998981 | KX999004 | KX999037 | - | KX999075 | - | - |
| 895 | Xylopia_lamarckii | *Xylopia lamarckii* Baill. | - | - | - | KX999038 | - | - | - | - |
| 896 | Xylopia_lastelliana | *Xylopia lastelliana* Baill. | - | - | - | KX999039 | - | KX999076 | - | - |
| 897 | Xylopia_lemurica | *Xylopia lemurica* Diels | - | - | KX999005 | KX999040 | - | KX999077 | - | - |
| 898 | Xylopia_letestui | *Xylopia letestui* Pellegr. | - | KX998982 | KX999006 | KX999041 | - | KX999078 | - | - |
| 899 | Xylopia_longipetala | *Xylopia longipetala* De Wild. & T. Durand | - | - | KX999007 | KX999042 | - | KX999079 | - | - |

| No. | Voucher information | | GenBank accession numbers | | | | | | | |
|---|---|---|---|---|---|---|---|---|---|---|
| | Tip name in the tree | Taxa name | *atpB-rbcL* | *matK* | *ndhF* | *psbA-trnH* | *rbcL* | *trnL-F* | *trnS-G* | *ycf1* |
| 900 | Xylopia_lukei | *Xylopia lukei* D. M. Johnson & Goyder | - | - | KX999008 | KX999043 | - | KX999080 | - | - |
| 901 | Xylopia_maccreae | *Xylopia maccreae* (F.Muell.) L.S. Sm. | - | KM924860 | KM924900 | KM924998 | KM924928 | KM924961 | - | - |
| 902 | Xylopia_madagascariensis | *Xylopia madagascariensis* Cavaco & Keraudren | - | - | - | KX999044 | - | - | - | - |
| 903 | Xylopia_malayana | *Xylopia malayana* Hook. f. & Thomson | - | KX998983 | KX999009 | KX999045 | - | KX999081 | - | - |
| 904 | Xylopia_mildbraedii | *Xylopia mildbraedii* Diels | - | - | KX999010 | KX999046 | - | KX999082 | - | - |
| 905 | Xylopia_monosperma | *Xylopia monosperma* Jessup | - | KM924861 | KM924901 | KM924999 | KM924929 | KM924962 | - | - |
| 906 | Xylopia_mwasumbii | *Xylopia mwasumbii* D. M. Johnson | - | KP052715 | KP052723 | KP052731 | - | KP052739 | - | - |
| 907 | Xylopia_nitida | *Xylopia nitida* Dunal | - | JQ626567 | - | GQ428667 | JQ626284 | FJ039295 | - | - |
| 908 | Xylopia_pallescens | *Xylopia pallescens* Baill. | - | - | - | KX999048 | - | KX999084 | - | - |
| 909 | Xylopia_parviflora | *Xylopia parviflora* Spruce | - | - | KX999012 | KX999049 | - | KX999085 | - | - |
| 910 | Xylopia_perrieri | *Xylopia perrieri* Hance | - | - | - | KX999050 | - | KX999086 | - | - |
| 911 | Xylopia_pierrei | *Xylopia pierrei* Hance | - | KP052716 | KP052724 | KP052732 | - | KP052739 | - | - |
| 912 | Xylopia_polyantha | *Xylopia polyantha* R. E. Fr. | - | - | KX999014 | KX999051 | - | KX999087 | - | - |
| 913 | Xylopia_pseudolemurica | *Xylopia pseudolemurica* Cavaco & Keraudren | - | - | KX999015 | KX999052 | - | KX999088 | - | - |
| 914 | Xylopia_pulchella | *Xylopia pulchella* Ridl. | - | KP052717 | KP052725 | KP052733 | - | KP052741 | - | - |
| 915 | Xylopia_quintasii | *Xylopia quintasii* Engl. & Diels | - | KP052718 | KP052726 | KP052734 | - | KP052742 | - | - |
| 916 | Xylopia_staudtii_3233 | *Xylopia staudtii Engl. & Diels; WAG3223* | - | KM924862 | KM924902 | KM925000 | KM924930 | KM924963 | - | - |

| No. | Voucher information | | GenBank accession numbers | | | | | | | |
|---|---|---|---|---|---|---|---|---|---|---|
| | Tip name in the tree | Taxa name | *atpB-rbcL* | *matK* | *ndhF* | *psbA-trnH* | *rbcL* | *trnL-F* | *trnS-G* | *ycf1* |
| 917 | Xylopia_vielana | *Xylopia vielana* Pierre | - | KM924863 | KM924903 | KM925001 | KM924931 | KM924964 | - | - |
| 918 | Xylopia_aff_africana | *Xylopia* sp. aff. *africana* (Benth.) Oliv. | - | KX998986 | KX999016 | KX999053 | - | KX999089 | - | - |
| 919 | Xylopia_aff_buxifolia | *Xylopia* sp. aff. *buxifolia* Baill. | - | KX998987 | KX999017 | KX999054 | - | KX999090 | - | - |
| 920 | Xylopia_aff_fananehanensis | *Xylopia* sp. aff. *fananehanensis* Cavaco & Keraudren | - | KX998988 | KX999018 | KX999055 | - | KX999091 | - | - |
| 921 | Xylopia_aff_lastelliana | *Xylopia* sp. aff. *lastelliana* Baill. | - | - | - | KX999059 | - | KX999095 | - | - |
| 922 | Xylopia_aff_perrieri | *Xylopia* sp. aff. *perrieri* Diels | - | - | KX999021 | KX999060 | - | KX999096 | - | - |
| 923 | Xylopia_aff_sahafariensis | *Xylopia* sp aff. *sahafariensis* Cavaco & Keraudren | - | - | KX999022 | KX999061 | - | KX999097 | - | - |

**Supplementary Appendix 2.** Priors for BAMM analyses.

```
###############################################
# Prior block chosen by BAMMtools::setBAMMpriors

expectedNumberOfShifts = 1.0

lambdaInitPrior = 3.11328036689519

lambdaShiftPrior = 0.0122566467800711

muInitPrior = 3.11328036689519

#### End Prior block
######################
```
